# Plant genome evolution in the genus *Eucalyptus* driven by structural rearrangements that promote sequence divergence

**DOI:** 10.1101/2023.04.19.537464

**Authors:** Scott ferguson, Ashley Jones, Kevin Murray, Rose Andrew, Benjamin Schwessinger, Justin Borevitz

## Abstract

Genomes have a highly organised architecture (non-random organisation of functional and non-functional genetic elements within chromosomes) that is essential for many biological functions, particularly, gene expression and reproduction. Despite the need to conserve genome architecture, a surprisingly high level of structural variation has been observed within species. As species separate and diverge, genome architecture also diverges, becoming increasingly poorly conserved as divergence time increases. However, within plant genomes, the processes of genome architecture divergence are not well described. Here we use long-read sequencing and *de novo* assembly of 33 phylogenetically diverse, wild and naturally evolving *Eucalyptus* species, covering 1-50 million years of diverging genome evolution to measure genome architectural conservation and describe architectural divergence. The investigation of these genomes revealed that following lineage divergence genome architecture is highly fragmented by rearrangements. As genomes continue to diverge, the accumulation of mutations and subsequent divergence beyond recognition of rearrangements becomes the primary driver of genome divergence. The loss of syntenic regions also contribute to genome divergence, but at a slower pace than rearrangements. We hypothesise that duplications and translocations are potentially the greatest contributors to *Eucalyptus* genome divergence.

## Introduction

Genomes from all kingdoms are highly organised, but vary greatly in their structural architecture (Koonin, 2009). Within eukaryotic genomes, genome architecture refers to the non-random organisation of functional and non-functional genetic elements within chromosomes (genes, regulatory regions, small RNAs, transposons, pseudogenes, introns, centromeres, telomeres, etc.), and is critical for many biological functions, in particular reproduction and gene expression. However, the conservation and divergence of genome architecture or structure among a group of radiating plant species that share a common karyotype has not been well described.

For effective recombination during meiosis and the production of viable reproducing offspring, the genome architecture of both parental haplotypes must be highly similar. Changes to the genetic architecture can result in reproductive isolation/incompatibility or non-viable gametes (Hardigan *et al*., 2020; Simakov *et al*., 2020). Therefore, a common genome architecture within individuals of a breeding population tends to be highly conserved, except at some loci with high diversity (Jiao and Schneeberger, 2020). Similarly, for expression of a gene to be correctly regulated, it must be placed on a chromosome alongside the required promoters, enhancers, and inhibitors. The 3D organisation of the surrounding chromatin must permit physical access to allow transcription (Heng *et al*., 2004; Dixon, Gorkin and Ren, 2016; Oudelaar and Higgs, 2021).

Despite this functional need for structural conservation, some structural differences are known to exist between genomes within species. The extent to which reproductively compatible genomes are structurally different is an open area of research; however, several studies have shown genomes with a surprising amount of structural differences to be reproductively compatible (Lin and Gokcumen, 2019; Alonge *et al*., 2020; Jiao and Schneeberger, 2020; Tang *et al*., 2022). Between diverged species, genomes share less of their architecture than genomes within species, but typically genome architecture is conserved in proportion to phylogenetic distance (Luo *et al*., 2020; Weissensteiner *et al*., 2020; Derežanin *et al*., 2022; Ruggieri *et al*., 2022), and becomes poorly conserved at larger evolutionary distances (Koonin, 2009).

However, genomes have often, but not always, been viewed as containers to hold genes (Heng, 2009; Marques, Meier and Seehausen, 2019). The legacy of a gene-centric genome has persisted due to the Modern Synthesis (Crkvenjakov and Heng, 2022) and the highly influential work of Dawkins (Dawkins, 1976) and others. Guided by an evolutionary view dominated by genes and gene variants, many genomes from various species have been sequenced, and by identifying their genes and gene variants have provided us with a better understanding of the processes of evolution, divergence, and speciation (Rellstab *et al*., 2015; Schumer *et al*., 2015; Meier *et al*., 2017). However, a heavily gene-centric view may also limit our understanding (Heng, 2009). This heavily gene variant-based view of evolution was common until recent advances in long-read sequencing technologies enabled genome-wide investigations into genome architecture (Amarasinghe *et al*., 2020). Larger structural genome changes were thought to be rare, and as such genomes have been treated as largely structurally static, with individuals typically conceptualised as differing mostly by single nucleotide polymorphisms (SNPs) (Feulner and De-Kayne, 2017). Pan-genome studies by using a collection of genes or sequences in a population or species (Bayer *et al*., 2020; Lei *et al*., 2021), have revealed a significant amount of structural variation within genomes (Torkamaneh, Lemay and Belzile, 2021; Tang *et al*., 2022; Li *et al*., 2023).

Shared genome architecture is measured by synteny. Synteny is the conservation of both the order and sequence of homologous chromosomes between genomes (Passarge, Horsthemke and Farber, 1999; Dawson *et al*., 2007; Heger and Ponting, 2007). Synteny can refer both to individual genome regions, or in aggregate when comparing whole genomes. A genome pair with a large proportion of syntenic loci can be said to be more syntenic than a genome pair with a small proportion of syntenic loci. Synteny can become disrupted by the loss, gain, duplication, rearrangement, or divergence of existing sequences.

Rearrangements can occur as inversions, translocations, and duplications; altering the order of sequences within chromosomes while maintaining gene content and are often labelled as structural variants (SVs) (Rieseberg, 2001). Species-specific sequences resulting from the insertion, deletion, or localised divergence of sequence, appear as unaligned regions when genomes are analysed. The true origin of unaligned regions is more difficult to infer than rearrangements or syntenic regions (Weisman, Murray and Eddy, 2020).

Crucial to the study of plant genome evolution is study group choice. The ideal study group would be naturally evolving, have low prezygotic reproductive barriers, highly specious, and exist over a wide and variable evolutionary range. *Eucalyptus* with over 800 wild and undomesticated species that exist across a wide geographic and environmental range (Potts and Wiltshire, 1997; Booth *et al*., 2015; Supple *et al*., 2018), retain a conserved karyotype (Grattapaglia *et al*., 2015; Butler *et al*., 2017), are pollinated by generalist pollinators (Pfeilsticker *et al*., 2023), are capable of wide-ranging dispersal of genetic material (Bezemer *et al*., 2016; Murray *et al*., 2019), and span 50 million years of divergent evolution (Thornhill *et al*., 2019) make an ideal genus to study plant genome evolution.

Continuing our study into plant genome evolution (Ferguson *et al*., 2022), we generated long read sequences and assembled the genomes of 30 undomesticated *Eucalyptus* genomes and outgroups from two closely related genera, *Angophora floribunda* and *Corymbia maculata*. Combined with three previously and identically assembled *Eucalyptus* genomes (Ferguson *et al*., 2022), we create a dataset covering approximately 1-50 million years of diverging genome evolution, including all eight *Eucalyptus* subgenera (Thornhill *et al*., 2019; Nicolle, 2022). Identifying all syntenic and rearranged regions between all species pairs we demonstrate the rapid pace at which ancestral genome architecture is lost. We further analysed our results to determine if ancestral genome architecture was being lost to sequence rearrangement, divergence beyond recognition, or insertions and deletions. Additionally, by framing synteny, rearrangement, and unaligned loss or gain with phylogenetic distance we sought to describe the overall pattern of genome evolution.

## Results

### Sequencing and Assembly

To investigate genome architecture, we performed nanopore long-read native DNA sequencing and *de novo* genome assembly for 32 Eucalypt species (30 *Eucalyptus*, one *Angophora*, and one *Corymbia*; Table 1). All read libraries were trimmed and filtered in preparation of assembly. Curated read libraries had an average haploid coverage of 42.8x (range: 24.7x to 78.0x). For details of read libraries and sequence length distributions, see Supplementary Table S1 and S2, and Supplementary Figures S1 and S2. *Eucalyptus pauciflora* fast5 files were obtained from Wang et al (Wang et al., 2020), processed and assembled as per our datasets, using a randomly selected 60x coverage of reads.

**Table 1.** Summary of de novo genome assembly, quality assessment, and annotation of 33 Eucalypt genomes. Alphabetically ordered list of genomes assembled, listing: scaffolded genome size, proportion of contigs that were placed in a scaffold, scaffold N50 score, contig N50 score, the number of contigs in unscaffolded genomes, BUSCO scores for scaffolded genomes, LAI score, and the proportion of genome occupied TEs. *Eucalyptus albens*, *E. melliodora*, and *E. sideroxylon*, have been previously reported, being assembled using the same pipeline (Ferguson *et al*., 2022).

| Species | Scaffolded genome size (Mbp) | % of genome in scaffolds | Scaffold N50 (Mbp) | Contig N50 (Mbp) | Contig count | BUSCO complete | LAI | TE % |
| --- | --- | --- | --- | --- | --- | --- | --- | --- |
| *A. floribunda* | 388.21 | 99.73% | 36.02 | 4.02 | 224 | 96.82% | 14.5 | 34.55% |
| *C. maculata* | 403.82 | 99.90% | 40.55 | 4.69 | 173 | 97.25% | 15.92 | 36.26% |
| * *E. albens* | 606.89 | 99.79% | 56.93 | 2.55 | 674 | 96.47% | 17.3 | 46.57% |
| *E. ANBG9806169* | 507.93 | 99.65% | 49.61 | 2.40 | 476 | 96.86% | 22.16 | 44.00% |
| *E. brandiana* | 507.08 | 99.82% | 45.47 | 7.28 | 168 | 98.11% | 23.85 | 44.21% |
| *E. caleyi* | 589.32 | 99.53% | 59.52 | 4.77 | 276 | 96.47% | 18.24 | 46.00% |
| *E. camaldulensis* | 558.45 | 99.87% | 52.65 | 2.48 | 418 | 96.73% | 16.99 | 45.31% |
| *E. cladocalyx* | 544.08 | 99.68% | 51.92 | 2.80 | 390 | 97.59% | 18.53 | 45.85% |
| *E. cloeziana* | 480.07 | 99.75% | 44.75 | 1.74 | 625 | 97.12% | 19.06 | 42.57% |
| *E. coolabah* | 606.31 | 99.53% | 53.56 | 1.29 | 935 | 95.44% | 15.9 | 45.89% |
| *E. curtisii* | 435.26 | 99.96% | 40.29 | 2.96 | 288 | 97.29% | 18.34 | 41.66% |
| *E. dawsonii* | 706.90 | 99.35% | 67.73 | 0.99 | 1,342 | 97.51% | 17.01 | 45.88% |
| *E. decipiens* | 590.95 | 99.50% | 60.20 | 1.99 | 552 | 96.99% | 18.87 | 46.95% |
| *E. erythrocorys* | 539.20 | 99.99% | 50.47 | 4.02 | 250 | 97.55% | 20.18 | 47.07% |
| *E. fibrosa* | 589.91 | 99.85% | 55.66 | 6.45 | 192 | 96.73% | 17.49 | 45.10% |
| *E. globulus* | 545.02 | 99.28% | 51.39 | 0.64 | 1,747 | 96.69% | 17.46 | 44.29% |
| *E. grandis* | 615.89 | 99.44% | 58.49 | 0.61 | 1,747 | 96.09% | 17.11 | 46.53% |
| *E. guilfoylei* | 472.36 | 99.97% | 44.61 | 4.25 | 209 | 98.02% | 16.39 | 41.22% |
| *E. lansdowneana* | 633.52 | 99.92% | 59.67 | 2.35 | 489 | 97.12% | 19.46 | 46.10% |
| *E. leucophloia* | 568.48 | 99.38% | 54.41 | 2.66 | 382 | 96.99% | 17.91 | 44.37% |
| *E. marginata* | 512.89 | 98.83% | 50.56 | 1.01 | 989 | 96.17% | 19.58 | 43.43% |
| * *E. melliodora* | 639.15 | 99.30% | 60.83 | 1.87 | 564 | 98.67% | 18.32 | 47.20% |
| *E. melliodora* x *E. sideroxylon* | 603.57 | 99.80% | 57.05 | 6.22 | 281 | 97.72% | 17.96 | 46.71% |
| *E. microcorys* | 440.91 | 99.92% | 41.20 | 4.00 | 233 | 97.21% | 16.2 | 41.39% |
| *E. paniculata* | 588.85 | 99.66% | 55.38 | 3.70 | 330 | 97.12% | 18.58 | 44.92% |
| *E. pauciflora* | 494.03 | 99.88% | 50.46 | 6.58 | 209 | 97.25% | 20.29 | 43.10% |
| *E. polyanthemos* | 603.28 | 99.56% | 57.46 | 4.66 | 300 | 96.82% | 17.52 | 45.55% |
| *E. pumila* | 529.75 | 99.70% | 48.19 | 2.49 | 473 | 97.38% | 17.74 | 44.17% |
| *E. regnans* | 494.97 | 99.84% | 47.06 | 5.26 | 205 | 97.25% | 20.18 | 43.06% |
| *E. shirleyi* | 597.18 | 99.88% | 56.34 | 6.91 | 181 | 97.29% | 19.89 | 45.85% |
| * *E. sideroxylon* | 592.133 | 99.87% | 62.13 | 5.22 | 297 | 96.65% | 18.68 | 46.57% |
| *E. tenuipes* | 397.78 | 99.99% | 35.74 | 3.43 | 207 | 96.39% | 15.07 | 37.82% |
| ***E. victrix*** | 557.16 | 99.85% | 53.19 | 11.10 | 120 | 96.65% | 18.71 | 44.34% |
| ***E. viminalis*** | 558.71 | 99.11% | 52.93 | 0.65 | 1,755 | 96.47% | 16.5 | 44.57% |
| ***E. virginea*** | 532.79 | 99.97% | 56.15 | 2.39 | 376 | 97.08% | 17.69 | 43.78% |

After assembling our trimmed and filtered read libraries, we curated our genomes, removing contigs identified as contamination and assembly artefacts. Additionally we filtered haplotigs from our primary assemblies to form pseudo-haploid genomes (Supplementary Table S3).

Our genomes, which have a known and conserved haploid karyotype of 11 chromosomes (Ribeiro et al., 2016), assembled into an average of 517 contigs (range: 120 to 1,755) (Table 1). At the completion of our assembly pipeline our genomes had an average contig N50 of 3.65 Mbp (range: 614.30 Kbp to 11.10 Mbp). Scaffolding contigs against *E. grandis* (Myburg et al., 2014) greatly increased our genome contiguity, placing on average 99.69% of our genomes into pseudo-chromosomes (range: 98.83% to 99.99%). On average each pseudo-chromosomes was comprised 46 contigs. We have found syntenic scaffolding within *Eucalyptus* to be suitable in the absences of chromosome conformation data (Ferguson *et al*., 2022) as Eucalypts have a conserved karyotype (Healey *et al*., 2021; Low *et al*., 2022). Additionally, within other genera closely related genomes have been found suitable for scaffolding (Burns *et al*., 2021). Additionally, RaGOO provides confidence scores for assigning contigs to a scaffold, ordering contigs within scaffolds, and orienting contigs within scaffolds. Confidence scores achieved by our genomes indicated scaffolding was satisfactory, Supplementary Figure S3.

The completeness of our genomes was evaluated with benchmarking universal single-copy orthologs (BUSCO) (Manni *et al*., 2021) and LTR Assembly Index (LAI) (Ou, Chen and Jiang, 2018). A more complete genome will contain a high proportion of single-copy BUSCO genes, and all our genomes were found to be highly BUSCO complete (average: 97.01%; range: 95.44% to 98.11). LAI searches a genome for long terminal repeat (LTR) sequences, and reports on the proportion that are intact. The LAI scores achieved by our genomes indicate that they are highly complete (average: 18.17; range: 14.50 to 23.85). Quality scores for all our genomes indicate that our genomes are of high quality, contiguity and completeness (Table 1 and Supplementary Table S4). For statistics and sequence distribution plots describing our genomes during and at the completion of assembly (see Supplementary Tables S5 and S6, and Supplementary Figures S4 to S6).

### Genome annotation

As masking of repeats within genomes aids in gene annotation, we annotated our genomes for both transposable elements (TEs) and simple repeats. Repeat annotation was performed using *de novo* repeat libraries built for each genome. Repeat annotation resulted in the classification of an average of 43.78% (range: 34.55% to 47.07%) of our genomes as TEs, and an average of 1.25% (range: 1.14% to 1.39%) as simple repeats, Table 1 and Supplementary Table S7. After soft-masking all genomes, we trained species-specific gene HMM models and subsequently annotated all genomes for genes. Gene models were trained on all available gene transcripts for *A. thaliana* (Taxonomy ID: 3702) and Myrtaceae (Taxonomy ID: 3931) found within the NCBI (Sayers *et al*., 2021). Annotation predicted an average of 53,390 (range: 41,623 to 77,764) gene candidates within our genomes, Supplementary Table S8. While the number of annotated genes is consistent with plant gene number estimates (Sterck *et al*., 2007) there is a wide variation between genomes. It is important to note the genes annotated within these genomes will contain both false positives and false negatives and are gene candidates, which in addition to real gene number variations will contribute to the variation in the number of annotated genes.

### *Eucalyptus* pan-genome

Due to the shared evolutionary history of our genomes, many gene candidates will be homologues that have arisen pre-speciation (orthologs) or post-speciation (paralogs) (Jensen, 2001). To examine the evolutionary relationship between *Eucalyptus* gene candidates we placed all highly similar primary (longest) gene transcripts into orthogroups (OG). Of the 1,761,851 identified gene candidates across our 33 *Eucalyptus* genomes, 1,726,511 (97.99%) were placed into one of 68,248 orthogroups (OG). The remaining 35,340 (2.01%) unique genes were not placed within an OG as their sequences were too dissimilar (>40% transcript identity and e-value < 0.001) to all other genes. On average each genome had 98.03% (range: 94.62% to 98.03%) of its gene candidates placed within an OG. 0.26% (4,551) of all gene candidates were found to occur within a genome-specific OG. For detailed statistics on orthogrouping see Supplementary Tables S8 and S9. Additionally, OGs were classified as core (present in all species), dispensable (present in at least two species), and private (present in a single species) (Figure 1). A total of 21.33% (14,552) of OG were core, likely representing key *Eucalyptus* genes. Most OGs were dispensable, 76.00% (51,858), which may be a source of phenotypic and adaptive variation within the species. Only a very small number were private, 2.67% (1,821), potentially representing highly species-specific genes and newly evolved genes.

**Figure 1.**
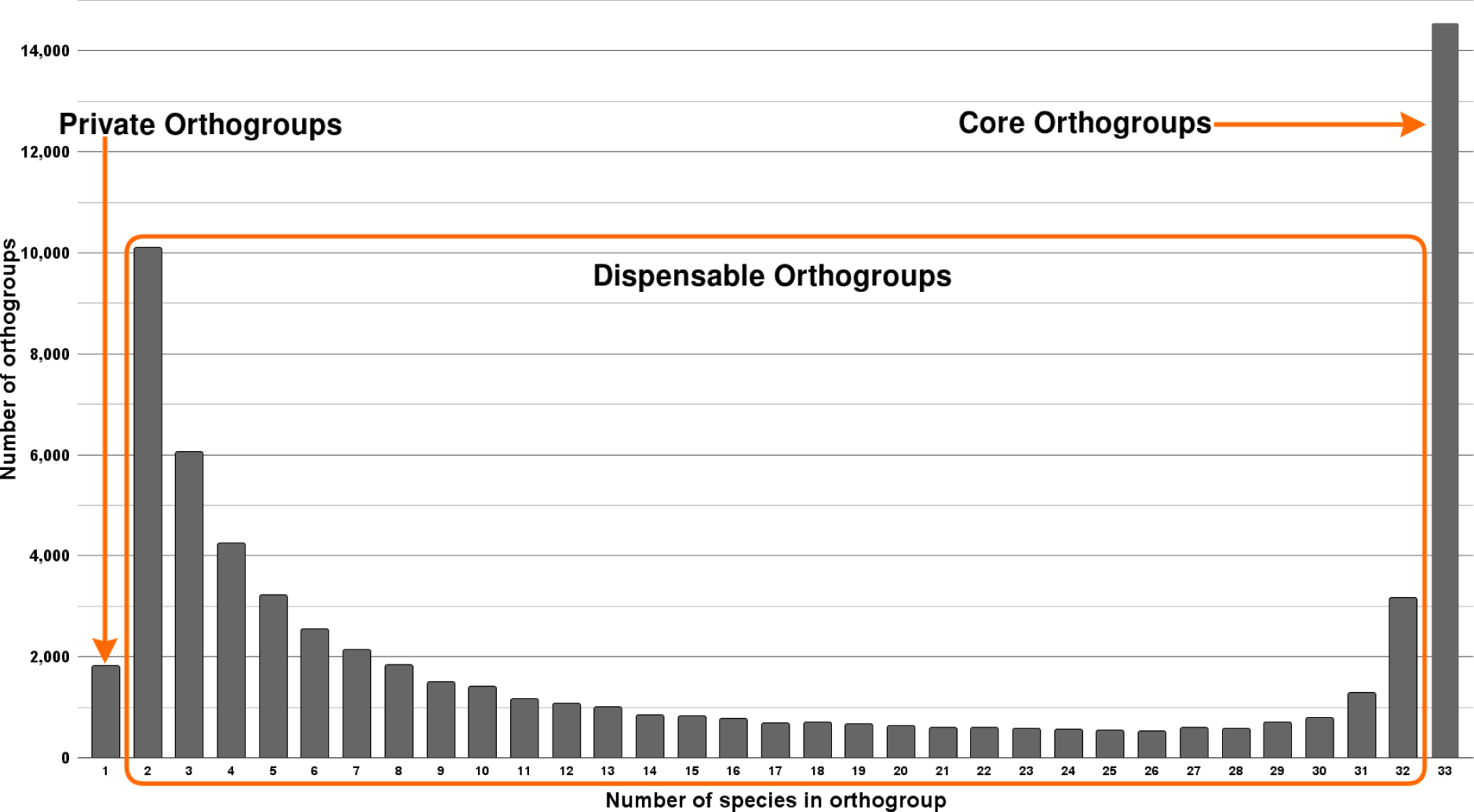
Pangenome of 33 species of *Eucalyptus*. Shows the number of orthogroups shared by an increasing number of genomes. Private orthogroup are orthogroups that exist within a single genome, core exist in all, and dispensable orthogroups are those that exist in 2 to 32 (n-1) genomes.

### *Eucalyptus* phylogeny

To describe the evolutionary patterns between our genomes we built a phylogenetic tree from single-copy BUSCO genes. We additionally included the genome of *C. calophylla* genome, which was identically assembled (Ahrens *et al*., 2021). From the initial BUSCO set of 2,326 genes, we selected only genes present within 30 or more genomes, leaving 2,106 BUSCO genes across our 36 genomes or 72,516 total genes. For each gene, we generated a multi-sequence alignment (MSA) with MAFFT, which we then trimmed and filtered, removing low abundance regions and genes with overall poor alignments, leaving 1,674 gene MSAs. Each MSA was used to construct a gene tree, subsequently all gene trees were combined into a consensus species tree. The species tree was manually rooted using the established relationship between *Angophora*, *Corymbia* and *Eucalyptus* (Thornhill *et al*., 2019), Figure 2 and Supplementary Figure S7. The species tree in Newick format is available within Supplementary results.

**Figure 2.**
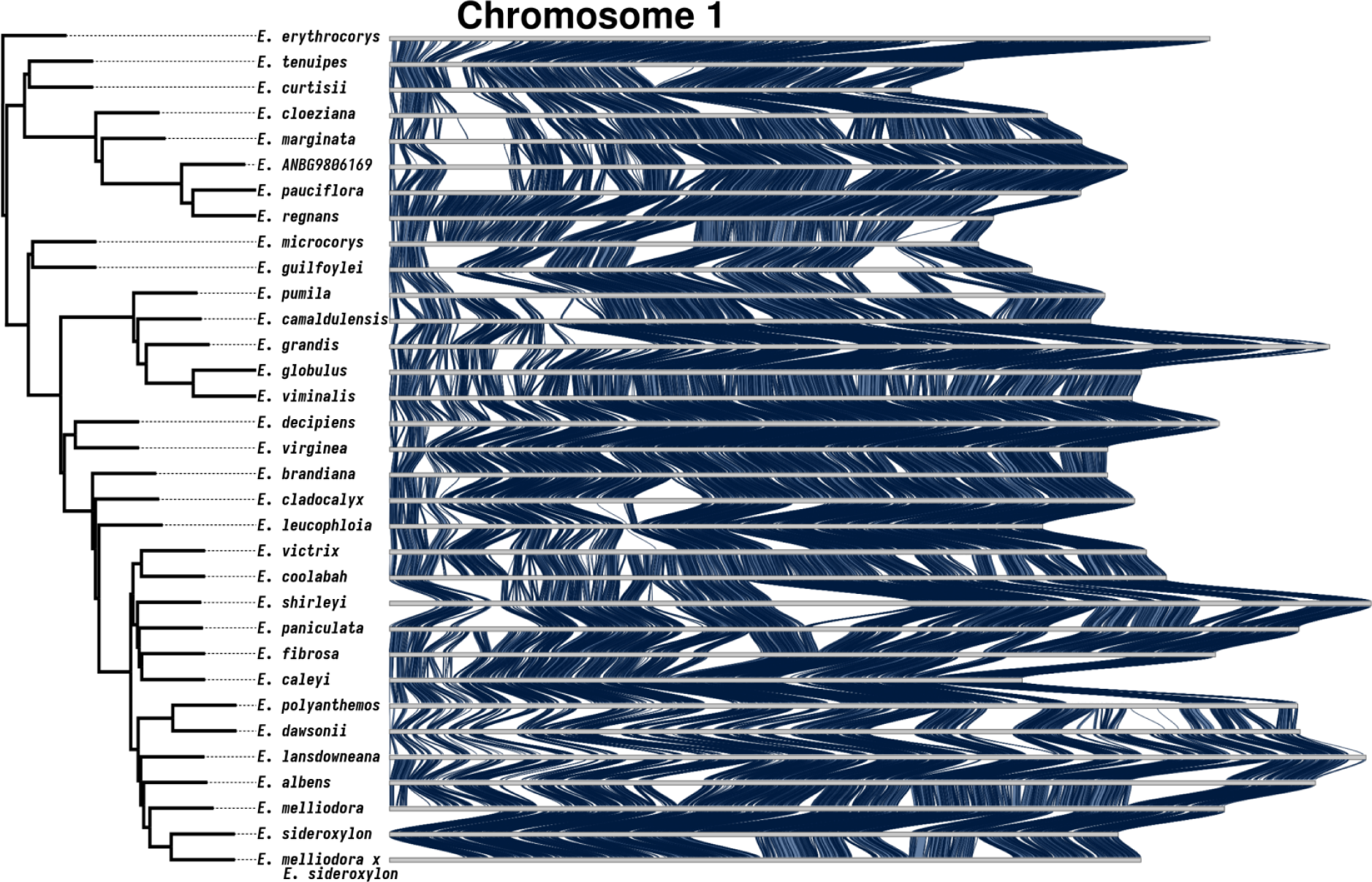
Synteny karyotype of chromosome 1. Blue ribbons between karyotypes indicate the presence of syntenic sequences between species pairs. In all other regions synteny has become lost. Synteny is lost to either rearrangements (inverted, translocated, or duplicated), sequence divergence, loss or gain. Chromosomes are ordered by our phylogenetic tree.

After constructing the species tree, *E. salubris* was unexpectedly found to be grouped with *E. pauciflora* and *E. regnans*. If correctly placed, *E. salubris* would be a sister lineage to the *Adnataria* group (*E. victrix* to *E. sideroxylon*) (Thornhill *et al*., 2019). Morphological examination of the sample tree revealed that the tree was incorrectly labelled. The correct species name is currently unknown, as such we use its NCBI name *E. ANBG9806169*.

### Genome conservation and loss

To resolve the syntenic and non-syntenic regions of our *Eucalyptus* genomes, we performed one-to-one genome comparisons for all genome pairs. Whole genome alignments for all comparisons were analysed with SyRI (Goel *et al*., 2019), and subsequently, all genomic regions within both genomes of an alignment pair were annotated as syntenic, rearranged (inversion, translocation, or duplication), or unaligned (sequence that only exists in one genome, resulting from either an insertion, deletion or sequence divergence). As repeat masking would inflate the unaligned proportion of genome alignments and bias results, all genomes remained unmasked. This analysis resulted in all genomes being annotated for syntenic, rearranged, and unaligned regions 32 times, giving a total of 1,056 annotated genomes. A visual summary of shared synteny was plotted using our phylogenetic ordering, Figure 2 and Supplementary Figures S8 to S17. Inspection of synteny plots indicated that syntenic regions exist across the length of all chromosomes, however, synteny has become highly fragmented, and is differently maintained.

Next, we calculated the proportion of sequences shared between genomes, how sequences were shared (syntenic, inverted, translocated, and duplicated), and the frequency at which rearrangements occurred between genomes. For this analysis, we excluded all events less than 200 bp in length (the majority of which were small unaligned annotations). The majority of sequence was shared (syntenic and rearranged) between genomes, averaging 69.35% (range 46.67% to 91.86%). Only four pairwise alignments had <50% shared sequence: *E. coolabah, E. dawsonii, E. grandis, and E. melliodora*, all when compared to *E. erythrocorys*. Synteny was the major contributor to shared sequence, averaging 39.32% (range: 21.34% to 60.44%). Rearrangements averaged: 30.24% (range: 16.97% to 49.49%). The remainder of sequence was annotated as unaligned, averaging 30.43% (range: 8.08% to 53.32%), Table 2. For a per-species comparison breakdown of the percent of genome shared, syntenic, rearranged, and unaligned see Supplementary Tables S10 to S13.

**Table 2.** Summary of synteny, rearranged, and unaligned statistics of all pairwise genome analyses. For each pairwise comparison the number and total size of syntenic, unaligned, inverted, translocated, duplicated, rearranged (sum of all inverted, translocated, duplicated regions), and total shared was calculated. Additionally, the proportion of the genome occupied by all event types was calculated. Shown here are the average and range of these statistics.

|  | Average event size<br>within each pairwise<br>alignment (Kbp) |  |  | Event counts |  |  | Percent of genome |  |  |
| --- | --- | --- | --- | --- | --- | --- | --- | --- | --- |
| Alignment<br>type | Least | Most | Average | Least | Most | Average | Least | Most | Average |
| Syntenic | 14.03 | 29.45 | 18.34 | 6,657 | 18,810 | 12,153 | 21.34% | 60.44% | 39.32% |
| Unaligned | 3.28 | 15.62 | 7.58 | 11,808 | 35,983 | 22,365 | 8.08% | 53.32% | 30.43% |
| Inverted | 29.48 | 477.33 | 125.18 | 81 | 209 | 148 | 0.87% | 10.46% | 3.28% |
| Translocated | 6.70 | 22.98 | 11.65 | 4,322 | 15,975 | 9,521 | 9.26% | 41.14% | 19.92% |
| Duplicated | 2.48 | 6.14 | 3.60 | 3,807 | 32,286 | 15,350 | 3.60% | 38.97% | 14.03% |
| Rearranged | - | - | - | 8,274 | 46,066 | 25,020 | 16.97% | 49.49% | 30.24% |
| Total shared | - | - | - | - | - | - | 46.67% | 91.86% | 69.35% |

Examination of the size and frequency of syntenic regions indicates that synteny between the 11 chromosomes of all genome pairs has, on average, fragmented into 12,153 (range: 6,657 to 18,810) regions with an average size of 17.97 kbp (range: 16.27 kbp to 24.26 kbp) (see Supplementary Figures S18 and S19 for per genome average event size and frequency plots). Rearrangements in total (inversions + duplications + translocations) contributed more to synteny loss than did unaligned regions, however, unaligned contributed more than any single rearrangement type. A more detailed examination of the relative size and frequency of syntenic, rearrangement, and unaligned events showed that syntenic regions were long and common, unaligned regions were short and common, inversions were long and very rare, duplications were shortest and very common, and translocations occurred at a moderate frequency and size. Syntenic regions are distributed over the entire length of all chromosomes between all genome pairs, however, synteny has become highly fragmented by rearrangements and unaligned regions.

### Divergence time and genome conservation/loss

To examine these trends of architecture change over increasing divergence time, we examined the relationship between phylogenetic distance and genome conservation and divergence, Figure 3. Not unexpectedly, we find that as phylogenetic distances increase, the proportion of syntenic (R^2^ = 0.261) and rearranged (R^2^ = 0.356) sequence decreased as lineages acquire unique genomic variation. Similarly, as phylogenetic distances increase, the proportion of genomes within duplications (R^2^ = 0.189) and translocations (R^2^ = 0.240) decreased. Whereas, the portion of genomes unaligned quickly increases with increasing phylogenetic distance (R^2^ = 0.536). Inversions consistently occupied a small proportion of genomes across all phylogenetic distances (R^2^ = 0.000).

**Figure 3.**
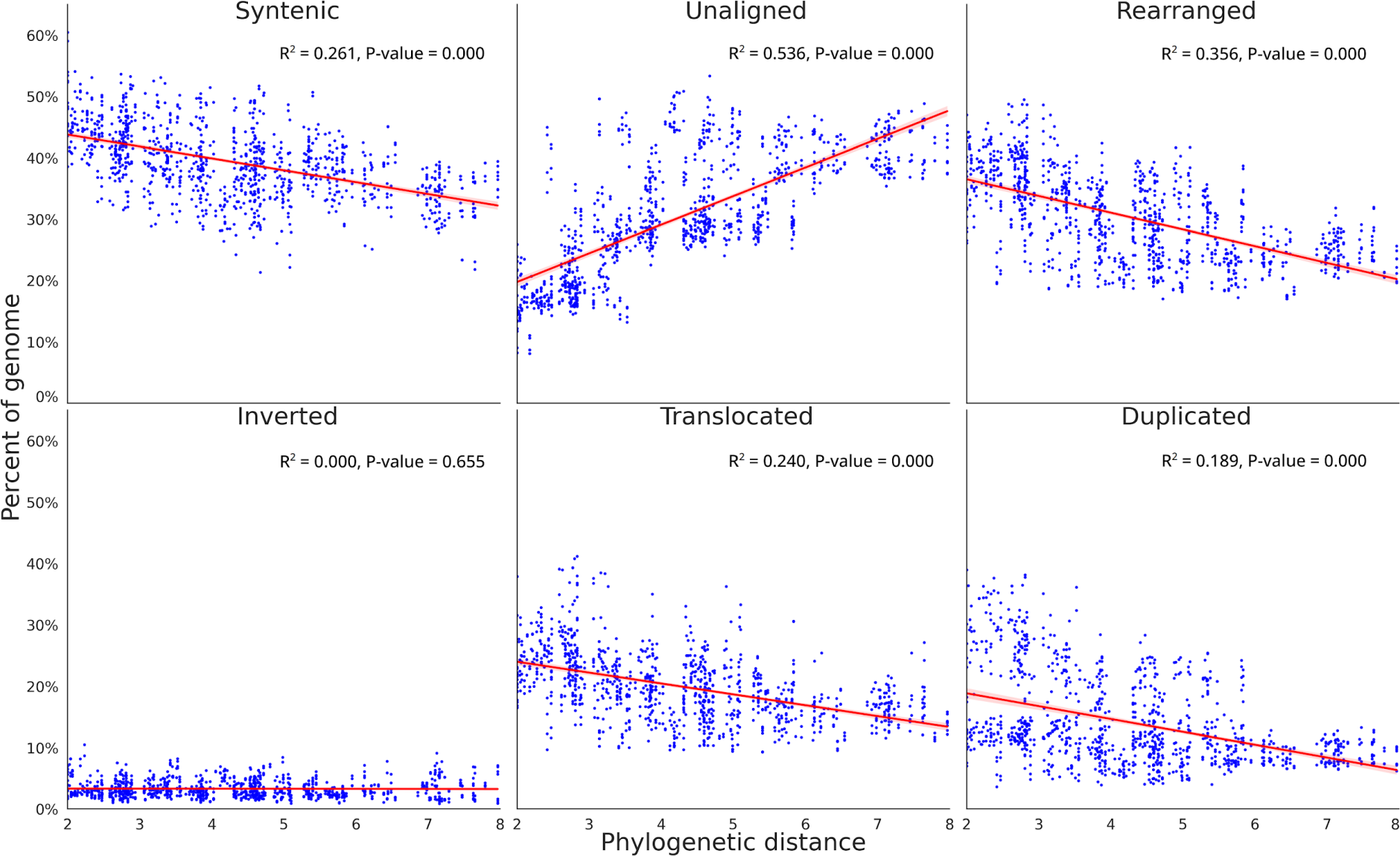
Pairwise genome conservation and loss, as phylogenetic distance increases. The proportion of both *Eucalyptus* genomes with an alignment pair that was identified as syntenic, rearranged, or unaligned, plotted against the phylogenetic distance of the two genomes. The unaligned proportion is the species-specific fraction of the genome between genome pairs, resulting from either an insertion, deletion, differential inheritance, or sequence divergence. When combined, the proportion of sequence that is syntenic, unaligned, and rearranged equals 100% for each genome within an alignment pair. The rearranged fraction is further broken down into inverted, translocated, and duplicated regions. Phylogenetic distance was calculated as the sum of branch lengths between each genome pair within phylogeny. P-value tests if the slope of the regression line is nonzero.

Unaligned sequences accumulate through the loss, gain, or divergence of sequences. As genome sizes are similar (average: 552.75 Mbp; standard deviation: 65.62 Mbp) sequence loss and gain are unlikely to fully explain the rapid accumulation of unaligned sequences. Divergence beyond recognition is likely the largest contributing factor. To test which regions were contributing to the growth of unaligned sequences we gathered all alignment identity scores for all syntenic, inverted, translocated, and duplicated regions, in each pairwise alignment. Plotting identities against phylogenetic distance, we examined the rate at which sequences diverge. Syntenic was observed to lose sequence homology more rapidly (R^2^: 0.516), than duplicated (R^2^: 0.236), translocated (R^2^: 0.303), and inverted (R^2^: 0.260), Supplementary Figure SX. However, in all cases the regression spanned a very small interval (syntenic: 91.58% - 93.14%, duplicated: 91.48% - 92.63%, translocated: 91.56% - 92.81%, and inverted: 91.49% - 92.72%) and none approached our 80% sequence similarity threshold for alignments.

Overall, we find that the syntenic proportion of the genome decreases slowly with increasing divergence time, while the proportion rearranged as duplications and translocations decrease faster. The loss of homology between synteny, duplicated and translocated regions leads to a strong increase in the unaligned portion of the genome (insertions, deletions, and diverged sequences) as divergence time increases. The loss of duplications and translocations contribute more to the growth of unaligned than does synteny.

### Genome-specific and group-wide sequences

Unaligned sequences occupied on average 30.65% of each *Eucalyptus* genome within each pairwise alignment. To determine if these sequences were unique to a single genome or shared between multiple, all pairwise alignments for each species were combined and the number of species sharing each base calculated. Subsequently, genome regions that were unique to a genome, shared by multiple genomes or shared by all genomes identified, Figure 4.

**Figure 4.**
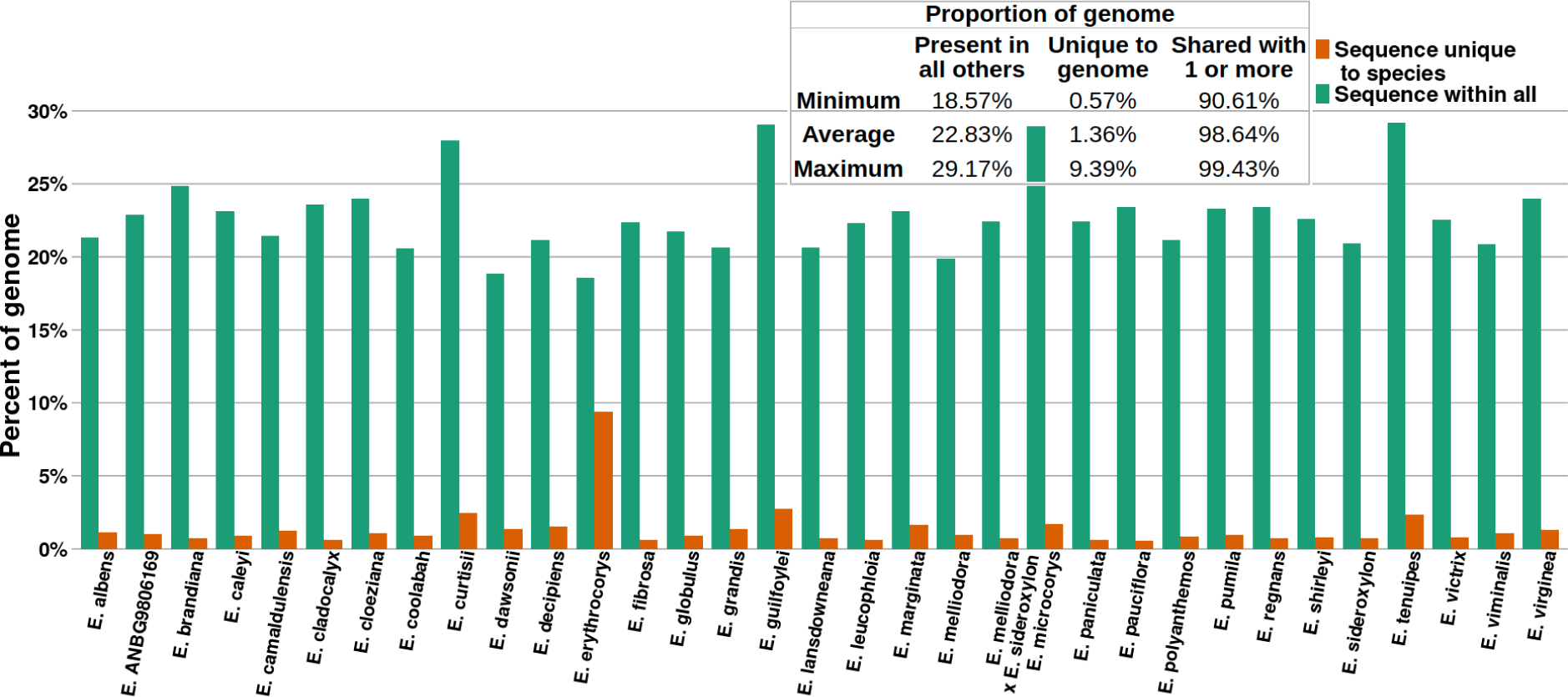
Proportion of *Eucalyptus* genomes unique and shared by all others. Sequence unique to species is the union of the genome that was classified as unaligned within all pairwise alignments. Sequence within all is the union of the genome that was classified syntenic and rearranged (i.e. common between genomes) within all pairwise alignments.

Genome-specific (unique) sequences occupied an average of 1.36% (241.55 Mbp) of the 33 *Eucalyptus* genomes, the remaining 98.64% of sequence was shared by one or more genomes. The proportion of each genome shared by all others averaged 22.83%. This finding mirrors our OG analysis where 2.67% of groups were private, 76.00% dispensable, and 21.33% were core.

Of note is *E. erythrocorys*, whose genome had a significantly lower proportion of genome-specific sequence and higher proportion of sequence shared by all other genomes. *Eucalyptus erythrocorys* is the sister taxon of all our other genomes within our *Eucalyptus* dataset. Given the age of the divergence between the *E. erythrocorys* lineage and its sister lineage, this genome was expected to display a unique pattern in this analysis, however, the extent to which *E. erythrocorys* is different from all others was surprising.

### Lineage-conserved rearrangements

The one-to-one analysis of our *Eucalyptus* genomes has described a genome structure that has become highly fragmented by frequently occurring rearrangements and unaligned regions. As genome structure is inherited by offspring, some of the rearrangements discovered during our analysis are assumed to exist within the genomes of monophyletic groups, i.e. a group of species that have descended from a single ancestral species. Rearrangements found within multiple genomes also help to confirm their validity. To search for evidence of inherited rearrangements we analysed the *Adnataria* section, for which we have the best coverage of genomes (13 genomes, as listed in Figure 5). Additionally, using only *Adnataria* genomes should maximise the occurrence of retained rearrangements, as the phylogenetic distances within the *Adnataria* group are relatively low, with many species still hybridising (Delaporte, Conran and Sedgley, 2001).

**Figure 5.**
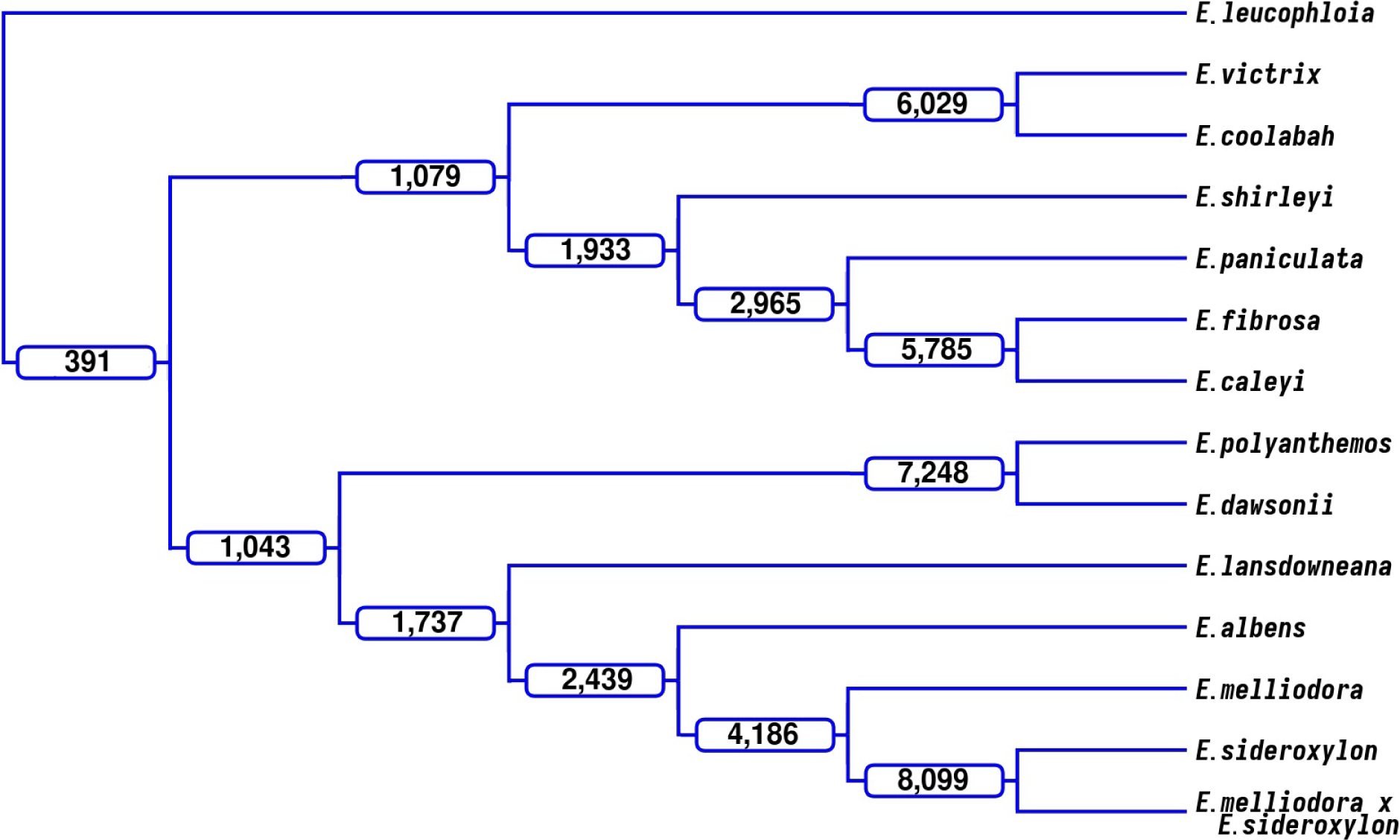
Lineage-conserved rearrangements. Using an outgroup genome, *E. leucophloia*, rearrangements were identified that were shared among members of a lineage, i.e. rearrangements with the same start and end points within the outgroup genome. At each branch in the dendrogram, the number of rearrangements shared by all taxa within that clade are labelled. e.g. *E. victrix* and *E. coolabah* share 6,029 rearrangements.

As all alignments and subsequent annotations are relative to the two species involved, directly comparing the breakpoints of annotations to find common rearrangements is not possible. Therefore, an outgroup genome, *E. leucophloia*, a sister species of the *Adnataria* group, is used for comparison with each of the 13 selected genomes. The outgroup genome imposes a single set of genetic coordinates and genome architecture, enabling comparisons of rearrangement breakpoints and subsequent identification of shared rearrangements. Shared rearrangements will contain the same sequence. While this method allows us to find common inversions, translocations, and duplications, it doesn’t allow us to find unaligned (insertions, deletions, and highly diverged) regions between our ingroup genomes, as genomes are not being directly compared.

Comparing the start and end breakpoints (± 50 bp) for events >1 Kbp (250,693 total rearrangements across all *Adnataria* genomes) identified 58,388 (23.29%) common rearrangements (rearrangements that exist within two or more genomes). Of the 58,388 common rearrangements, 28,059 (48.06%) were shared by two genomes, and 391 (0.67%) were shared by all. The number of common rearrangements quickly decreased as the number of genomes increased, (Supplementary Figure S20). Lineage-conserved rearrangements were identified by tracing common rearrangements through *Adnataria’s* phylogeny, Figure 5. As expected, more closely related genomes shared the largest number of rearrangements, while more distant genomes shared less. Additionally, as the number of descendant genomes of nodes increased, the number of shared rearrangements also decreased. Inherited rearrangements were identified within the *Adnataria* group. We repeated this analysis twice using *E. brandiana* and *E. cladocalyx* as the outgroup genome achieving similar results, Supplementary Figures S21 and S22.

### Gene content of synteny, rearrangements, and unaligned

To assess whether rearrangements that encompass genes are selected against, we calculated the proportion of genic (contains a gene/s) and non-genic (contains no gene/s) rearrangements, as well as syntenic and unaligned events per genome. Initially, all events too small to contain a gene and genes unplaced within an orthogroup were removed. A conservative event length of 1 kbp was used to filter out events, as events smaller than this are unlikely to contain a gene (Xu *et al*., 2006). Genes unplaced within an orthogroup are highly dissimilar to all other gene candidates and may be false positives resulting from incorrect annotation. The remaining rearrangement, synteny, and unaligned events were examined for the presence of genes placed within an OG and subsequently classed as genic or non-genic.

For each genome, when compared to all other genomes, we calculated the average proportion of genic syntenic, inverted, translocated, duplicated, and unaligned events, and plotted the results (Figure 6). An average of 88.80% (range: 82.52% to 95.57%) genic syntenic events were observed across our genomes. Genic unaligned averaged 41.13% (range: 19.76% to 73.48%), genic inversions averaged 94.93% (range: 81.65% to 99.13%), genic translocations averaged 65.70% (range: 48.77% to 83.98%), and genic duplications averaged 45.71% (range: 30.59% to 79.20%).

**Figure 6.**
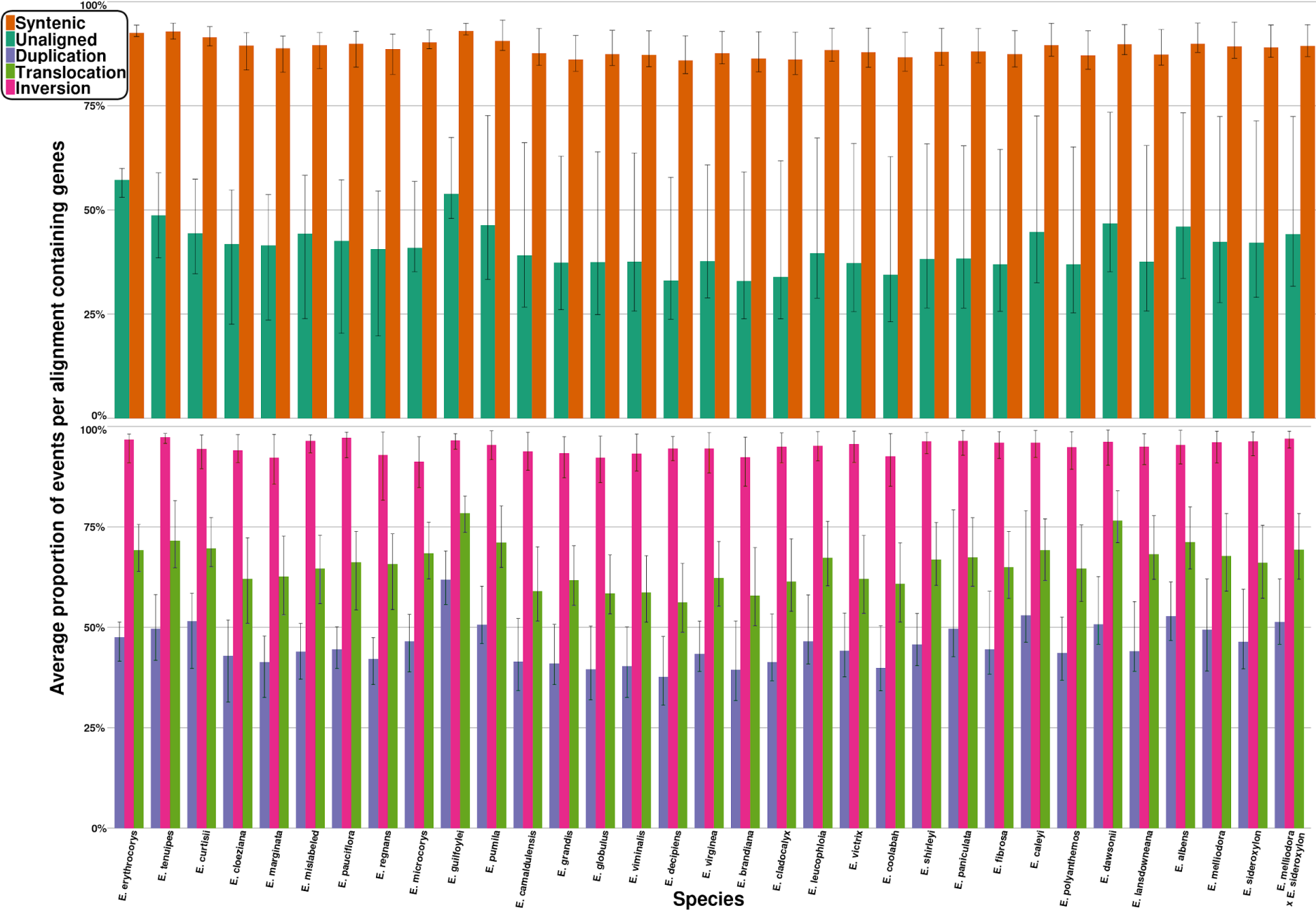
Average proportion of genic events for each species genome. The proportion of genic events was calculated for each pairwise alignment, and averaged. Error bars indicate the minimum and maximum proportion of genic events found when aligned to all other genomes.

Additionally, we analysed the effects of divergence time on the proportion of genic events for all rearrangement types, synteny, and unaligned, Supplementary Figures S23. Results indicated that phylogenetic distance has little to no impact on the proportion of genic syntenic events (R^2^ = 0.162), inverted events (R^2^ = 0.000), translocation events (R^2^ = 0.004), or duplication events (R^2^ = 0.002). Unaligned was the only event type whose genic proportion was affected by phylogenetic distance. As phylogenetic distance increases, unaligned events become more genic (R^2^ = 0.292; p-value = 0.000).

## Discussion

In this study we created a large collection of wild and naturally evolving high-quality *Eucalyptus* genomes covering 1-50 million years of divergent evolution. Using these genomes we find a pattern of genome evolution led by an initial rapid accumulation of rearrangements and subsequently a slow loss of both rearranged and syntenic sequences as lineage-specific mutations erode sequence homology. Rearrangements, likely due to their recombination effects and subsequent fixation/reduction of alleles (Faria and Navarro, 2010), are lost more rapidly than syntenic regions. Translocations and duplications were the major disruptors of synteny and were rapidly lost as divergence times increased. Inversions did not contribute substantially to the loss of synteny or the loss of rearrangements, instead occurring at consistently low rates across all divergence times. As genome sizes remain constant and little species-specific sequence exists across our dataset, loss of existing sequence or gain of new sequence provides an unlikely explanation for the growth of unaligned sequences as divergence times increase.

Duplications are a major contributor to functional and genome divergence (Lynch and Conery, 2000; Adams and Wendel, 2005) especially within plant lineages (Hanada *et al*., 2008; Van de Peer, Maere and Meyer, 2009). We found that duplications were highly abundant between all genomes, and that at smaller divergence times, duplications contributed strongly to genome divergence. As time since divergence increased, the contribution of duplications to genome divergence lessened, becoming overshadowed by unaligned portions of the genome. However, at all phylogenetic distances duplications were a major contributor to genome divergence (occupying on average 14.03% of genomes across an average of 15,350 events). The observed pattern of duplication loss as time since divergence increased was unsurprising, as duplications, while highly important to adaptation and evolution, are rarely conserved (Inoue *et al*., 2015; Naseeb *et al*., 2017). Why some duplications are preserved while the majority are lost is speculative; however, theory centres on neofunctionalisation, subfunctionalisation, and novel function evolution (Freeling, Scanlon and Fowler, 2015; Braasch *et al*., 2016; Lien *et al*., 2016; Wu and Cox, 2019). These hypotheses rely upon the genic properties of duplications, i.e. if duplications don’t gain novel function or retain ancestral function, purifying selection will likely result in their removal (Wu and Cox, 2019). While duplications were the least genic of all rearrangement types, a significant number (45.71%) were found to contain genes, likely contributing to their preservation. Non-genic duplications, being less visible to selection, are likely to experience increased evolutionary rates (mutations) and genetic drift (Scannell and Wolfe, 2008), eventually mutating beyond recognition, and ultimately contributing to the unaligned proportion of alignments. Duplications, both preserved and unpreserved, are likely one of the greatest sources of genome divergence.

Chromosomal inversions, which are known to be associated with the development of complex phenotypes, local adaptation, and speciation (Lowry and Willis, 2010; Twyford and Friedman, 2015; Arostegui *et al*., 2019), were extremely rare between all genomes (average 148 between genomes) and contributed less than all other types of rearrangement to genome divergence. This observation was consistent at all phylogenetic distances: as time since divergence increased, the number of inversions remained constant. A similar finding was made by Hirabayashi and Owens (Hirabayashi and Owens, 2023). Inversions likely occur at a high rate within plant genomes (Huang and Rieseberg, 2020), however, a low number of inversions was identified, suggesting that inversions are strongly selected against and rarely maintained. To survive underdominant selection, a novel inversion must provide enough selective advantages to outweigh its disadvantages. Inversions may provide a selective advantage by rearranging recombination loci, and linking alleles captured within their bounds. Inversion-linked alleles can be strongly selected for, if adaptive, and rise to high frequencies within populations (Rieseberg, 2001; Harringmeyer and Hoekstra, 2022). Additionally, adaptive alleles linked by inversions can be protected from strong gene flow (Yeaman, 2013). Alternatively, inversions may instead hinder adaptation. If selective conditions were to alter, previously adaptive inversions could prevent recombination from producing new allele combinations suitable for the new conditions (Rieseberg, 2001). Inversions, due to recombination suppression, also reduce effective population size and increase genetic load, as purifying selection can not purge linked deleterious mutations (Jay *et al*., 2021). The inversions identified here, which are assumed to have survived selection, were all very large, as expected (Wellenreuther and Bernatchez, 2018), with the majority (94.93%) containing genes. Inversions are rare and contribute little to genome divergence, but are highly genic and likely play a significant role in adaptation, evolution, and speciation processes.

Translocations can have similar genomic effects to inversions (Ortiz-Barrientos, Engelstädter and Rieseberg, 2016), contributing to the development of complex phenotypes, local adaptation, and speciation by disrupting recombination (Martin *et al*., 2020). As for inversions, novel translocations that survive drift must provide enough selective advantages to outweigh their disadvantages or be removed by purifying or underdominant selection. Translocations were highly abundant between recently diverged, phylogenetically close genomes. As time since divergence increased, translocations reduced in frequency but remained common. Translocations were the most common type of large rearrangement (average size: 11.2 Kbp), mirroring results obtained by Martin et al. (Martin *et al*., 2020). Translocations were much more abundant than inversions, especially when genomes have recently diverged, suggesting that translocations are less strongly selected against than inversions, despite having a similar effect on recombination. Additionally, translocations, while highly genic (65.70%) were less genic than inversions (94.93%). The different genomic pattern observed for translocations and inversions is possibly due to the effects of local versus non-local changes to recombination. Meiotic recombination may be more disrupted when reordered recombination loci are close to their location of origin. If true, purifying selection acts more strongly on inversions than on translocations. Translocations are common and along with duplications are a major contributor to genome divergence, possibly aiding in adaptation, evolution, and speciation processes. However, as the effects and mechanisms of translocations have been less studied than other rearrangements (Robberecht *et al*., 2013), it remains to be seen if they are more likely to have functional/adaptive significance.

To further investigate the potential importance of the syntenic, rearranged, and unaligned genome regions identified in our study further research using genome-wide association studies (GWAS) of phenotypes measured on seedlings in pots or field trials, as well as landscape and genome-wide genotyping for Genome Environment Association (GEA) scans for adaptive rearrangements are needed. Within species derived rearrangements are predicted to be predominantly neutral and exist at low frequencies, while others rising to higher frequencies could be true lineage-specific adaptive rearrangements. With additional genomes from populations, the frequency of rearrangements within each species could be assessed. This would provide insight into the functional significance of the widespread genomic rearrangements we have found and potentially identify rearrangements conferring adaptive traits across the landscape.

The *de novo* assembled *Eucalyptus* genomes used in this study were scaffolded based on homology to *E. grandis* (Myburg *et al*., 2014). Using non-species-specific scaffolding risks incorrectly joining contigs, resulting in the inflation or deflation of synteny and introduction of false positive and false negative rearrangements. However, *Eucalyptus* are known to have a largely conserved karyotype (Potts and Wiltshire, 1997; Booth *et al*., 2015; Grattapaglia *et al*., 2015; Butler *et al*., 2017; Supple *et al*., 2018; Healey *et al*., 2021). Furthermore, a previous study has demonstrated the viability of such a scaffolding strategy for *Eucalyptus* genomes in the absence of species-specific scaffolding data such as Hi-C, genetic maps, or optical maps (Ferguson *et al*., 2022). Lastly, our pseudo-chromosomes were observed to consist of very few contigs, suggesting that scaffolding is unlikely to be a significant source of bias.

*Eucalyptus* contains over 800 species that exist across a wide geographic and environmental range, while retaining a largely conserved karyotype (Potts and Wiltshire, 1997; Booth *et al*., 2015; Grattapaglia *et al*., 2015; Butler *et al*., 2017; Supple *et al*., 2018), which makes the genus ideal to study plant genome evolution. Here we assembled representative genomes of 33 species, creating one of the most comprehensive datasets to study plant genome evolution. These genomes provide a genus-wide resource to study genome rearrangements, and support future *Eucalyptus* research that require genomic references. Our findings suggest that following divergence, genome architecture is highly fragmented predominantly by rearrangements. As genomes continue to diverge, genome architecture continues to be slowly lost. Additionally, as genomes diverge they increasingly become unalignable due to the divergence of duplications and translocations. Syntenic regions also contribute to the growing unalignable proportion of genomes, but at a slower rate than rearrangements. Duplications and translocations are potentially the greatest contributors to functional and genome divergence, aiding in the development of complex phenotypes, and local adaptation. Inversions occur at consistently low rates, contributing little to genome architecture loss or accumulation of unalignable sequences. However, inversions were highly genic, much more so than either duplications or translocations, and likely also play a crucial role in the development of complex phenotypes, and local adaptation. Genome architecture results from a complex interaction of positive, neutral, and negative forces, all of which contribute to the evolution, divergence, and adaptability of species (Koonin, 2009; Huang and Rieseberg, 2020; Mérot *et al*., 2020). However, owing to technical limitations, the evolution of genome architecture and its role within biology is not well understood (Lynch *et al*., 2011; Cortés *et al*., 2018; Jiggins, 2019). Here, by describing the pattern of genome architecture as time since divergence increases of 33 *Eucalyptus* genomes, we contribute to a better understanding of the evolution of plant genomes. Rearrangements, along with polyploidy, TEs, and other genome evolutionary mechanisms, play an important role in plant genome evolution (Galindo-González *et al*., 2017; Marques, Meier and Seehausen, 2019; Meudt *et al*., 2021). Further research in other plant lineages is required to assess the prominence of rearrangements upon genome evolution.

## Materials and Methods

### Sampling

Eucalyptus species used in this study were collected throughout multiple locations in Australia, which are detailed in supplementary results. The majority of collected species are living collections with accession numbers at the Australian National Botanic Gardens (Canberra, Australian Capital Territory (ACT)) and Currency Creek Arboretum (Currency Creek, South Australia). Additional samples were sourced from the Australian National University (Acton, ACT) the National Arboretum Canberra (Molonglo Valley, ACT), the University of Tasmania Herbarium (Sandy Bay, Tasmania) and within Eucalyptus woodlands of southern Tasmania. Leaves were placed in plastic zip-lock bags, lightly sprayed with water to keep them moist and transported to the lab as soon as possible, where they were washed with water and stored at -80°C until DNA extraction.

### DNA extraction, sequencing, and basecalling

To extract high-molecular weight DNA from recalcitrant Eucalyptus samples, we developed two methods. Initially we combined a protocol to purify nuclei with hexylene glycol (Bolger et al., 2014) with a magnetic bead-based DNA extraction protocol (Mayjonade et al., 2016), which was further developed and is available on Protocols.io in detail (Jones and Borevitz, 2019). This was further optimised and developed, which led to the second method of adopting a sorbitol pre-wash of homogenate (Inglis et al., 2018) to wash crude nuclei instead of isolating pure nuclei, followed by a magnetic bead-based DNA extraction, according to (Jones et al., 2021). We found this method to be more time and resource efficient, hence we switched to this method for all subsequent high-molecular weight DNA extractions. For each Eucalyptus sample, the method which was used is listed within the supplementary material, Table Supplementary Table S1, with the two methods being referred to as nuclei and sorbitol respectively.

After isolating high-molecular weight DNA, we further purified and size selected the DNA by using a PippinHT (Sage Science). The DNA was size selected for fragments ≥ 20 kb or ≥ 40 kb depending on DNA yield and molecular weight, which is listed in the supplementary material, Table S1, for each sample. Two Oxford Nanopore Technologies long-read native DNA sequencing libraries were prepared for each species according to the manufacturer’s protocol 1D genomic DNA by ligation (SQK-LSK109). E. marginata was an exception, which had one ligation library as described but the second was a transposome library prep, according to the manufacturer’s protocol for rapid sequencing (SQK-RAD004). Sequencing was performed on MinION Mk1B devices using two FLO-MIN106D R9.4.1 flow cells per species. Sequencing output was improved when ONT Flow Cell Wash Kits (EXP-WSH003 and EXP-WSH004) were made available, whereby flow cells were washed when sequencing declined, primed again and more library was loaded, according to the manufacturer’s instructions. After sequencing was complete, the fast5 reads were basecalled with ONT Guppy (versions: 3.3.0, 4.0.11, 4.0.14, and 4.0.15; See Supplementary Table S14 for per species versions).

We complemented the long-read sequencing with highly accurate Illumina short-read sequencing for later use in genome polishing of the long-read de novo assemblies. Illumina short-read, whole-genome DNA sequencing libraries were generated using a cost-optimised, transposome protocol based on Illumina Nextera DNA prep methods (Jones et al., 2023). The pooled libraries were then size selected for fragments with insert sizes between 350 and 600 bp with a PippinHT (Sage Science). Multiplexed sequencing with other projects was performed on a NovaSeq 6000 (Illumina), using a lane of an S4 flow cell with a 300 cycle kit (150 bp paired-end sequencing), at the Biomolecular Resource Facility, Australian National University, Australian Capital Territory, Australia.

### *De novo* assembly

*De novo* assembly and annotation was performed using the long-read *de novo* plant assembly protocol developed by Ferguson et al. (Ferguson, Jones and Borevitz, 2022). Briefly, fastq reads are quality screened, removing DNA control strand, sequencing adaptors and low quality read ends (the first and last 200 bp), short reads (>1 Kbp in length), and low quality reads (average quality <Q7), using the NanoPack set of tools (De Coster *et al*., 2018). Curated reads are next assembled using the long-read assembler Canu (versions: 1.9 and 2.0) (Koren *et al*., 2017), which assembles high-quality *Eucalyptus* genomes (Ferguson *et al*., 2022). Assemblies were filtered of contamination (non-plant contigs), assembly artefact, plasmid, and haplotig (contigs that span the same genomic region but originate from different parental chromosomes) contigs using Blobtools (Laetsch and Blaxter, 2017) and Purge Haplotigs (version: 1.1.0) (Roach, Schmidt and Borneman, 2018). Next, all assemblies were long-read and then short-read polished, using assembly reads and Illumina reads originating from the same individual as used for assembly. Long-read polishing was performed with Racon (Vaser *et al*., 2017), and short-read with Pilon (version: 1.3.1) (Walker *et al*., 2014). Long-read polishing made use of the long-read aligner minimap2 (version: 2.17) (Li, 2018, p. 2), while short-read polishing used (version: 0.7.17) (Li, 2013). Next, assemblies were filtered to remove all contigs less than 1 Kbp in length. We chose this contig length threshold so as to maximise genome contiguity while removing all contigs too small to contain a gene. Finally, assemblies were scaffolded using homology to *E. grandis* (Myburg *et al*., 2014). Scaffolding was performed with RaGOO (version 1.1) (Alonge *et al*., 2019) and minimap2.

After assembly all genomes were quality assessed using Benchmarking Universal Single-Copy Orthologs (BUSCO; version 5; database: eudicots_odb10.2020-09-10) (Manni *et al*., 2021), long terminal repeat assembly index (version: 2.9.0; LAI) (Ou, Chen and Jiang, 2018), and assembly statistics.

### Transposon and gene annotation, and gene ortho grouping

Genome repeat and gene annotation was also performed using the long-read *de novo* plant assembly protocol developed by Ferguson et al. (Ferguson, Jones and Borevitz, 2022). Firstly, *de novo* repeat libraries were created for each genome using EDTA (version: 1.9.6) (Ou *et al*., 2019), subsequently all genomes were repeat annotated with RepeatMasker (version: 4.0.9) (Smit, Hubley and Green, 2020). All genomes were repeat soft-masked and subsequently annotated for genes. Gene annotation was performed with BRAKER (version 2.1.5; Brůna *et al*., 2021, p. 2) using GeneMark-EP (version: 4; Brůna, Lomsadze and Borodovsky, 2020). Gene transcript sequences for model training were obtained from the National Center for Biotechnology Information (NCBI) (Sayers *et al*., 2021). Included in gene training data were all Myrtaceae (Taxonomy ID: 3931) and Arabidopsis thaliana (Taxonomy ID: 3702) transcripts. All gene candidates were grouped into orthogroups using Orthofinder (version: 2.5.4) (Emms and Kelly, 2019). Using DIAMOND (Buchfink, Reuter and Drost, 2021) Orthofinder aligned all gene transcripts, grouping those with >40% identity and achieving an e-score < 0.001.

### Genome synteny, rearrangement and unaligned annotation

Identification of all shared sequences began by aligning all pairwise combinations of genomes with the MUMmer (version 3) (Kurtz *et al*., 2004) tool nucmer (parameters: -- maxmatch -l 40 -b 500 -c 200). Nucmer first identifies all shared 40-mers between genomes and their locations. Next, 40-mers within 500bp are clustered, creating a list of collinear blocks or alignments. Lastly, using MUMmer’s delta-filter tool alignments are filtered, removing all alignments less than 200 bp in length and less than 80% similar. A low 80% alignment similarity score was used as *Eucalyptus* are highly heterozygous (Murray *et al*., 2019), and a more stringent similarity score may incorrectly filter out real alignments.

Having identified all shared sequences we next annotated all syntenic, rearranged (inverted, translocated, and duplicated), and unaligned (sequence that only exists in one genome, resulting from either an insertion, deletion or sequence divergence) sequence between pairwise genomes using SyRI (version: 1.5) (Goel *et al*., 2019).

SyRI’s use of a directed acyclic graph results in genomes being annotated for smaller regions, which when occurring in an unbroken series of a single type, get combined. The resulting output includes both levels of annotations, smaller more fragmented, and larger and more contiguous. We make use of the larger and more continuous alignments. Additionally, we combined inverted duplications with duplications, and inverted translocations with translocations.

### Phylogeny

Using highly conserved and single-copy genes, BUSCO genes, we built a eucalypt phylogenetic tree describing the evolutionary relationships between all genomes included in this study. The phylogenetic tree included four previously and identically assembled genomes for *E. albens*, *E. melliodora*, *E. sideroxylon*, and *C. calophylla*, creating a dataset of 36 genomes. To begin, fasta sequences for all single-copy BUSCO genes found within 30+ genomes were collected. Using masce (version 2.03; Ranwez *et al*., 2018, p. 2) multi-sequence alignment was performed individually on all genes. As errors within gene MSAs will subsequently lead to errors in phylogenetic inferences, we trimmed and filtered all gene MSAs. Gene sequence errors were detected and removed using HmmCleaner version: 0.180750; (Di Franco *et al*., 2019). HmmCleaner uses a profile-hidden Markov model to identify sequence segments that poorly fit the gene MSA and subsequently removes them. Errors resulting from poor alignments were removed using report2AA (parameters: - min_NT_to_keep_seq 30, -min_seq_to_keep_site 4,-dist_isolate_AA 3, - min_homology_to_keep_seq 0.5, -min_percent_NT_at_ends 0.7) from the macse program. report2AA removed sites within MSAs that included less than 30 genomes, had less than 4 informative characters, or had isolated sites (site was more than 3 characters away from the next non-gap character). Additionally, report2AA removed genome from MSAs that had less than 50% homology to another genome within the MSA, and trimmed both MSA ends that had less than 70% of aligned sites as nucleotides (i.e. 26+ genomes had to have a non-gap character). Additionally, as a result of filtering and trimming MSAs of low quality are removed.

Individual gene trees were constructed for all filtered and trimmed MSAs using iqtree (version: 1.6.12; Nguyen *et al*., 2015). Finally, all gene trees were concatenated into a single file, from which a species tree was generated using Astral III version 5.7.3; (Zhang *et al*., 2018). The resulting species tree was manually rooting at the *Angophora/Corymbia* and *Eucalyptus* branch, using Figtree (version: 1.4.4; Rambaut, no date).

## Data access

Sequencing data and reference genomes generated in this project are publicly available on the Sequence Read Archive (SRA) and NCBI genome repository under BioProject PRJNA509734. Gene predictions and SyRI annotations have been deposited in FigShare and are available at: https://figshare.com/projects/Plant_genome_evolution_in_the_genus_Eucalyptus_driven_by_structural_rearrangements_that_promote_sequence_divergence/97010. All analysis scripts created and used by this project have been deposited within our github repository: https://github.com/fergsc/33-Eucs.

## Authors’ contributions

Scott Ferguson led the project and ran all the analysis. Ashley Jones managed, developed, and performed DNA sampling and sequencing. The project was conceived and designed by all authors. Scott Ferguson wrote the first manuscript draft. All authors contributed to writing and review of the final manuscript.

## Acknowledgements

We would like to thank the Australian National Botanic Gardens in Canberra, Australia for providing plant samples and associated metadata. This research acknowledges the support provided by the Director of National Parks, the park staff of the Australian National Botanic Gardens, and Parks Australia. The views expressed in this document do not necessarily represent the views of the Australian Government.

We thank Dean Nicolle, owner of the Currency Creek Arboretum, South Australia, for providing samples and support for this project. We also thank Yoav Daniel Bar-Ness, Giant Tree Expeditions, for collecting E. regnans (the Centurion). We also thank Tamera Beath, David Stanley, Cynthia Torkel, Rob Lanfear, Wang Weiwen and Brad Potts for their support and collecting samples.

We would like to thank the ACRF Biomolecular Resource Facility at the John Curtin School of Medical Research, ANU in Canberra, Australia, where Oxford Nanopore Technologies PromethION and Illumina NovaSeq 6000 sequencing was conducted. This research acknowledges the support provided by NCRIS-enabled Bioplatforms Australia infrastructure.

Computational resources were provided by the Australian Government through the National Computational Infrastructure (NCI) under the ANU Merit Allocation Scheme.

This research is funded by The Australian Research Council Centre of Excellence in Plant Energy Biology and an Australian Research Council Discovery Grant.

Kevin Murray and Scott Ferguson were supported by Australian Government Research Training Program scholarships.

