## Supplemental figures and tables for "Plant genome evolution in the genus *Eucalyptus* driven by structural rearrangements that promote sequence divergence"

### Supplementary Figures and Tables

| Species | DNA extraction method | Size selection (kb) | Actual Coverage | N50 (Kbp) | Number of reads | Total size (Mbp) | Longest read (Kbp) |
|---|---|---|---|---|---|---|---|
| ***A. floribunda*** | Nuclei | 20 | 68.5 | 30,913 | 1,165,378 | 26,218.73 | 234.42 |
| ***C. maculata*** | Nuclei | 20 | 62.0 | 32,785 | 1,068,357 | 24,727.52 | 186.19 |
| ***E. brandiana*** | Nuclei | 20 | 59.0 | 25,064 | 1,572,473 | 29,661.57 | 185.11 |
| ***E. caleyi*** | Sorbitol | 20 | 44.2 | 38,986 | 981,871 | 25,568.27 | 245.74 |
| ***E. camaldulensis*** | Nuclei | 20 | 43.1 | 26,925 | 1,246,437 | 23,709.64 | 236.32 |
| ***E. cladocalyx*** | Nuclei | 20 | 44.3 | 22,608 | 1,603,737 | 23,813.99 | 871.14 |
| ***E. cloeziana*** | Nuclei | 20 | 35.8 | 23,441 | 1,106,171 | 16,977.71 | 202.46 |
| ***E. coolabah*** | Nuclei | 40 | 33.5 | 38,840 | 843,066 | 19,942.69 | 258.17 |
| ***E. curtisii*** | Nuclei | 20 | 61.2 | 26,200 | 1,482,403 | 26,364.96 | 184.51 |
| ***E. dawsonii*** | Sorbitol | 20 | 31.4 | 39,210 | 833,846 | 21,893.54 | 340.53 |
| ***E. decipiens*** | Nuclei | 20 | 36.9 | 29,788 | 937,250 | 21,488.84 | 241.66 |
| ***E. erythrocorys*** | Nuclei | 20 | 43.4 | 30,515 | 994,422 | 23,167.70 | 201.42 |
| ***E. fibrosa*** | Sorbitol | 20 | 39.1 | 40,269 | 808,634 | 22,664.55 | 239.76 |
| ***E. globulus*** | Nuclei | 20 | 27.5 | 26,580 | 789,684 | 14,775.47 | 196.77 |
| ***E. grandis*** | Nuclei | 40 | 26.6 | 40,622 | 566,332 | 16,120.84 | 223.69 |
| ***E. guilfoylei*** | Sorbitol | 20 | 44.9 | 37,750 | 748,484 | 21,030.20 | 238.04 |
| ***E. lansdowneana*** | Sorbitol | 20 | 33.5 | 40,454 | 766,963 | 20,885.35 | 264.89 |
| ***E. leucophloia*** | Nuclei | 20 | 42.3 | 31,376 | 1,112,561 | 23,719.81 | 220.51 |
| ***E. marginata*** | Nuclei | 20 | 37.6 | 19,330 | 2,030,612 | 18,996.49 | 403.50 |
| ***E. melliodora x E. sideroxylon*** | Sorbitol | 20 | 63.5 | 28,218 | 2,096,002 | 38,010.87 | 493.59 |
| ***E. microcorys*** | Nuclei | 20 | 83.8 | 26,233 | 1,891,415 | 36,501.91 | 259.59 |
| ***E. ANBG9806169*** | Nuclei | 20 | 45.0 | 23,768 | 1,361,586 | 22,622.88 | 125.92 |
| ***E. paniculata*** | Sorbitol | 20 | 37.5 | 36,669 | 878,971 | 21,750.73 | 223.73 |
| ***E. pauciflora*** | NA | NA | 72.3 | 26.93 | 2,583,680 | 35,691.93 | 537.11 |
| ***E. polyanthemos*** | Sorbitol | 20 | 43.8 | 30,700 | 1,317,377 | 26,018.59 | 220.24 |
| ***E. pumila*** | Nuclei | 20 | 40.7 | 25,051 | 1,632,628 | 21,358.64 | 165.40 |
| ***E. regnans*** | Sorbitol | 40 | 47.9 | 46,297 | 718,316 | 23,254.44 | 253.70 |
| ***E. shirleyi*** | Sorbitol | 20 | 44.3 | 44,993 | 862,028 | 26,095.39 | 273.96 |
| ***E. tenuipes*** | Nuclei | 40 | 48.7 | 41,950 | 663,327 | 19,070.76 | 296.92 |
| ***E. victrix*** | Sorbitol | 20 | 50.6 | 41,561 | 1,149,096 | 27,764.76 | 270.78 |
| ***E. viminalis*** | Nuclei | 20 | 32.0 | 22,308 | 1,535,956 | 17,640.60 | 163.61 |
| ***E. virginea*** | Nuclei | 20 | 47.7 | 25,319 | 1,406,739 | 25,089.92 | 164.74 |

**Supplementary Table S1.** Raw read library statistics. Actual coverage is calculated after assembly from the final genome size.

| Species | Coverage | Coverage lost | N50 (Kbp) | Number of reads | Total size (Mbp) | Longest read (Kbp) |
|---|---|---|---|---|---|---|
| ***A. floribunda*** | 62.9 | 5.6 | 30,973 | 961,453 | 24,080.30 | 234.02 |
| ***C. maculata*** | 58.0 | 4 | 32,784 | 902,490 | 23,154.27 | 185.79 |
| ***E. brandiana*** | 54.7 | 4.3 | 24,964 | 1,318,778 | 27,465.22 | 184.71 |
| ***E. caleyi*** | 41.1 | 3.1 | 39,257 | 797,371 | 23,780.80 | 245.34 |
| ***E. camaldulensis*** | 39.8 | 3.3 | 26,890 | 1,036,444 | 21,912.23 | 235.92 |
| ***E. cladocalyx*** | 40.3 | 4 | 22,553 | 1,357,574 | 21,658.32 | 167.28 |
| ***E. cloeziana*** | 32.4 | 3.4 | 23,421 | 891,520 | 15,359.12 | 150.84 |
| ***E. coolabah*** | 31.1 | 2.4 | 39,047 | 696,864 | 18,498.71 | 257.77 |
| ***E. curtisii*** | 56.8 | 4.4 | 26,212 | 1,260,701 | 24,482.11 | 184.11 |
| ***E. dawsonii*** | 29.1 | 2.3 | 39,530 | 705,446 | 20,309.35 | 340.13 |
| ***E. decipiens*** | 34.8 | 2.1 | 29,702 | 818,918 | 20,220.97 | 232.70 |
| ***E. erythrocorys*** | 40.1 | 3.3 | 30,501 | 854,347 | 21,394.22 | 201.02 |
| ***E. fibrosa*** | 36.7 | 2.4 | 40,481 | 710,154 | 21,285.37 | 239.36 |
| ***E. globulus*** | 24.7 | 2.8 | 26,783 | 625,059 | 13,252.63 | 196.37 |
| ***E. grandis*** | 24.7 | 1.9 | 40,618 | 478,371 | 14,946.13 | 223.29 |
| ***E. guilfoylei*** | 41.8 | 3.1 | 37,759 | 663,545 | 19,576.72 | 237.64 |
| ***E. lansdowneana*** | 31.1 | 2.4 | 40,677 | 662,726 | 19,393.64 | 264.49 |
| ***E. leucophloia*** | 39.4 | 2.9 | 31,506 | 966,679 | 22,088.48 | 220.11 |
| ***E. marginata*** | 32.6 | 5 | 19,997 | 1,441,634 | 16,472.64 | 266.42 |
| ***E. melliodora x E. sideroxylon*** | 58.3 | 5.2 | 28,356 | 1,760,405 | 34,919.15 | 203.52 |
| ***E. microcorys*** | 78.0 | 5.8 | 26,255 | 1,628,396 | 33,963.49 | 259.19 |
| ***E. ANBG9806169*** | 41.3 | 3.7 | 23,609 | 1,134,657 | 20,769.73 | 125.52 |
| ***E. paniculata*** | 35.3 | 2.2 | 36,846 | 769,367 | 20,476.85 | 223.33 |
| ***E. pauciflora*** | 73.3 | 0 | 26,926 | 2,583,680 | 35,691.93 | 537.11 |
| ***E. polyanthemos*** | 40.4 | 3.4 | 30,888 | 1,121,971 | 24,048.44 | 219.84 |
| ***E. pumila*** | 36.8 | 3.9 | 25,254 | 1,217,779 | 19,269.00 | 165.00 |
| ***E. regnans*** | 45.6 | 2.3 | 46,254 | 611,946 | 22,142.04 | 212.45 |
| ***E. shirleyi*** | 41.7 | 2.6 | 45,127 | 740,464 | 24,593.44 | 273.56 |
| ***E. tenuipes*** | 46.0 | 2.7 | 41,997 | 574,526 | 18,007.59 | 296.52 |
| ***E. victrix*** | 46.9 | 3.7 | 42,005 | 923,073 | 25,731.32 | 266.30 |
| ***E. viminalis*** | 28.8 | 3.2 | 22,386 | 955,619 | 15,865.49 | 163.21 |
| ***E. virginea*** | 44.4 | 3.3 | 25,299 | 1,196,234 | 23,361.01 | 164.34 |

**Supplementary Table S2.** Filtered read library statistics. Statistics describing read libraries after trimming, length filtering, and quality filtering. Note: *E. pauciflora* reads were filtered and trimmed before being randomly down sampled, hence no loss of sequence.

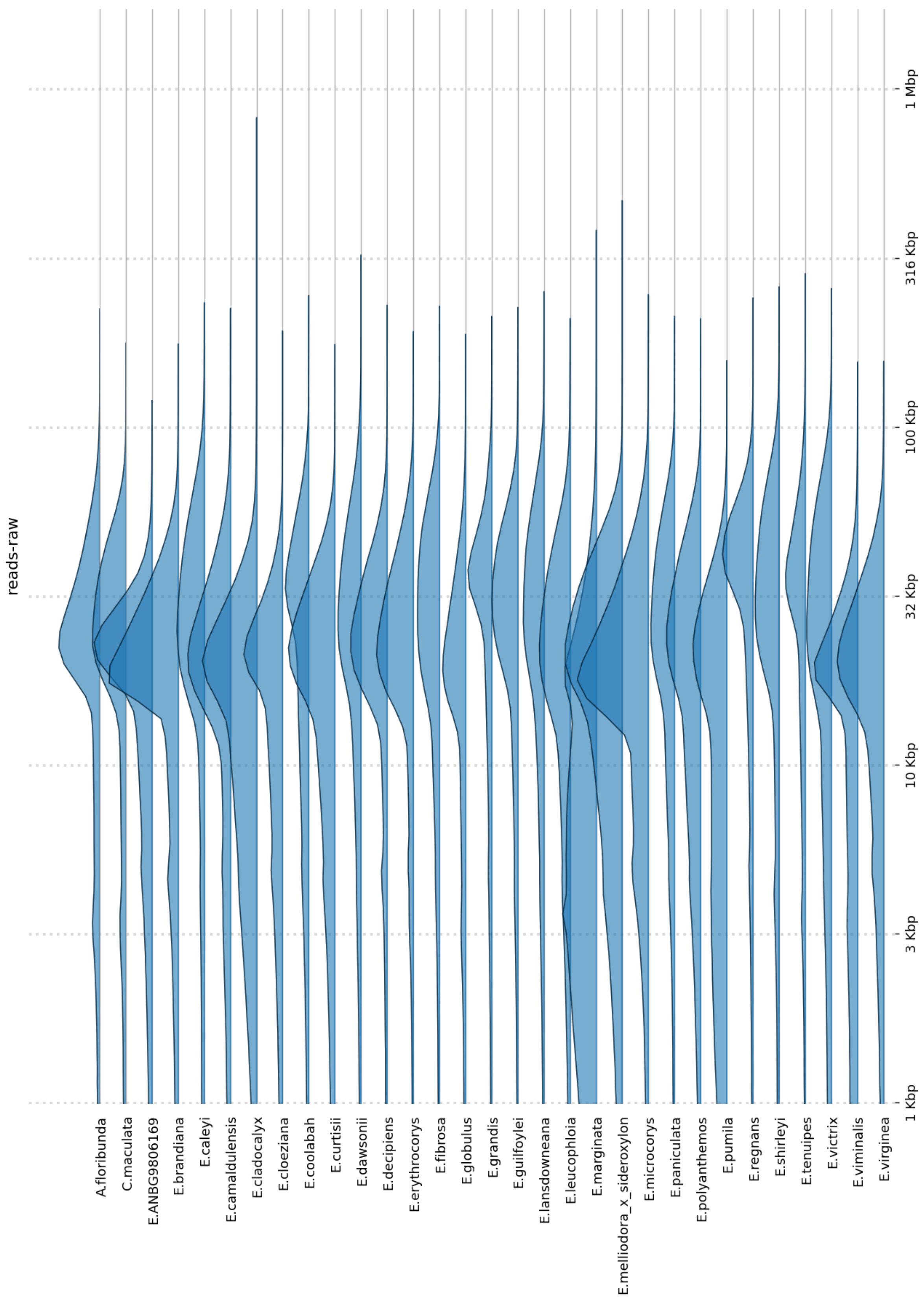


**Supplementary Figures S1.** Distribution of read lengths of raw read libraries

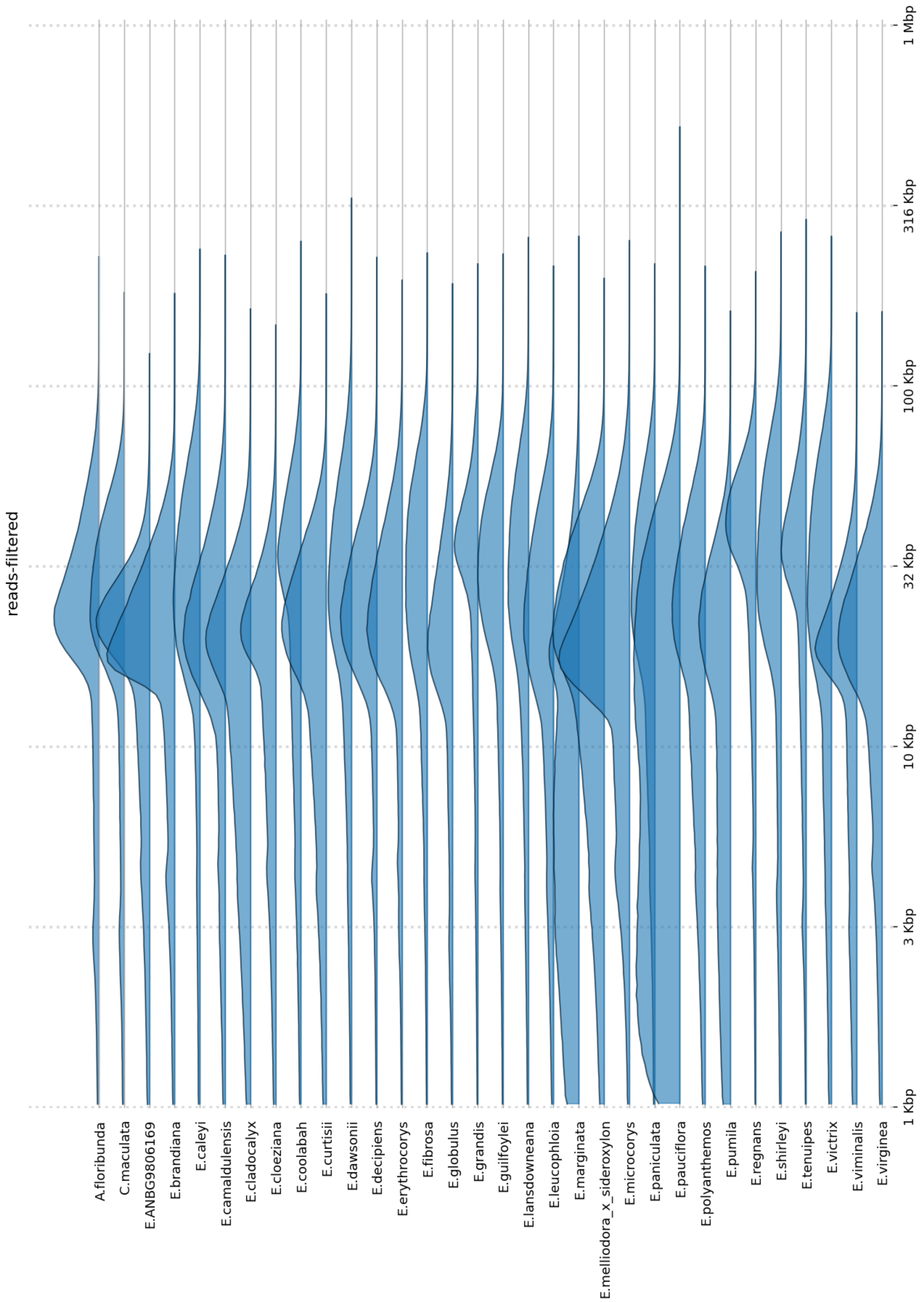


**Supplementary Figures S2.** Distribution of read lengths of read libraries aftering trimming, length filtering, and quality filtering.

| | Contamination | Haplotigs | | Assembly artifacts | |
|---|---|---|---|---|---|
| **Species** | **Total Size (Kbp)** | **Total Size (Mbp)** | **% of Genome** | **Total Size (Mbp)** | **% of Genome** |
| ***A. floribunda*** | None | 253.37 | 36.74% | 53.73 | 7.79% |
| ***C. maculata*** | None | 265.09 | 36.74% | 57.48 | 7.97% |
| ***E. brandiana*** | 161.18 | 38.62 | 7.01% | 10.05 | 1.82% |
| ***E. caleyi*** | 3,448.47 | 458.40 | 42.15% | 46.71 | 4.29% |
| ***E. camaldulensis*** | None | 394.82 | 39.70% | 49.34 | 4.96% |
| ***E. cladocalyx*** | 33.10 | 383.48 | 41.54% | 1.86 | 0.20% |
| ***E. cloeziana*** | None | 188.67 | 27.07% | 33.94 | 4.87% |
| ***E. coolabah*** | None | 381.48 | 38.25% | 21.07 | 2.11% |
| ***E. curtisii*** | None | 234.46 | 35.06% | 3.51 | 0.53% |
| ***E. dawsonii*** | 18,227.31 | 426.59 | 37.01% | 9.88 | 0.86% |
| ***E. decipiens*** | 40.93 | 384.89 | 37.82% | 51.03 | 5.02% |
| ***E. erythrocorys*** | None | 115.73 | 16.78% | 40.38 | 5.86% |
| ***E. fibrosa*** | 1.38 | 553.42 | 48.66% | 3.49 | 0.31% |
| ***E. globulus*** | None | 228.73 | 29.17% | 18.52 | 2.36% |
| ***E. grandis*** | None | 280.46 | 30.84% | 23.47 | 2.58% |
| ***E. guilfoylei*** | None | 93.14 | 16.44% | 4.66 | 0.82% |
| ***E. lansdowneana*** | 7.00 | 463.32 | 42.41% | 5.02 | 0.46% |
| ***E. leucophloia*** | 241.53 | 446.00 | 44.08% | 4.61 | 0.46% |
| ***E. marginata*** | 34.66 | 290.90 | 35.09% | 32.13 | 3.88% |
| ***E. melliodora x E. sideroxylon*** | 133.04 | 507.89 | 45.74% | 3.94 | 0.36% |
| ***E. microcorys*** | 133.10 | 228.43 | 30.66% | 81.15 | 10.89% |
| ***E. ANBG9806169*** | None | 231.78 | 30.09% | 36.05 | 4.68% |
| ***E. paniculata*** | 249.95 | 501.04 | 46.14% | 4.47 | 0.41% |
| ***E. pauciflora*** | 107.77 | 441.32 | 47.36% | 3.62 | 0.39% |
| ***E. polyanthemos*** | None | 552.62 | 48.03% | 3.35 | 0.29% |
| ***E. pumila*** | 42.45 | 272.67 | 34.07% | 3.18 | 0.40% |
| ***E. regnans*** | 267.24 | 370.84 | 40.60% | 56.32 | 6.17% |
| ***E. shirleyi*** | None | 442.43 | 42.74% | 3.67 | 0.35% |
| ***E. tenuipes*** | None | 204.32 | 31.75% | 47.28 | 7.35% |
| ***E. victrix*** | None | 482.44 | 46.63% | 3.68 | 0.36% |
| ***E. viminalis*** | 43.43 | 296.85 | 34.18% | 20.42 | 2.35% |
| ***E. virginea*** | None | 281.01 | 32.69% | 52.26 | 6.08% |
| **Average** | 724.14 | 334.23 | 36.15% | 24.70 | 3.04% |

**Supplementary Table S3.** Amount and proportion of sequence removed from genomes during contamination filtering, haplotig purging, and artifact filtering.

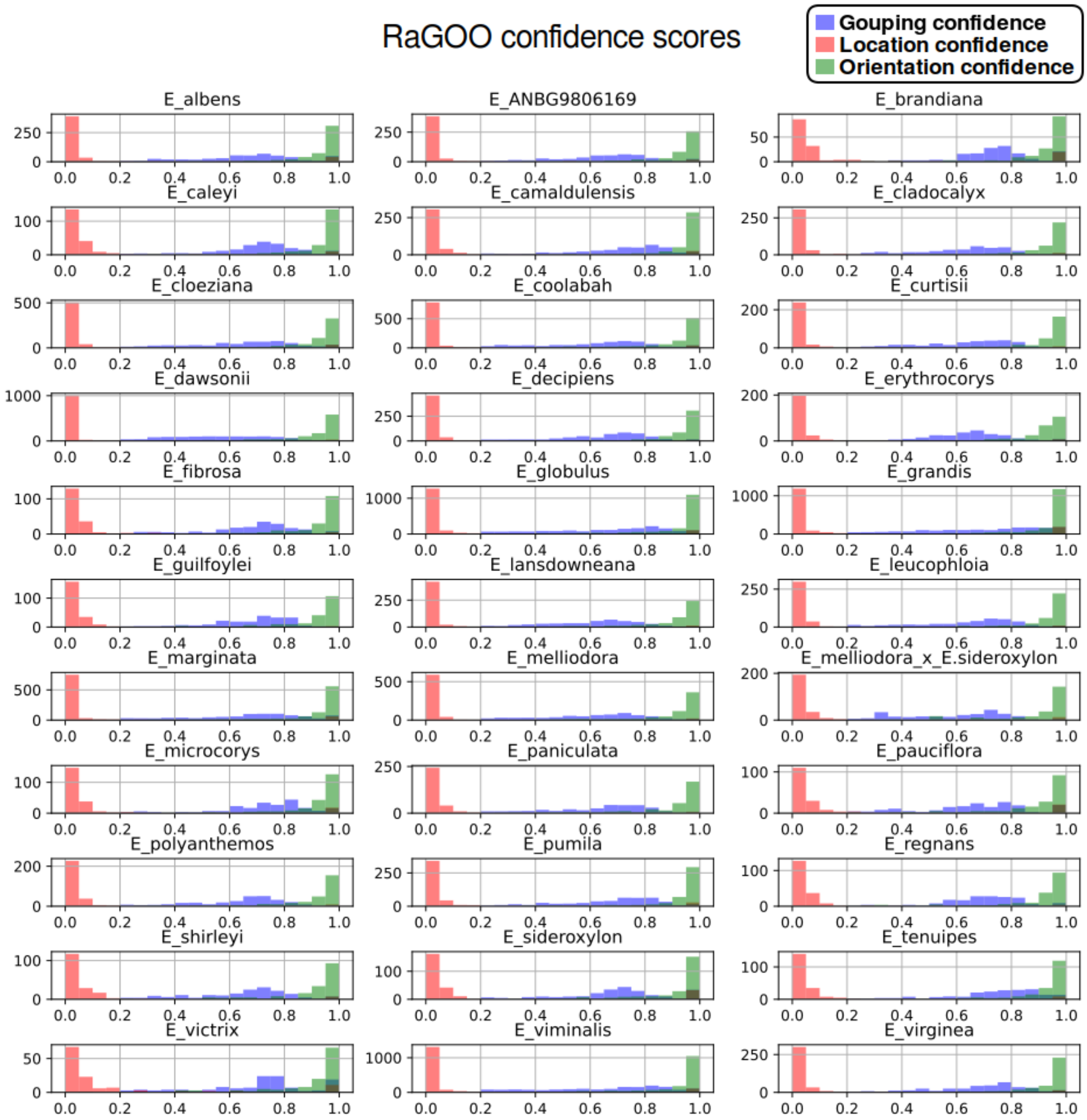


**Supplementary Figure S3.** Confidence scores reported by RaGOO during scaffolding of genomes. Grouping confidence = assigning contigs to a chromosome; location confidence = ordering contigs relative to each other along the scaffold; orientation confidence = orienting contigs within scaffolds.

| species | Complete (single-copy + duplicated) | single-copy | duplicated | fragmented | missing | LAI |
|---|---|---|---|---|---|---|
| ***A. floribunda*** | 96.82% | 92.78% | 4.04% | 1.33% | 1.85% | 14.5 |
| ***C. maculata*** | 97.25% | 94.93% | 2.32% | 1.12% | 1.63% | 15.92 |
| ***E. brandiana*** | 98.11% | 95.06% | 3.05% | 1.03% | 0.86% | 23.85 |
| ***E. caleyi*** | 96.47% | 90.80% | 5.67% | 1.59% | 1.93% | 18.24 |
| ***E. camaldulensis*** | 96.73% | 92.09% | 4.64% | 1.33% | 1.93% | 16.99 |
| ***E. cladocalyx*** | 97.59% | 93.29% | 4.30% | 1.12% | 1.29% | 18.53 |
| ***E. cloeziana*** | 97.12% | 93.55% | 3.57% | 1.59% | 1.29% | 19.06 |
| ***E. coolabah*** | 95.44% | 88.52% | 6.92% | 2.15% | 2.41% | 15.9 |
| ***E. curtisii*** | 97.29% | 93.21% | 4.08% | 1.42% | 1.29% | 18.34 |
| ***E. dawsonii*** | 97.51% | 83.10% | 14.40% | 0.95% | 1.55% | 17.01 |
| ***E. decipiens*** | 96.99% | 91.49% | 5.50% | 1.33% | 1.68% | 18.87 |
| ***E. erythrocorys*** | 97.55% | 94.41% | 3.14% | 0.95% | 1.50% | 20.18 |
| ***E. fibrosa*** | 96.73% | 87.96% | 8.77% | 1.63% | 1.63% | 17.49 |
| ***E. globulus*** | 96.69% | 91.87% | 4.82% | 1.25% | 2.06% | 17.46 |
| ***E. grandis*** | 96.09% | 90.11% | 5.98% | 1.59% | 2.32% | 17.11 |
| ***E. guilfoylei*** | 98.02% | 94.84% | 3.18% | 0.90% | 1.07% | 16.39 |
| ***E. lansdowneana*** | 97.12% | 86.76% | 10.36% | 1.12% | 1.76% | 19.46 |
| ***E. leucophloia*** | 96.99% | 90.58% | 6.41% | 1.29% | 1.72% | 17.91 |
| ***E. marginata*** | 96.17% | 90.93% | 5.25% | 1.72% | 2.11% | 19.58 |
| ***E. melliodora x E. sideroxylon*** | 97.72% | 91.70% | 6.02% | 1.03% | 1.25% | 17.96 |
| ***E. microcorys*** | 97.21% | 93.90% | 3.31% | 1.25% | 1.55% | 16.2 |
| ***E. ANBG9806169*** | 96.86% | 93.29% | 3.57% | 1.29% | 1.85% | 22.16 |
| ***E. paniculata*** | 97.12% | 90.58% | 6.53% | 1.20% | 1.68% | 18.58 |
| ***E. pauciflora*** | 97.25% | 93.29% | 3.96% | 1.25% | 1.50% | 20.29 |
| ***E. polyanthemos*** | 96.82% | 89.85% | 6.96% | 1.55% | 1.63% | 17.52 |
| ***E. pumila*** | 97.38% | 93.29% | 4.08% | 1.42% | 1.20% | 17.74 |
| ***E. regnans*** | 97.25% | 93.47% | 3.78% | 1.25% | 1.50% | 20.18 |
| ***E. shirleyi*** | 97.29% | 88.99% | 8.30% | 1.16% | 1.55% | 19.89 |
| ***E. tenuipes*** | 96.39% | 92.56% | 3.83% | 1.81% | 1.81% | 15.07 |
| ***E. victrix*** | 96.65% | 91.32% | 5.33% | 1.46% | 1.89% | 18.71 |
| ***E. viminalis*** | 96.47% | 91.83% | 4.64% | 1.29% | 2.24% | 16.5 |
| ***E. virginea*** | 97.08% | 92.30% | 4.77% | 1.16% | 1.76% | 17.69 |

**Supplementary Table S4**. BUSCO and LAI scores for all genomes.

| | N50 (Kbp) | Number of contigs | Size (Mbp) | Longest contig (Mbp) |
|---|---|---|---|---|
| ***A. floribunda*** | 1,139.19 | 2,857 | 689.71 | 10.21 |
| ***C. maculata*** | 1,540.02 | 3,174 | 721.55 | 12.94 |
| ***E. brandiana*** | 6,578.56 | 802 | 551.18 | 24.53 |
| ***E. caleyi*** | 1,805.12 | 2,810 | 1,087.60 | 25.88 |
| ***E. camaldulensis*** | 939.69 | 4,219 | 994.59 | 8.75 |
| ***E. cladocalyx*** | 953.07 | 4,591 | 923.15 | 16.82 |
| ***E. cloeziana*** | 855.51 | 4,172 | 697.06 | 7.22 |
| ***E. coolabah*** | 552.28 | 4,066 | 997.21 | 5.38 |
| ***E. curtisii*** | 1,332.99 | 2,628 | 668.67 | 16.95 |
| ***E. dawsonii*** | 474.20 | 4,951 | 1,152.64 | 8.02 |
| ***E. decipiens*** | 747.03 | 4,718 | 1,017.59 | 6.82 |
| ***E. erythrocorys*** | 2,931.08 | 2,208 | 689.47 | 18.40 |
| ***E. fibrosa*** | 1,730.72 | 1,790 | 1,137.28 | 24.72 |
| ***E. globulus*** | 317.08 | 5,378 | 784.04 | 3.78 |
| ***E. grandis*** | 321.42 | 5,311 | 909.45 | 4.43 |
| ***E. guilfoylei*** | 3,303.65 | 1,041 | 566.64 | 21.44 |
| ***E. lansdowneana*** | 1,031.71 | 2,572 | 1,092.36 | 13.56 |
| ***E. leucophloia*** | 1,038.45 | 3,108 | 1,011.89 | 12.88 |
| ***E. marginata*** | 458.19 | 5,121 | 828.93 | 4.33 |
| ***E. melliodora x E. sideroxylon*** | 1,719.77 | 2,555 | 1,110.47 | 21.42 |
| ***E. microcorys*** | 1,549.62 | 3,462 | 745.17 | 13.19 |
| ***E. ANBG9806169*** | 1,006.91 | 4,115 | 770.36 | 14.88 |
| ***E. paniculata*** | 1,259.10 | 2,337 | 1,085.95 | 17.29 |
| ***E. pauciflora*** | 1,379.37 | 2,654 | 931.94 | 23.39 |
| ***E. polyanthemos*** | 1,287.56 | 2,369 | 1,150.67 | 22.17 |
| ***E. pumila*** | 1,064.83 | 3,313 | 800.22 | 15.45 |
| ***E. regnans*** | 1,571.93 | 2,834 | 913.33 | 22.03 |
| ***E. shirleyi*** | 2,635.00 | 1,462 | 1,035.28 | 25.64 |
| ***E. tenuipes*** | 1,548.79 | 2,406 | 643.48 | 16.67 |
| ***E. victrix*** | 3,210.46 | 1,298 | 1,034.58 | 36.56 |
| ***E. viminalis*** | 289.36 | 7,315 | 868.55 | 4.50 |
| ***E. virginea*** | 1,222.18 | 3,963 | 859.54 | 11.49 |

**Supplementary Table S5**. Raw genome statistics. Genomes have not been curated in any way.

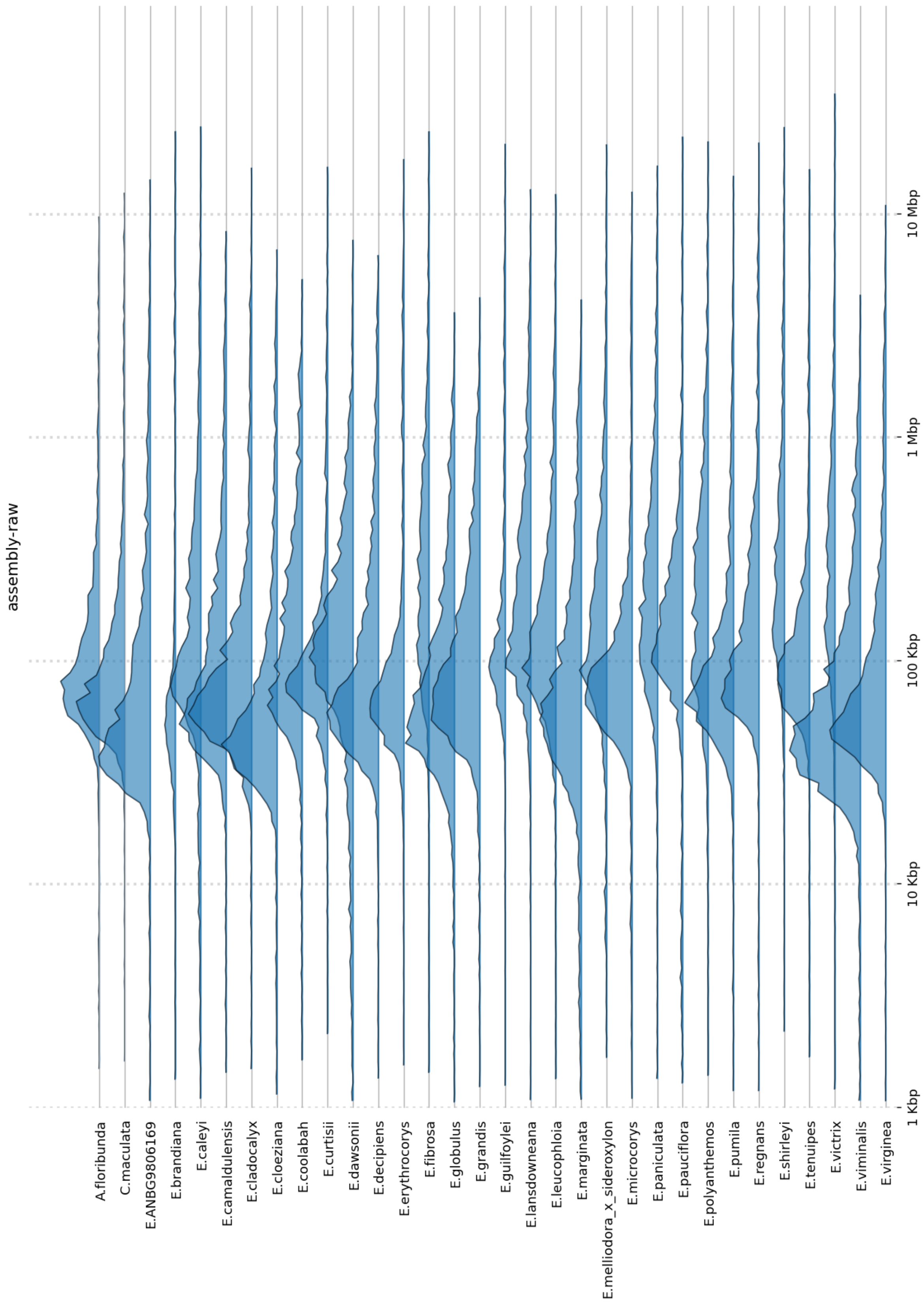


**Supplementary Figure S4**. Raw genome contig distributions. Genomes have not been curated in any way.

| | N50 (Kbp) | Number of contigs | Size (Mbp) | Longest contig (Mbp) |
|---|---|---|---|---|
| ***A. floribunda*** | 4,011.83 | 225 | 388.67 | 10.21 |
| ***C. maculata*** | 4,690.14 | 176 | 404.38 | 12.94 |
| ***E. brandiana*** | 7,278.40 | 170 | 507.20 | 24.53 |
| ***E. caleyi*** | 4,769.92 | 276 | 589.30 | 25.88 |
| ***E. camaldulensis*** | 2,481.60 | 420 | 558.53 | 8.75 |
| ***E. cladocalyx*** | 2,796.55 | 390 | 544.04 | 16.82 |
| ***E. cloeziana*** | 1,739.94 | 626 | 480.25 | 7.22 |
| ***E. coolabah*** | 1,289.21 | 937 | 606.36 | 5.38 |
| ***E. curtisii*** | 2,961.00 | 288 | 435.23 | 16.95 |
| ***E. dawsonii*** | 985.62 | 1,343 | 706.79 | 8.02 |
| ***E. decipiens*** | 1,985.74 | 553 | 590.96 | 6.82 |
| ***E. erythrocorys*** | 4,024.36 | 253 | 539.36 | 18.40 |
| ***E. fibrosa*** | 6,447.41 | 192 | 589.89 | 24.72 |
| ***E. globulus*** | 636.15 | 1,750 | 545.00 | 3.78 |
| ***E. grandis*** | 613.64 | 1,749 | 616.23 | 4.43 |
| ***E. guilfoylei*** | 4,251.44 | 209 | 472.34 | 21.44 |
| ***E. lansdowneana*** | 2,352.90 | 489 | 633.47 | 13.56 |
| ***E. leucophloia*** | 2,659.36 | 382 | 568.44 | 12.88 |
| ***E. marginata*** | 1,011.74 | 991 | 513.33 | 4.33 |
| ***E. melliodora x E. sideroxylon*** | 6,217.05 | 282 | 603.80 | 21.42 |
| ***E. microcorys*** | 4,004.79 | 234 | 440.96 | 13.19 |
| ***E. ANBG9806169*** | 2,396.19 | 477 | 507.94 | 14.88 |
| ***E. paniculata*** | 3,699.06 | 330 | 588.82 | 17.29 |
| ***E. pauciflora*** | 6,583.95 | 216 | 494.01 | 23.39 |
| ***E. polyanthemos*** | 4,657.75 | 300 | 603.25 | 22.17 |
| ***E. pumila*** | 2,492.89 | 473 | 529.71 | 15.45 |
| ***E. regnans*** | 5,259.26 | 206 | 495.05 | 22.03 |
| ***E. shirleyi*** | 6,912.49 | 181 | 597.16 | 25.64 |
| ***E. tenuipes*** | 3,433.87 | 211 | 398.08 | 16.67 |
| ***E. victrix*** | 11,095.62 | 120 | 557.15 | 36.56 |
| ***E. viminalis*** | 652.95 | 1,759 | 558.84 | 4.50 |
| ***E. virginea*** | 2,391.64 | 379 | 532.97 | 11.49 |

**Supplementary Table S6**. Final genome statistics. Genomes have been fully curated, i.e. all contaminate, haplotig, plastid, and artifact contigs have been removed. Genomes have also been polished.

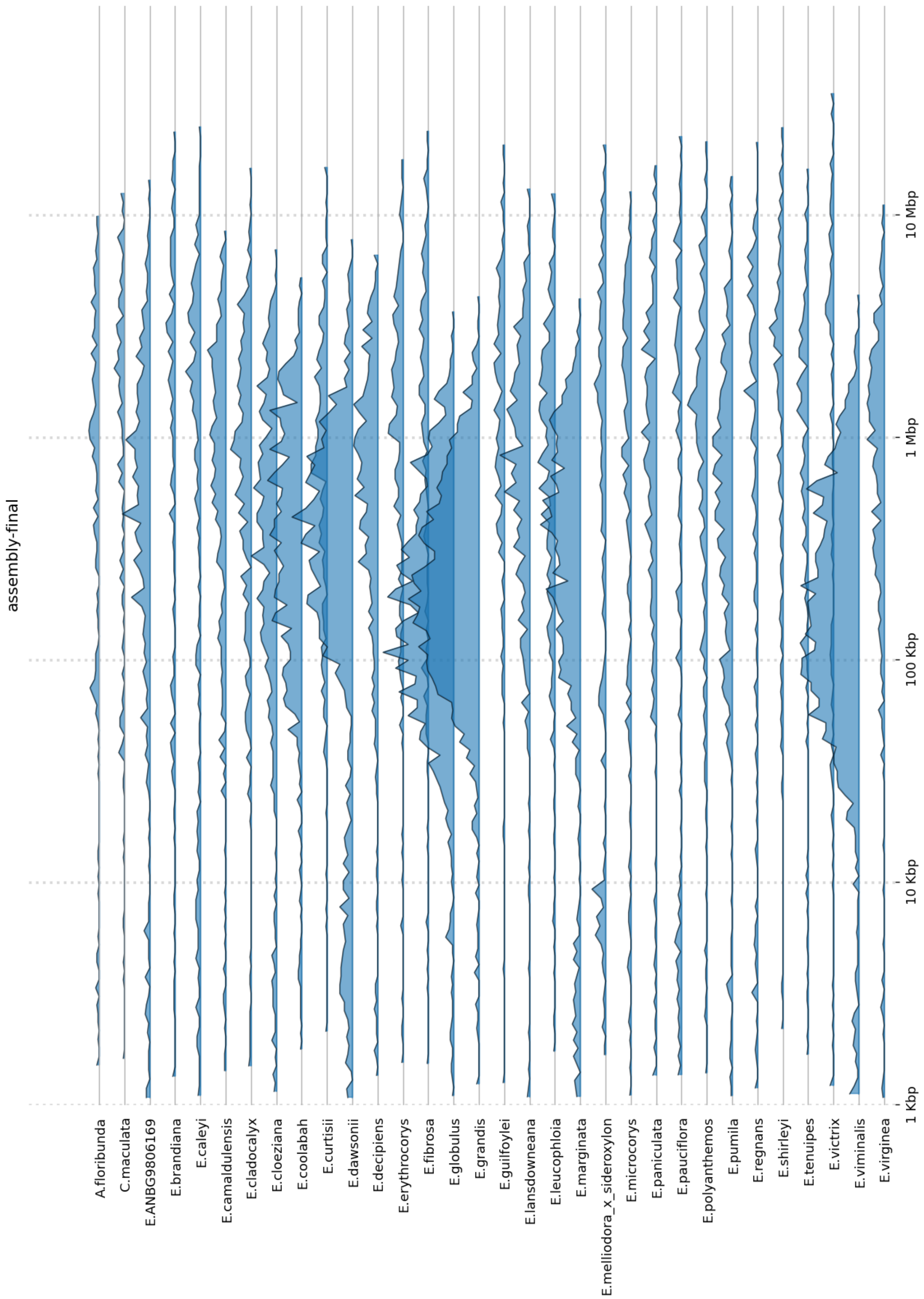


**Supplementary Figure S5**. Final genome contig distributions. Genomes have been fully curated, i.e. all contaminate, haplotig, plastid, and artifact contigs have been removed. Genomes have also been polished.

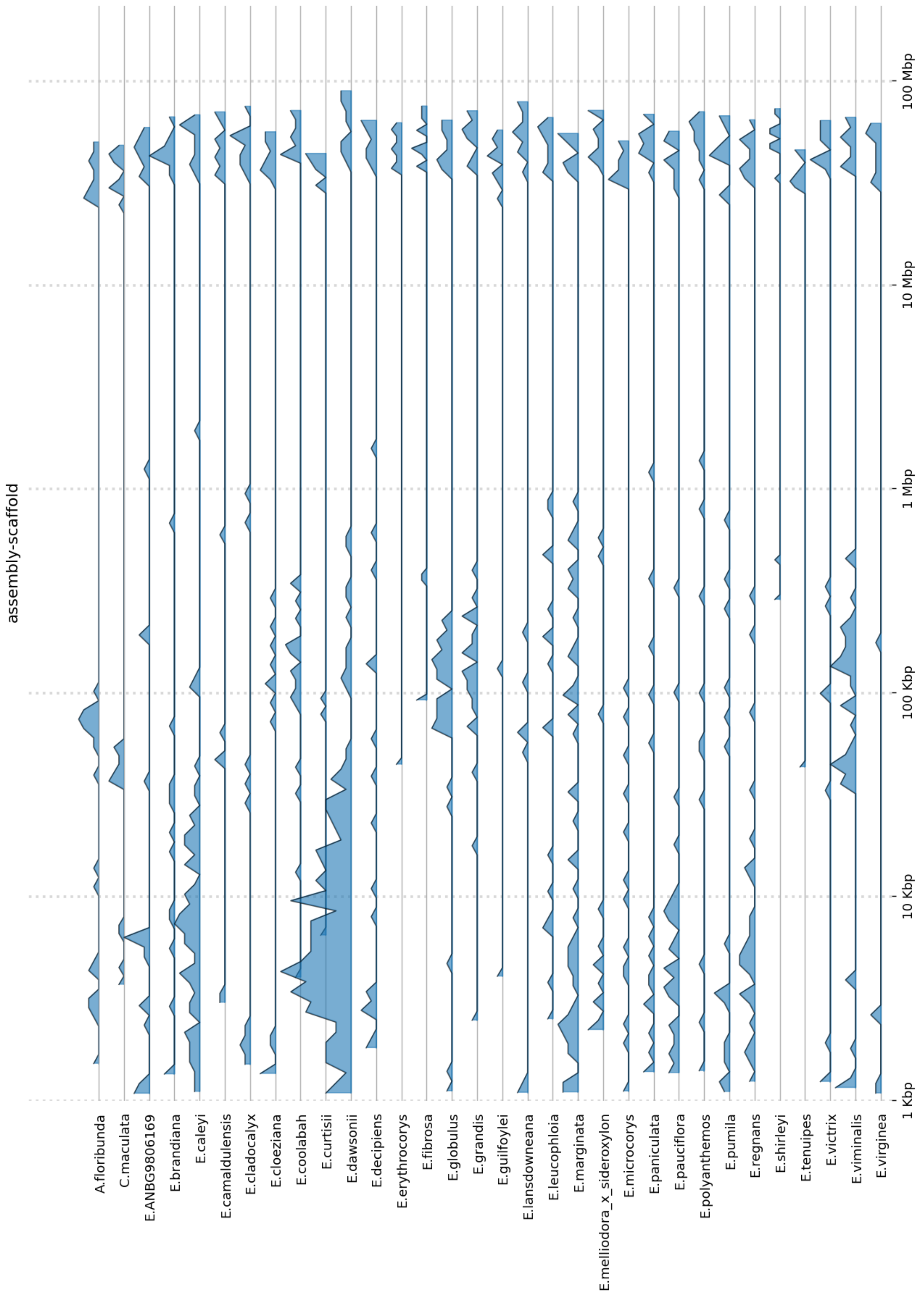


**Supplementary Figure S6**. Scaffolded genome contig distributions.

| | Proportion of TE | Proportion of simple repeats | Total repeat proportion |
|---|---|---|---|
| ***A. floribunda*** | 34.55% | 1.26% | 35.81% |
| ***C. calophylla*** | 34.79% | 1.14% | 35.93% |
| ***E. brandiana*** | 44.21% | 1.24% | 45.45% |
| ***E. caleyi*** | 46.00% | 1.19% | 47.19% |
| ***E. camaldulensis*** | 45.31% | 1.25% | 46.56% |
| ***E. cladocalyx*** | 45.85% | 1.28% | 47.13% |
| ***E. cloeziana*** | 42.57% | 1.25% | 43.82% |
| ***E. coolabah*** | 45.89% | 1.27% | 47.16% |
| ***E. curtisii*** | 41.66% | 1.31% | 42.97% |
| ***E. dawsonii*** | 45.88% | 1.17% | 47.05% |
| ***E. decipiens*** | 46.95% | 1.19% | 48.14% |
| ***E. erythrocorys*** | 47.07% | 1.30% | 48.37% |
| ***E. fibrosa*** | 45.10% | 1.22% | 46.32% |
| ***E. globulus*** | 44.29% | 1.23% | 45.52% |
| ***E. grandis*** | 46.53% | 1.17% | 47.70% |
| ***E. guilfoylei*** | 41.22% | 1.20% | 42.42% |
| ***E. lansdowneana*** | 46.10% | 1.27% | 47.37% |
| ***E. leucophloia*** | 44.37% | 1.28% | 45.65% |
| ***E. marginata*** | 43.43% | 1.21% | 44.64% |
| ***E. melliodora x E. sideroxylon*** | 46.71% | 1.19% | 47.90% |
| ***E. microcorys*** | 41.39% | 1.31% | 42.70% |
| ***E. ANBG9806169*** | 44.00% | 1.21% | 45.21% |
| ***E. paniculata*** | 44.92% | 1.32% | 46.24% |
| ***E. pauciflora*** | 43.10% | 1.23% | 44.33% |
| ***E. polyanthemos*** | 45.55% | 1.23% | 46.78% |
| ***E. pumila*** | 44.17% | 1.28% | 45.45% |
| ***E. regnans*** | 43.06% | 1.20% | 44.26% |
| ***E. shirleyi*** | 45.85% | 1.25% | 47.10% |
| ***E. tenuipes*** | 37.82% | 1.34% | 39.16% |
| ***E. victrix*** | 44.34% | 1.39% | 45.73% |
| ***E. viminalis*** | 44.57% | 1.19% | 45.76% |
| ***E. virginea*** | 43.78% | 1.29% | 45.07% |

**Supplementary Table S7.** Repeat content of all genomes.

| Species | Number of genes | Genes in orthogroup | Unassigned genes | Paralogue groups | Genes in paralogues |
|---|---|---|---|---|---|
| ***E. albens*** | 65,121 | 63,903 (98.13%) | 1,218 (1.87%) | 43 | 94 (0.14%) |
| ***E. brandiana*** | 46,176 | 45,632 (98.82%) | 544 (1.18%) | 20 | 44 (0.1%) |
| ***E. caleyi*** | 62,000 | 60,606 (97.75%) | 1,394 (2.25%) | 59 | 143 (0.23%) |
| ***E. camaldulensis*** | 51,855 | 50,949 (98.25%) | 906 (1.75%) | 33 | 80 (0.15%) |
| ***E. cladocalyx*** | 48,487 | 47,797 (98.58%) | 690 (1.42%) | 30 | 68 (0.14%) |
| ***E. cloeziana*** | 43,093 | 42,349 (98.27%) | 744 (1.73%) | 26 | 61 (0.14%) |
| ***E. coolabah*** | 51,879 | 50,855 (98.03%) | 1,024 (1.97%) | 29 | 62 (0.12%) |
| ***E. curtisii*** | 43,242 | 42,182 (97.55%) | 1,060 (2.45%) | 79 | 194 (0.45%) |
| ***E. dawsonii*** | 77,763 | 73,579 (94.62%) | 4,184 (5.38%) | 207 | 481 (0.62%) |
| ***E. decipiens*** | 49,909 | 49,012 (98.2%) | 897 (1.8%) | 43 | 101 (0.2%) |
| ***E. erythrocorys*** | 47,392 | 45,659 (96.34%) | 1,733 (3.66%) | 299 | 966 (2.04%) |
| ***E. fibrosa*** | 53,954 | 53,276 (98.74%) | 678 (1.26%) | 18 | 40 (0.07%) |
| ***E. globulus*** | 51,735 | 50,781 (98.16%) | 954 (1.84%) | 23 | 47 (0.09%) |
| ***E. grandis*** | 55,224 | 53,939 (97.67%) | 1,285 (2.33%) | 50 | 114 (0.21%) |
| ***E. guilfoylei*** | 54,024 | 52,853 (97.83%) | 1,171 (2.17%) | 95 | 236 (0.44%) |
| ***E. lansdowneana*** | 59,574 | 58,536 (98.26%) | 1,038 (1.74%) | 42 | 93 (0.16%) |
| ***E. leucophloia*** | 53,950 | 52,869 (98%) | 1,081 (2%) | 43 | 101 (0.19%) |
| ***E. marginata*** | 48,255 | 47,059 (97.52%) | 1,196 (2.48%) | 61 | 151 (0.31%) |
| ***E. melliodora*** | 63,907 | 62,839 (98.33%) | 1,068 (1.67%) | 88 | 224 (0.35%) |
| ***E. melliodora x E. sideroxylon*** | 64,299 | 63,447 (98.67%) | 852 (1.33%) | 35 | 73 (0.11%) |
| ***E. microcorys*** | 44,367 | 43,578 (98.22%) | 789 (1.78%) | 34 | 88 (0.2%) |
| ***E. ANBG9806169*** | 48,845 | 47,891 (98.05%) | 954 (1.95%) | 55 | 134 (0.27%) |
| ***E. paniculata*** | 55,821 | 55,011 (98.55%) | 810 (1.45%) | 25 | 57 (0.1%) |
| ***E. pauciflora*** | 47,634 | 46,796 (98.24%) | 838 (1.76%) | 59 | 156 (0.33%) |
| ***E. polyanthemos*** | 56,111 | 55,253 (98.47%) | 858 (1.53%) | 21 | 44 (0.08%) |
| ***E. pumila*** | 59,604 | 58,260 (97.75%) | 1,344 (2.25%) | 63 | 142 (0.24%) |

| | | | | | |
|---|---|---|---|---|---|
| ***E. regnans*** | 45,608 | 44,753 (98.13%) | 855 (1.87%) | 49 | 120 (0.26%) |
| ***E. shirleyi*** | 55,816 | 55,022 (98.58%) | 794 (1.42%) | 34 | 81 (0.15%) |
| ***E. sideroxylon*** | 60,360 | 59,278 (98.21%) | 1,082 (1.79%) | 27 | 59 (0.1%) |
| ***E. tenuipes*** | 41,622 | 40,587 (97.51%) | 1,035 (2.49%) | 50 | 123 (0.3%) |
| ***E. victrix*** | 51,710 | 51,070 (98.76%) | 640 (1.24%) | 21 | 44 (0.09%) |
| ***E. viminalis*** | 50,955 | 50,223 (98.56%) | 732 (1.44%) | 26 | 56 (0.11%) |
| ***E. virginea*** | 51,559 | 50,667 (98.27%) | 892 (1.73%) | 35 | 74 (0.14%) |

**Supplementary Table S8.** Genome orthogrouping statistics. For each genome lists the number of annotated genes, the number and proportion of genes placed within an orthogroup, the number and proportion of genes not placed within an orthogroup, the number and proportion of genes placed within a species-specific orthogroup, and the number of paralogues for each species.

| | |
|---|---|
| Number of species | 33 |
| Number of genes | 1,761,851 |
| Number of genes in orthogroups | 1,726,511 |
| Number of unassigned genes | 35,340 |
| Percentage of genes in orthogroups | 97.99% |
| Percentage of unassigned genes | 2.01% |
| Number of orthogroups | 68,245 |
| Number of species-specific orthogroups | 1,822 |
| Number of genes in species-specific orthogroups | 4,551 |
| Percentage of genes in species-specific orthogroups | 0.26% |
| Mean orthogroup size | 25.3 |

**Supplementary Table S9.** Orthogrouping summary statistics.

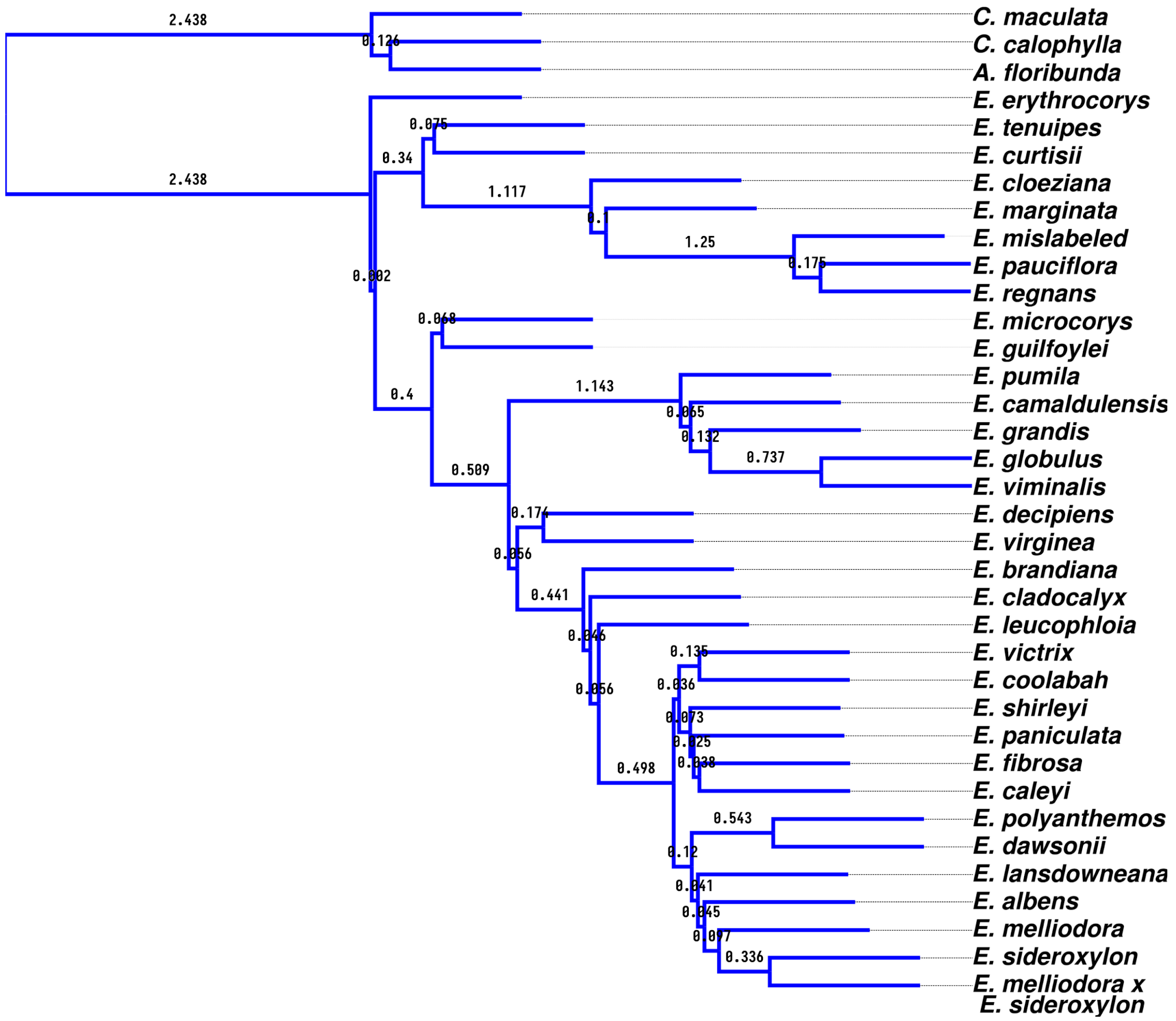


**Supplementary Figure S7.** Maximum likelihood species phylogeny estimated from 1,674 BUSCO genes. Branch lengths are in coalescent units, as calculated by Astral III. Astral doesn't calculate branch lengths for tips, as tips are a single genome.

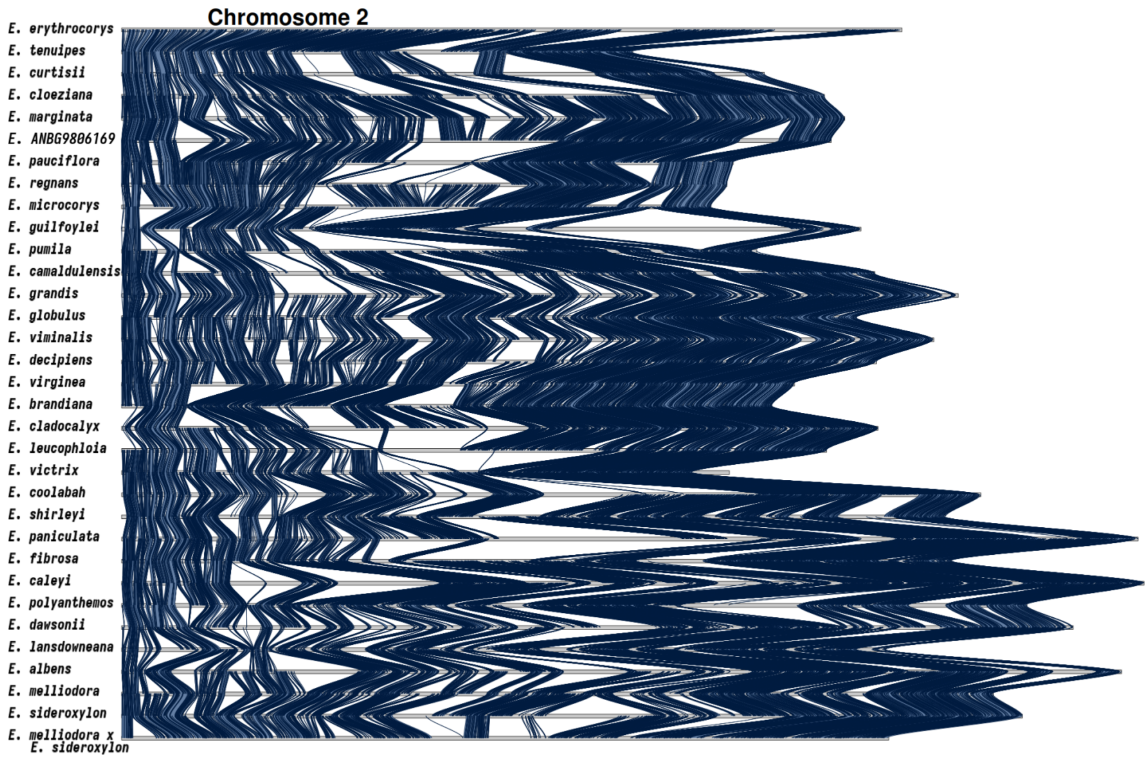


**Supplementary Figure S8. Synteny karyotype of chromosome 2.** The blue ribbons between karyotypes indicate the presence of syntenic sequences between species pairs. In all other regions synteny has become lost. Synteny is lost to either rearrangements (inverted, translocated, or duplicated), sequence divergence, loss or gain.

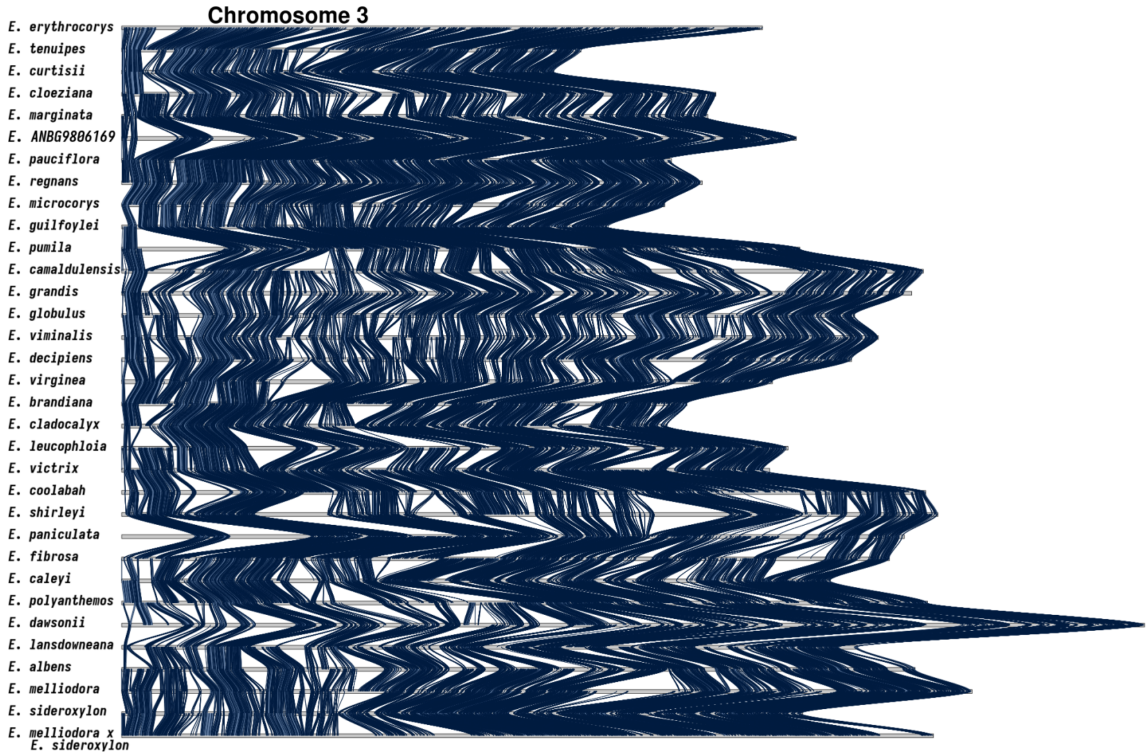


**Supplementary Figure S9. Synteny karyotype of chromosome 3.** The blue ribbons between karyotypes indicate the presence of syntenic sequences between species pairs. In all other regions synteny has become lost. Synteny is lost to either rearrangements (inverted, translocated, or duplicated), sequence divergence, loss or gain.

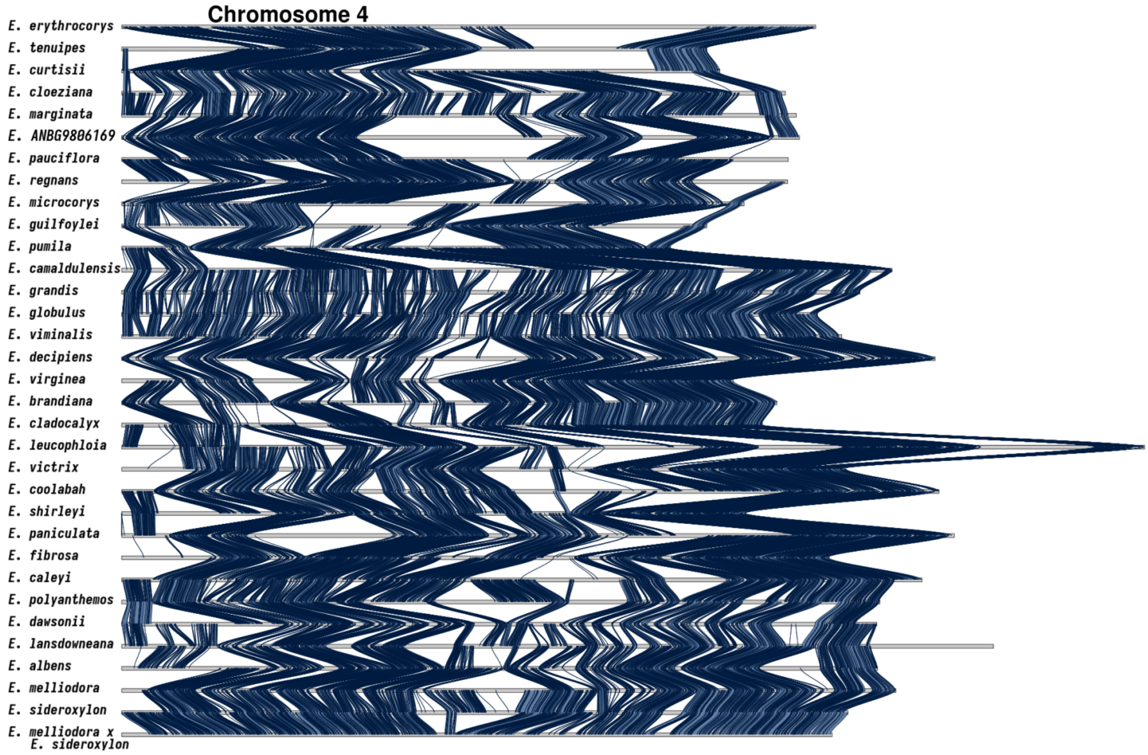


**Supplementary Figure S10. Synteny karyotype of chromosome 4.** The blue ribbons between karyotypes indicate the presence of syntenic sequences between species pairs. In all other regions synteny has become lost. Synteny is lost to either rearrangements (inverted, translocated, or duplicated), sequence divergence, loss or gain.

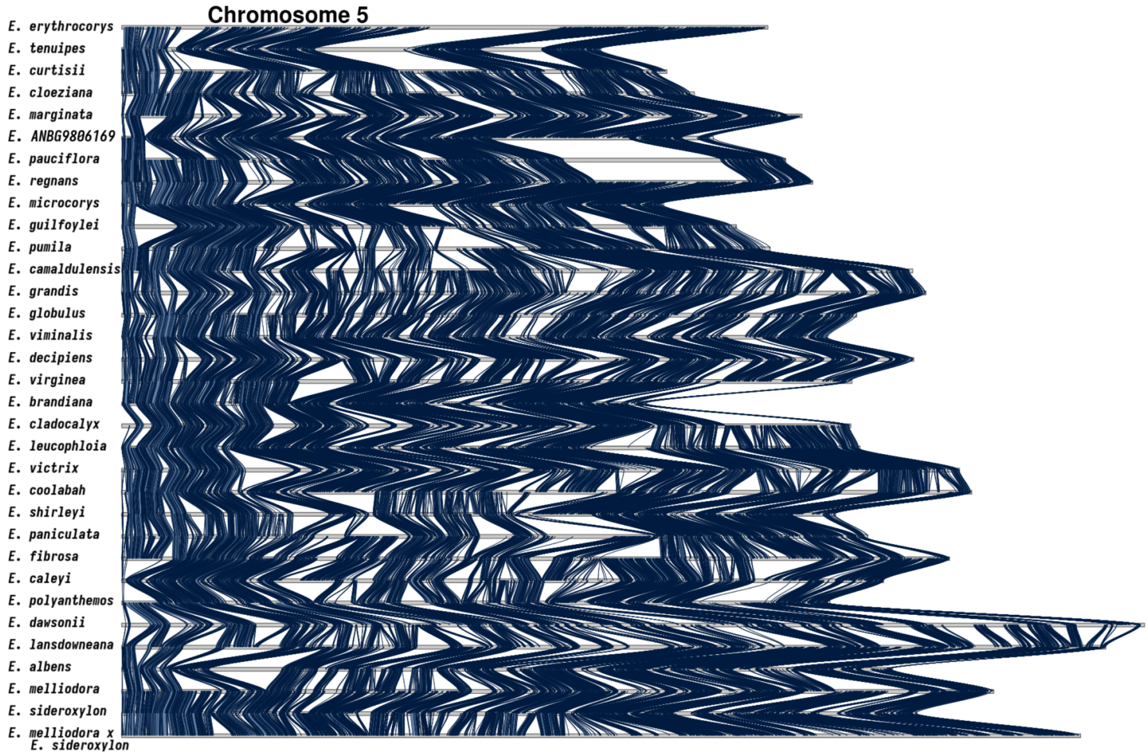


**Supplementary Figure S11. Synteny karyotype of chromosome 5.** The blue ribbons between karyotypes indicate the presence of syntenic sequences between species pairs. In all other regions synteny has become lost. Synteny is lost to either rearrangements (inverted, translocated, or duplicated), sequence divergence, loss or gain.

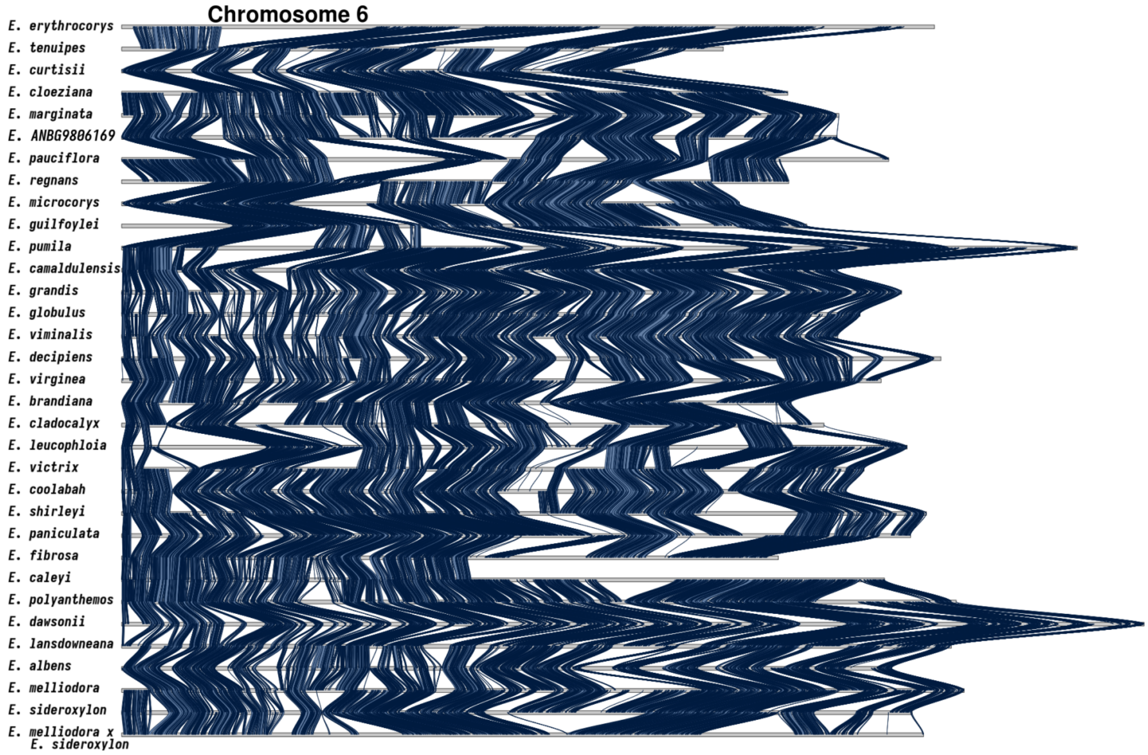


**Supplementary Figure S12. Synteny karyotype of chromosome 6.** The blue ribbons between karyotypes indicate the presence of syntenic sequences between species pairs. In all other regions synteny has become lost. Synteny is lost to either rearrangements (inverted, translocated, or duplicated), sequence divergence, loss or gain.

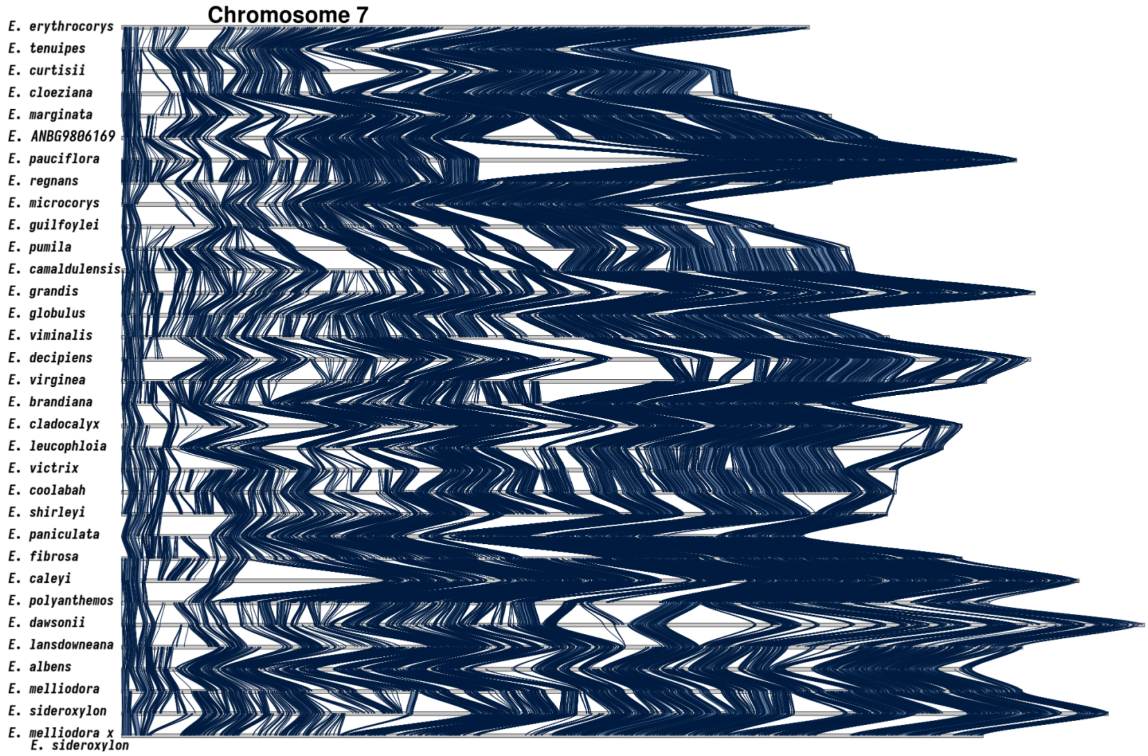


**Supplementary Figure S13. Synteny karyotype of chromosome 7.** The blue ribbons between karyotypes indicate the presence of syntenic sequences between species pairs. In all other regions synteny has become lost. Synteny is lost to either rearrangements (inverted, translocated, or duplicated), sequence divergence, loss or gain.

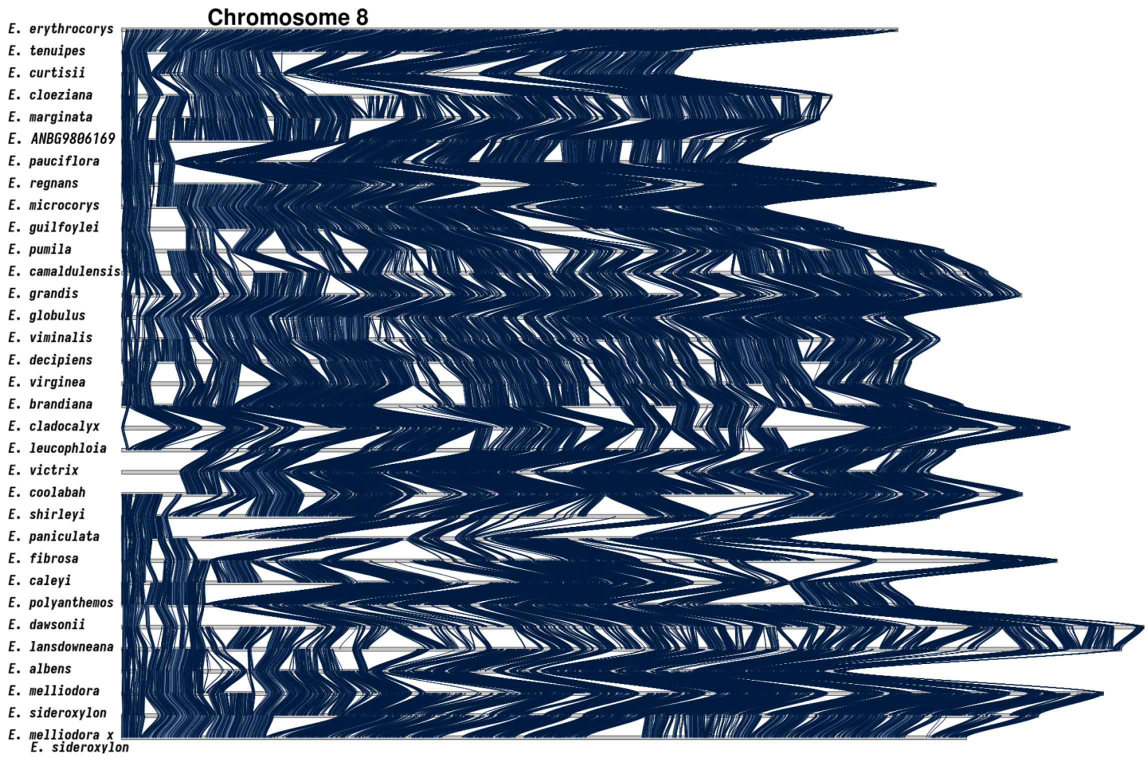


**Supplementary Figure S14. Synteny karyotype of chromosome 8.** The blue ribbons between karyotypes indicate the presence of syntenic sequences between species pairs. In all other regions synteny has become lost. Synteny is lost to either rearrangements (inverted, translocated, or duplicated), sequence divergence, loss or gain.

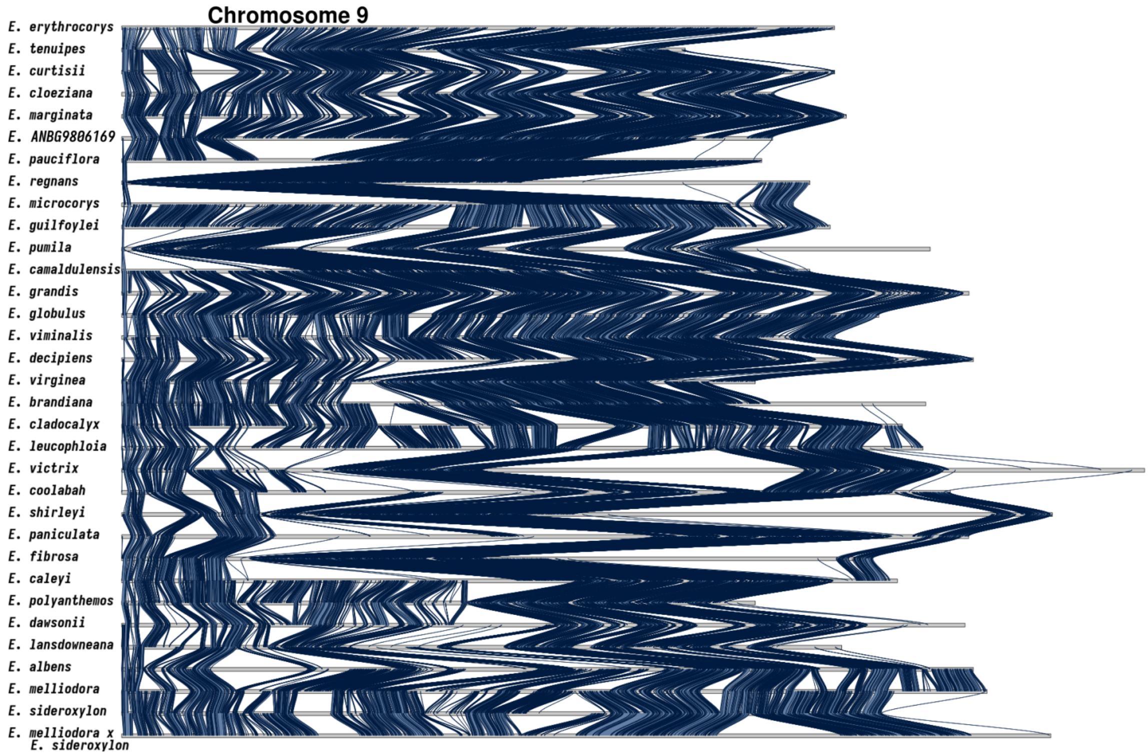


**Supplementary Figure S15. Synteny karyotype of chromosome 9.** The blue ribbons between karyotypes indicate the presence of syntenic sequences between species pairs. In all other regions synteny has become lost. Synteny is lost to either rearrangements (inverted, translocated, or duplicated), sequence divergence, loss or gain.

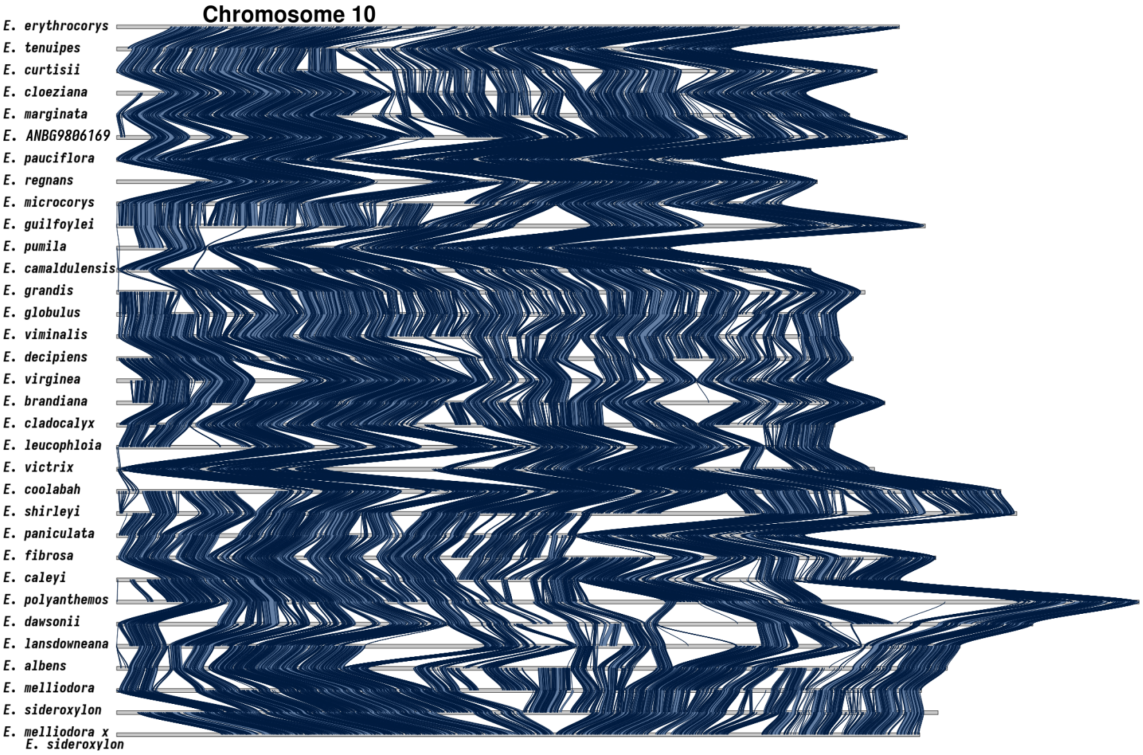


**Supplementary Figure S16. Synteny karyotype of chromosome 10.** The blue ribbons between karyotypes indicate the presence of syntenic sequences between species pairs. In all other regions synteny has become lost. Synteny is lost to either rearrangements (inverted, translocated, or duplicated), sequence divergence, loss or gain.

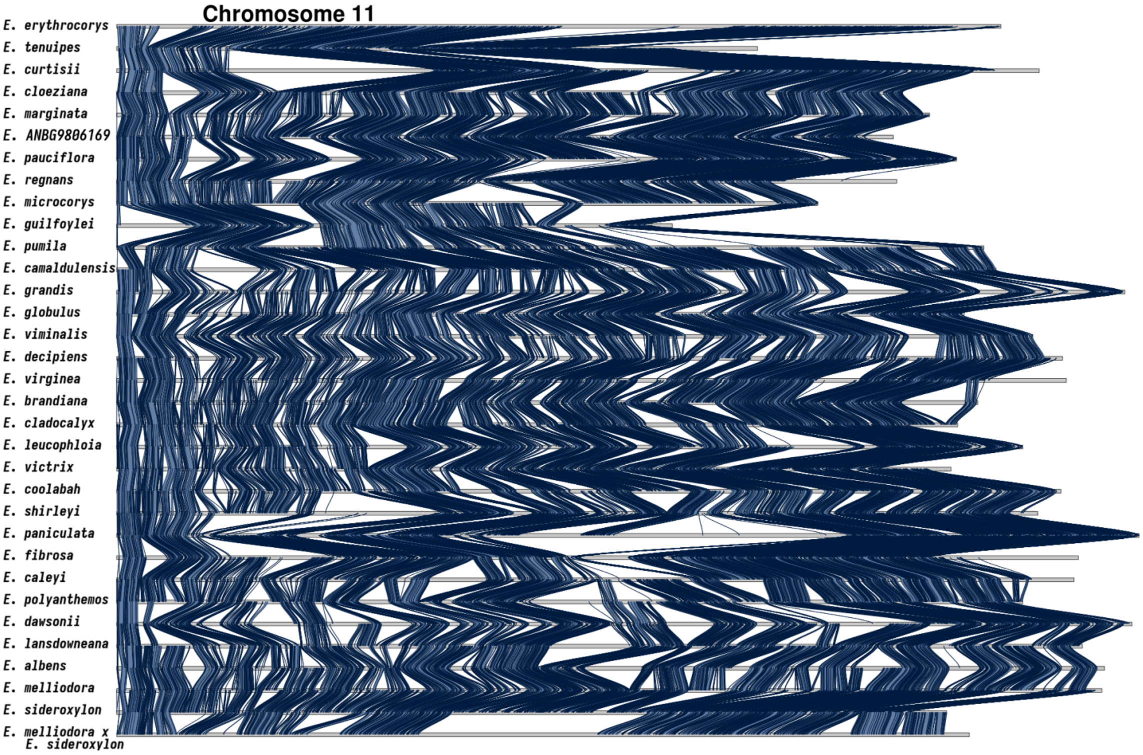


**Supplementary Figure S17. Synteny karyotype of chromosome 11.** The blue ribbons between karyotypes indicate the presence of syntenic sequences between species pairs. In all other regions synteny has become lost. Synteny is lost to either rearrangements (inverted, translocated, or duplicated), sequence divergence, loss or gain.

| | albens | brandiana | caleyi | camaldulensis | cladocalyx | cloeziana | coolabah | curtisii | dawsonii | decipiens | erythrocorys |
|---|---|---|---|---|---|---|---|---|---|---|---|
| albens | | 39.2% | 43.1% | 35.8% | 36.8% | 32.0% | 43.8% | 28.9% | 37.5% | 36.9% | 28.4% |
| brandiana | 46.7% | | 49.2% | 45.5% | 46.9% | 41.8% | 50.2% | 36.6% | 42.5% | 48.2% | 36.5% |
| caleyi | 44.6% | 42.6% | | 38.6% | 40.5% | 35.0% | 47.5% | 29.5% | 40.3% | 39.9% | 29.3% |
| camaldulensis | 38.7% | 41.5% | 40.6% | | 39.1% | 38.4% | 42.1% | 32.9% | 35.3% | 42.3% | 32.1% |
| cladocalyx | 41.3% | 44.0% | 43.8% | 40.3% | | 36.7% | 44.2% | 31.4% | 37.9% | 41.5% | 30.9% |
| cloeziana | 39.7% | 43.8% | 42.1% | 43.8% | 40.6% | | 44.1% | 40.5% | 36.5% | 43.6% | 37.2% |
| coolabah | 44.0% | 42.4% | 46.3% | 39.1% | 39.7% | 35.9% | | 31.0% | 40.1% | 40.2% | 30.4% |
| curtisii | 39.0% | 41.6% | 38.9% | 41.3% | 38.5% | 44.4% | 41.8% | | 34.7% | 41.3% | 37.8% |
| dawsonii | 32.3% | 30.7% | 33.7% | 28.0% | 29.2% | 25.7% | 34.4% | 22.1% | | 28.8% | 21.3% |
| decipiens | 38.1% | 41.9% | 40.1% | 40.3% | 38.4% | 36.5% | 41.3% | 31.3% | 34.7% | | 31.7% |
| erythrocorys | 32.9% | 36.1% | 33.1% | 34.7% | 32.5% | 35.3% | 35.2% | 32.8% | 28.7% | 35.8% | |
| fibrosa | 43.9% | 42.7% | 45.2% | 38.2% | 38.5% | 34.2% | 47.0% | 30.3% | 39.2% | 39.2% | 29.5% |
| globulus | 41.2% | 44.1% | 43.0% | 53.3% | 41.6% | 40.2% | 44.7% | 34.5% | 37.7% | 44.8% | 33.9% |
| grandis | 34.6% | 36.9% | 35.8% | 45.4% | 34.6% | 34.0% | 37.6% | 29.0% | 32.0% | 37.8% | 28.4% |
| guilfoylei | 38.0% | 41.2% | 40.5% | 40.8% | 38.7% | 39.8% | 41.8% | 36.2% | 34.9% | 41.9% | 35.3% |
| lansdowneana | 40.9% | 38.8% | 40.7% | 34.1% | 36.6% | 31.1% | 42.1% | 27.1% | 36.7% | 35.5% | 26.4% |
| leucophloia | 40.5% | 41.6% | 42.0% | 38.1% | 39.0% | 34.5% | 44.2% | 30.9% | 36.0% | 39.4% | 29.5% |
| marginata | 37.8% | 41.6% | 40.0% | 40.9% | 37.9% | 50.8% | 41.6% | 38.6% | 34.1% | 41.5% | 35.9% |
| melliodora | 45.5% | 41.5% | 46.2% | 38.1% | 39.0% | 34.5% | 46.9% | 30.0% | 40.2% | 39.3% | 29.1% |
| melliodora x sideroxylon | 40.7% | 40.7% | 40.7% | 40.7% | 40.7% | 40.7% | 40.7% | 40.7% | 40.7% | 40.7% | 40.7% |
| microcorys | 46.6% | 50.5% | 49.7% | 49.7% | 46.9% | 48.6% | 50.4% | 42.6% | 42.4% | 50.5% | 41.9% |
| ANBG9806169 | 35.1% | 38.0% | 36.5% | 38.1% | 35.0% | 48.9% | 38.4% | 36.4% | 31.4% | 38.0% | 33.6% |
| paniculata | 42.0% | 39.8% | 44.2% | 36.5% | 37.5% | 33.2% | 44.6% | 29.0% | 38.2% | 37.9% | 28.2% |
| pauciflora | 34.2% | 38.9% | 34.9% | 36.4% | 34.1% | 46.5% | 36.8% | 34.7% | 30.3% | 37.3% | 32.7% |
| polyanthemos | 43.1% | 40.8% | 45.4% | 36.6% | 38.1% | 32.9% | 45.0% | 29.1% | 45.1% | 38.2% | 28.3% |
| pumila | 37.2% | 40.0% | 38.3% | 47.9% | 37.8% | 36.6% | 40.2% | 30.9% | 34.4% | 40.4% | 30.5% |
| regnans | 34.8% | 39.2% | 37.2% | 37.8% | 35.8% | 46.7% | 38.2% | 36.6% | 31.8% | 37.9% | 34.1% |
| shirleyi | 40.8% | 39.7% | 44.2% | 35.8% | 37.3% | 33.0% | 44.8% | 28.5% | 37.2% | 37.4% | 28.1% |
| sideroxylon | 45.9% | 43.1% | 47.6% | 38.6% | 41.3% | 35.4% | 47.7% | 30.9% | 41.6% | 40.3% | 30.5% |
| tenuipes | 45.6% | 49.7% | 48.0% | 49.0% | 46.0% | 52.5% | 49.3% | 48.0% | 40.8% | 50.2% | 46.0% |
| victrix | 45.6% | 45.6% | 47.2% | 40.7% | 41.6% | 36.9% | 52.5% | 32.8% | 40.6% | 41.6% | 32.3% |
| viminalis | 39.9% | 42.5% | 41.5% | 51.4% | 39.9% | 39.1% | 43.4% | 33.8% | 36.6% | 43.2% | 32.8% |
| virginea | 41.4% | 45.5% | 43.4% | 43.7% | 41.0% | 38.5% | 44.7% | 34.1% | 37.2% | 48.0% | 32.6% |

| | ***fibrosa*** | ***globulus*** | ***grandis*** | ***guilfoylei*** | ***lansdowneana*** | ***leucophloia*** | ***marginata*** | ***melliodora*** | ***melliodora x sideroxylon*** | ***microcorys*** | ***ANBG9806169*** |
|---|---|---|---|---|---|---|---|---|---|---|---|
| ***albens*** | 42.7% | 37.0% | 34.9% | 30.5% | 42.6% | 37.6% | 32.5% | 47.5% | 43.7% | 34.9% | 30.1% |
| ***brandiana*** | 49.7% | 47.3% | 44.6% | 39.0% | 48.3% | 46.4% | 42.5% | 51.7% | 48.3% | 44.8% | 38.8% |
| ***caleyi*** | 45.4% | 39.8% | 37.4% | 33.4% | 43.7% | 40.3% | 35.5% | 49.9% | 45.3% | 38.2% | 32.3% |
| ***camaldulensis*** | 40.3% | 51.9% | 49.8% | 35.4% | 38.6% | 38.6% | 38.4% | 43.2% | 40.0% | 40.4% | 35.5% |
| ***cladocalyx*** | 41.9% | 41.8% | 39.2% | 34.6% | 42.7% | 40.7% | 36.5% | 45.6% | 42.7% | 39.2% | 33.6% |
| ***cloeziana*** | 41.1% | 45.1% | 42.7% | 39.4% | 40.0% | 39.9% | 54.1% | 44.4% | 40.9% | 45.2% | 52.1% |
| ***coolabah*** | 45.9% | 40.2% | 38.2% | 33.9% | 44.1% | 41.4% | 36.2% | 49.2% | 44.7% | 37.9% | 33.2% |
| ***curtisii*** | 39.9% | 42.3% | 39.9% | 39.4% | 38.0% | 38.9% | 45.3% | 42.4% | 40.7% | 43.5% | 42.2% |
| ***dawsonii*** | 32.8% | 29.2% | 27.9% | 24.2% | 33.0% | 28.9% | 25.1% | 36.2% | 33.7% | 27.4% | 23.3% |
| ***decipiens*** | 39.4% | 41.5% | 39.5% | 34.4% | 38.2% | 37.9% | 36.7% | 42.4% | 39.9% | 38.6% | 33.5% |
| ***erythrocorys*** | 33.3% | 36.0% | 33.7% | 33.3% | 32.0% | 32.4% | 36.3% | 35.3% | 33.5% | 36.7% | 33.3% |
| ***fibrosa*** | | 39.3% | 37.2% | 32.7% | 44.2% | 39.2% | 34.4% | 48.6% | 45.3% | 36.9% | 32.5% |
| ***globulus*** | 42.6% | | 53.7% | 37.2% | 41.1% | 41.1% | 40.5% | 45.4% | 42.1% | 42.5% | 37.0% |
| ***grandis*** | 35.7% | 47.7% | | 31.4% | 34.1% | 34.0% | 33.6% | 38.3% | 35.5% | 35.9% | 31.0% |
| ***guilfoylei*** | 39.9% | 41.9% | 39.9% | | 38.0% | 38.0% | 40.6% | 42.0% | 39.8% | 45.6% | 38.0% |
| ***lansdowneana*** | 41.4% | 35.5% | 33.1% | 29.4% | | 36.2% | 31.3% | 44.5% | 41.8% | 33.3% | 28.7% |
| ***leucophloia*** | 40.9% | 39.6% | 37.1% | 32.7% | 40.7% | | 35.0% | 44.7% | 41.6% | 37.1% | 31.9% |
| ***marginata*** | 39.0% | 42.4% | 39.8% | 37.8% | 37.8% | 38.0% | | 41.9% | 39.2% | 42.6% | 49.0% |
| ***melliodora*** | 45.2% | 39.0% | 37.1% | 32.3% | 44.5% | 39.9% | 34.7% | | 35.9% | 35.9% | 35.9% |
| ***melliodora x sideroxylon*** | 40.7% | 40.7% | 40.7% | 40.7% | 40.7% | 40.7% | 40.7% | 40.7% | | 40.7% | 40.7% |
| ***microcorys*** | 48.2% | 51.7% | 48.9% | 48.9% | 46.5% | 46.5% | 49.0% | 51.5% | 47.7% | | 45.0% |
| ***ANBG9806169*** | 37.2% | 39.1% | 36.8% | 35.6% | 35.1% | 34.8% | 49.2% | 38.9% | 36.5% | 39.5% | |
| ***paniculata*** | 44.5% | 37.6% | 35.5% | 30.4% | 41.9% | 38.4% | 33.0% | 46.8% | 43.2% | 34.7% | 30.1% |
| ***pauciflora*** | 37.6% | 37.6% | 35.3% | 35.7% | 34.5% | 33.9% | 47.0% | 37.7% | 36.1% | 38.7% | 50.7% |
| ***polyanthemos*** | 43.7% | 37.8% | 36.0% | 31.7% | 43.7% | 38.4% | 33.3% | 48.5% | 44.9% | 36.5% | 30.4% |
| ***pumila*** | 38.4% | 48.7% | 46.1% | 33.2% | 38.0% | 37.0% | 36.4% | 41.2% | 37.7% | 38.3% | 33.9% |
| ***regnans*** | 39.3% | 39.4% | 36.3% | 36.5% | 35.3% | 35.4% | 47.2% | 38.5% | 36.9% | 39.8% | 52.0% |
| ***shirleyi*** | 45.1% | 37.2% | 35.3% | 32.1% | 41.3% | 38.3% | 33.2% | 45.4% | 42.3% | 35.4% | 30.6% |
| ***sideroxylon*** | 46.8% | 40.1% | 37.7% | 33.9% | 46.6% | 40.8% | 35.8% | 53.1% | 53.4% | 38.3% | 32.6% |
| ***tenuipes*** | 49.1% | 50.7% | 47.6% | 45.9% | 44.8% | 45.0% | 53.1% | 50.6% | 47.7% | 51.2% | 49.3% |
| ***victrix*** | 50.6% | 41.9% | 39.4% | 36.2% | 46.2% | 42.6% | 37.5% | 50.2% | 46.1% | 40.1% | 34.5% |
| ***viminalis*** | 41.1% | 59.0% | 52.2% | 36.3% | 39.6% | 39.7% | 39.2% | 43.8% | 40.8% | 41.4% | 36.2% |
| ***virginea*** | 43.6% | 44.8% | 42.2% | 37.3% | 41.7% | 41.4% | 39.1% | 45.8% | 42.6% | 41.8% | 35.8% |

| | ***paniculata*** | ***pauciflora*** | ***polyanthemos*** | ***pumila*** | ***regnans*** | ***shirleyi*** | ***sideroxylon*** | ***tenuipes*** | ***victrix*** | ***viminalis*** | ***virginea*** |
|---|---|---|---|---|---|---|---|---|---|---|---|
| ***albens*** | 40.6% | 28.7% | 42.7% | 32.6% | 29.2% | 40.1% | 44.8% | 31.4% | 41.8% | 36.7% | 36.6% |
| ***brandiana*** | 46.1% | 38.6% | 48.2% | 41.9% | 39.0% | 46.6% | 50.2% | 40.3% | 49.8% | 46.5% | 47.9% |
| ***caleyi*** | 44.2% | 30.1% | 46.5% | 34.6% | 32.2% | 44.9% | 47.9% | 34.0% | 44.7% | 39.3% | 39.5% |
| ***camaldulensis*** | 38.3% | 33.1% | 39.3% | 45.6% | 34.5% | 38.2% | 40.9% | 36.6% | 40.4% | 51.2% | 41.9% |
| ***cladocalyx*** | 40.5% | 31.9% | 42.2% | 37.0% | 33.6% | 41.0% | 45.0% | 35.1% | 42.6% | 41.0% | 40.5% |
| ***cloeziana*** | 39.8% | 48.4% | 40.4% | 39.9% | 48.6% | 40.2% | 42.4% | 44.3% | 42.0% | 45.0% | 43.0% |
| ***coolabah*** | 43.3% | 31.1% | 44.8% | 35.4% | 32.5% | 44.3% | 46.7% | 34.2% | 48.4% | 40.0% | 39.7% |
| ***curtisii*** | 38.3% | 39.4% | 38.9% | 37.0% | 41.5% | 37.9% | 40.8% | 44.4% | 40.9% | 42.4% | 41.5% |
| ***dawsonii*** | 31.8% | 21.8% | 38.6% | 26.0% | 23.1% | 31.5% | 34.9% | 24.2% | 32.0% | 28.9% | 28.3% |
| ***decipiens*** | 38.1% | 32.2% | 39.1% | 36.6% | 32.8% | 38.2% | 40.6% | 35.4% | 39.3% | 40.9% | 43.6% |
| ***erythrocorys*** | 31.5% | 31.9% | 32.4% | 31.3% | 33.2% | 32.2% | 34.4% | 36.8% | 34.2% | 35.5% | 34.0% |
| ***fibrosa*** | 44.4% | 32.2% | 44.4% | 34.7% | 33.6% | 45.7% | 46.8% | 34.5% | 47.6% | 38.7% | 39.6% |
| ***globulus*** | 40.5% | 34.8% | 41.7% | 47.4% | 36.4% | 40.7% | 43.5% | 38.1% | 42.7% | 60.4% | 44.1% |
| ***grandis*** | 33.9% | 28.9% | 35.1% | 39.8% | 29.8% | 34.3% | 36.3% | 32.1% | 35.7% | 47.5% | 36.9% |
| ***guilfoylei*** | 36.7% | 37.3% | 39.2% | 36.5% | 38.0% | 39.3% | 40.9% | 39.1% | 41.3% | 41.8% | 41.7% |
| ***lansdowneana*** | 38.8% | 27.8% | 41.6% | 31.9% | 28.4% | 39.1% | 43.5% | 29.4% | 40.5% | 34.9% | 35.3% |
| ***leucophloia*** | 39.8% | 30.4% | 40.7% | 34.7% | 31.9% | 40.5% | 42.7% | 33.1% | 41.9% | 39.0% | 39.1% |
| ***marginata*** | 37.2% | 45.8% | 38.4% | 37.2% | 46.2% | 38.1% | 40.5% | 41.8% | 39.6% | 42.1% | 40.7% |
| ***melliodora*** | 35.9% | 35.9% | 35.9% | 35.9% | 35.9% | 35.9% | 35.9% | 35.9% | 35.9% | 35.9% | 35.9% |
| ***melliodora x sideroxylon*** | 40.7% | 40.7% | 40.7% | 40.7% | 40.7% | 40.7% | 40.7% | 40.7% | 40.7% | 40.7% | 40.7% |
| ***microcorys*** | 44.8% | 43.1% | 48.2% | 45.3% | 44.6% | 46.7% | 49.6% | 46.7% | 49.1% | 51.2% | 49.9% |
| ***ANBG9806169*** | 34.1% | 49.6% | 35.0% | 35.0% | 50.9% | 35.3% | 37.0% | 39.4% | 36.9% | 39.2% | 37.5% |
| ***paniculata*** | | 29.8% | 43.5% | 32.8% | 28.4% | 41.7% | 44.2% | 31.6% | 40.5% | 37.3% | 37.2% |
| ***pauciflora*** | 34.5% | | 35.9% | 33.3% | 54.0% | 34.9% | 36.7% | 38.5% | 38.4% | 37.0% | 37.2% |
| ***polyanthemos*** | 42.5% | 30.5% | | 33.5% | 30.6% | 41.9% | 46.5% | 32.1% | 43.2% | 37.7% | 37.9% |
| ***pumila*** | 36.4% | 31.4% | 37.9% | | 33.4% | 36.9% | 39.2% | 34.3% | 38.2% | 48.3% | 39.3% |
| ***regnans*** | 32.8% | 54.0% | 36.1% | 35.4% | | 36.7% | 37.6% | 40.9% | 38.7% | 38.6% | 38.2% |
| ***shirleyi*** | 41.1% | 29.5% | 42.2% | 32.8% | 31.1% | | 43.1% | 32.9% | 44.3% | 36.7% | 37.1% |
| ***sideroxylon*** | 43.8% | 31.8% | 47.3% | 35.4% | 32.5% | 43.6% | | 34.4% | 45.6% | 39.3% | 39.5% |
| ***tenuipes*** | 44.8% | 47.2% | 46.1% | 44.2% | 50.2% | 47.3% | 48.7% | | 48.9% | 50.1% | 48.0% |
| ***victrix*** | 42.8% | 35.1% | 46.8% | 36.7% | 35.6% | 47.6% | 48.6% | 36.9% | | 41.1% | 41.6% |
| ***viminalis*** | 39.3% | 33.6% | 40.6% | 46.1% | 34.9% | 39.5% | 41.7% | 36.9% | 41.0% | | 42.6% |
| ***virginea*** | 40.9% | 34.6% | 42.5% | 39.2% | 35.6% | 41.5% | 43.7% | 36.8% | 43.1% | 44.4% | |

**Supplementary Tables S10. Matrix of pairwise shared synteny.** Shows the proportion of the genome that is syntenic for all genome pairs. Read down the genome list (far left column) and extend across until the comparison genome is found. As genomes are different lengths the proportion that species X has syntenic to species Y is different to the proportion that species Y has syntenic to species X.

| | albens | brandiana | caleyi | camaldulensis | cladocalyx | cloeziana | coolabah | curtisii | dawsonii | decipiens | erythrocorys |
|---|---|---|---|---|---|---|---|---|---|---|---|
| albens | | 35.2% | 39.5% | 34.5% | 40.0% | 22.9% | 36.3% | 26.8% | 45.8% | 34.5% | 20.9% |
| brandiana | 33.1% | | 31.6% | 28.2% | 32.3% | 18.5% | 28.3% | 24.9% | 38.8% | 29.1% | 19.1% |
| caleyi | 37.9% | 32.1% | | 33.4% | 36.6% | 21.4% | 34.1% | 28.0% | 43.5% | 33.5% | 22.6% |
| camaldulensis | 32.6% | 27.6% | 31.3% | | 31.3% | 18.3% | 27.1% | 24.1% | 36.9% | 27.7% | 20.4% |
| cladocalyx | 39.2% | 33.1% | 36.7% | 32.0% | | 21.4% | 33.9% | 27.9% | 43.1% | 33.6% | 22.6% |
| cloeziana | 23.2% | 18.4% | 21.7% | 18.4% | 21.3% | | 18.3% | 27.4% | 26.1% | 19.9% | 20.6% |
| coolabah | 36.6% | 30.6% | 35.5% | 29.1% | 34.9% | 18.7% | | 23.7% | 40.9% | 30.2% | 19.5% |
| curtisii | 28.2% | 25.4% | 29.6% | 25.7% | 28.8% | 27.7% | 24.6% | | 32.0% | 26.9% | 24.5% |
| dawsonii | 48.3% | 42.0% | 46.9% | 40.2% | 45.1% | 27.5% | 43.4% | 30.3% | | 40.8% | 25.3% |
| decipiens | 34.2% | 29.9% | 32.3% | 29.0% | 32.7% | 20.2% | 28.7% | 26.1% | 37.6% | | 18.6% |
| erythrocorys | 22.6% | 19.2% | 22.7% | 20.2% | 22.1% | 20.2% | 18.9% | 22.2% | 24.3% | 18.7% | |
| fibrosa | 40.3% | 33.6% | 40.1% | 34.6% | 39.9% | 22.8% | 36.0% | 27.4% | 44.9% | 35.1% | 21.8% |
| globulus | 30.9% | 26.3% | 30.0% | 28.6% | 29.2% | 17.6% | 25.1% | 23.6% | 34.8% | 26.7% | 18.3% |
| grandis | 36.1% | 31.6% | 35.8% | 35.2% | 34.8% | 21.6% | 30.8% | 26.3% | 38.3% | 32.4% | 21.0% |
| guilfoylei | 32.4% | 30.1% | 30.8% | 29.4% | 31.5% | 27.8% | 27.6% | 32.5% | 35.5% | 30.1% | 26.4% |
| lansdowneana | 43.4% | 36.6% | 42.9% | 36.9% | 40.6% | 24.3% | 39.2% | 28.4% | 47.0% | 37.5% | 23.7% |
| leucophloia | 39.8% | 34.9% | 38.9% | 33.5% | 37.3% | 23.2% | 34.9% | 27.1% | 43.3% | 33.3% | 23.1% |
| marginata | 23.8% | 19.6% | 22.7% | 20.2% | 23.2% | 30.4% | 19.3% | 28.1% | 27.2% | 20.3% | 20.6% |
| melliodora | 37.9% | 32.3% | 35.5% | 31.1% | 36.0% | 18.9% | 32.4% | 24.6% | 42.6% | 31.7% | 20.0% |
| melliodora x sideroxylon | 22.9% | 22.9% | 22.9% | 22.9% | 22.9% | 22.9% | 22.9% | 22.9% | 22.9% | 22.9% | 22.9% |
| microcorys | 25.9% | 21.5% | 22.8% | 21.6% | 25.0% | 19.4% | 20.9% | 27.4% | 30.2% | 23.1% | 21.6% |
| ANBG9806169 | 25.2% | 22.7% | 25.4% | 23.0% | 25.2% | 35.0% | 21.7% | 29.9% | 28.1% | 23.2% | 23.4% |
| paniculata | 42.2% | 36.9% | 41.4% | 36.3% | 41.6% | 25.0% | 37.8% | 29.2% | 46.0% | 35.8% | 22.9% |
| pauciflora | 28.4% | 22.3% | 28.5% | 26.2% | 28.3% | 40.3% | 24.8% | 33.1% | 31.7% | 26.0% | 24.9% |
| polyanthemos | 40.1% | 34.2% | 37.8% | 34.2% | 39.1% | 22.9% | 35.9% | 26.6% | 42.7% | 33.8% | 22.3% |
| pumila | 36.4% | 32.1% | 35.8% | 35.1% | 35.1% | 22.7% | 31.0% | 28.5% | 39.0% | 33.2% | 23.6% |
| regnans | 26.9% | 22.9% | 24.9% | 23.4% | 24.6% | 39.2% | 22.1% | 29.9% | 28.6% | 24.5% | 21.9% |
| shirleyi | 42.8% | 35.8% | 40.4% | 35.8% | 39.9% | 23.6% | 37.7% | 29.1% | 45.7% | 35.3% | 24.4% |
| sideroxylon | 38.1% | 31.7% | 35.5% | 32.4% | 34.8% | 20.3% | 33.2% | 24.7% | 42.2% | 32.0% | 19.6% |
| tenuipes | 24.3% | 19.0% | 22.7% | 20.4% | 23.0% | 22.1% | 19.0% | 30.4% | 27.7% | 19.7% | 19.7% |
| victrix | 37.1% | 29.9% | 36.2% | 31.1% | 35.8% | 20.3% | 31.8% | 24.6% | 42.5% | 31.7% | 19.7% |
| viminalis | 31.0% | 26.8% | 30.2% | 29.1% | 29.7% | 17.4% | 25.9% | 23.1% | 34.4% | 27.1% | 19.1% |
| virginea | 34.0% | 28.6% | 32.7% | 29.4% | 33.4% | 21.9% | 28.5% | 27.3% | 38.8% | 31.5% | 23.1% |

| | fibrosa | globulus | grandis | guilfoylei | lansdowneana | leucophloia | marginata | melliodora | melliodora x sideroxylon | microcorys | ANBG9806169 |
|---|---|---|---|---|---|---|---|---|---|---|---|
| albens | 41.0% | 32.5% | 37.0% | 32.4% | 41.4% | 41.0% | 23.1% | 36.1% | 41.0% | 27.4% | 23.7% |
| brandiana | 31.5% | 25.7% | 30.5% | 30.3% | 32.5% | 34.5% | 18.8% | 27.7% | 33.2% | 22.6% | 21.8% |
| caleyi | 39.3% | 31.5% | 36.0% | 31.3% | 39.8% | 39.8% | 22.5% | 32.8% | 38.5% | 25.0% | 24.1% |
| camaldulensis | 32.8% | 28.7% | 33.7% | 29.1% | 33.1% | 32.8% | 19.2% | 27.4% | 33.3% | 22.9% | 21.6% |
| cladocalyx | 39.9% | 30.3% | 35.2% | 32.3% | 38.2% | 38.2% | 22.8% | 34.0% | 39.2% | 26.5% | 23.9% |
| cloeziana | 22.4% | 17.0% | 21.1% | 27.7% | 22.6% | 23.4% | 28.6% | 17.2% | 24.2% | 19.9% | 32.2% |
| coolabah | 37.1% | 27.8% | 32.2% | 28.4% | 37.7% | 36.9% | 19.5% | 31.3% | 37.8% | 23.7% | 21.2% |
| curtisii | 28.2% | 24.5% | 28.4% | 33.3% | 29.0% | 28.7% | 27.9% | 24.0% | 28.8% | 27.5% | 29.4% |
| dawsonii | 48.6% | 38.6% | 41.7% | 36.5% | 48.7% | 47.0% | 28.7% | 45.3% | 48.5% | 32.6% | 29.1% |
| decipiens | 35.0% | 28.4% | 32.4% | 30.9% | 35.1% | 33.7% | 20.5% | 29.8% | 33.8% | 25.7% | 23.1% |
| erythrocorys | 22.2% | 18.7% | 21.9% | 25.1% | 23.0% | 22.3% | 19.0% | 18.7% | 23.2% | 21.1% | 22.1% |
| fibrosa | | 32.7% | 37.3% | 33.9% | 39.6% | 41.9% | 23.7% | 35.4% | 40.0% | 28.3% | 23.6% |
| globulus | 31.0% | | 31.3% | 29.0% | 32.0% | 30.1% | 17.7% | 25.8% | 32.6% | 22.2% | 20.6% |
| grandis | 36.2% | 33.0% | | 32.0% | 37.2% | 36.0% | 23.0% | 31.8% | 37.4% | 27.4% | 24.9% |
| guilfoylei | 32.2% | 28.5% | 31.8% | | 33.1% | 33.6% | 27.7% | 28.4% | 32.9% | 28.5% | 28.0% |
| lansdowneana | 42.3% | 35.0% | 39.2% | 34.4% | | 43.2% | 25.2% | 39.8% | 43.8% | 30.1% | 26.1% |
| leucophloia | 40.8% | 30.8% | 35.9% | 33.2% | 39.5% | | 23.2% | 34.7% | 39.5% | 27.7% | 25.6% |
| marginata | 23.4% | 18.3% | 23.1% | 28.8% | 24.0% | 23.9% | | 18.9% | 24.9% | 21.4% | 34.1% |
| melliodora | 37.2% | 29.9% | 34.0% | 29.9% | 38.2% | 36.7% | 20.1% | | 39.6% | 39.6% | 39.6% |
| melliodora x sideroxylon | 22.9% | 22.9% | 22.9% | 22.9% | 22.9% | 22.9% | 22.9% | 22.9% | | 22.9% | 22.9% |
| microcorys | 25.6% | 19.8% | 24.8% | 27.9% | 26.5% | 26.9% | 19.6% | 19.4% | 26.4% | | 23.1% |
| ANBG9806169 | 24.0% | 21.3% | 25.9% | 28.6% | 25.4% | 26.4% | 34.7% | 21.1% | 26.7% | 24.0% | |
| paniculata | 41.7% | 34.5% | 38.2% | 35.0% | 42.2% | 41.0% | 25.8% | 37.0% | 42.3% | 30.6% | 26.8% |
| pauciflora | 25.7% | 24.8% | 28.0% | 30.9% | 27.9% | 27.8% | 39.2% | 22.7% | 27.2% | 26.8% | 41.1% |
| polyanthemos | 40.3% | 32.4% | 36.2% | 31.9% | 39.7% | 39.7% | 23.0% | 35.1% | 39.5% | 26.1% | 25.1% |
| pumila | 36.1% | 33.0% | 37.5% | 35.0% | 36.2% | 36.0% | 23.5% | 31.7% | 38.2% | 28.3% | 24.2% |
| regnans | 23.1% | 21.1% | 25.9% | 30.4% | 26.3% | 26.3% | 38.3% | 21.4% | 26.5% | 24.9% | 39.3% |
| shirleyi | 39.6% | 33.7% | 37.8% | 32.7% | 41.8% | 41.4% | 24.6% | 38.0% | 41.9% | 29.7% | 26.1% |
| sideroxylon | 37.3% | 29.4% | 33.9% | 29.9% | 37.4% | 37.7% | 20.9% | 31.7% | 34.8% | 24.1% | 22.9% |
| tenuipes | 21.7% | 18.7% | 23.2% | 29.2% | 24.8% | 25.2% | 21.7% | 17.0% | 22.6% | 21.5% | 24.2% |
| victrix | 32.9% | 29.4% | 33.8% | 29.4% | 36.5% | 38.1% | 20.4% | 32.0% | 37.6% | 23.9% | 22.7% |
| viminalis | 32.3% | 26.5% | 31.5% | 28.4% | 31.8% | 30.4% | 17.8% | 26.1% | 32.5% | 22.0% | 20.5% |
| virginea | 33.1% | 28.3% | 32.9% | 31.6% | 34.2% | 33.7% | 22.2% | 28.8% | 34.8% | 26.5% | 24.6% |

| | paniculata | pauciflora | polyanthemos | pumila | regnans | shirleyi | sideroxylon | tenuipes | victrix | viminalis | virginea |
|---|---|---|---|---|---|---|---|---|---|---|---|
| albens | 43.3% | 27.8% | 40.3% | 37.4% | 24.2% | 42.7% | 38.4% | 24.1% | 38.6% | 32.7% | 36.1% |
| brandiana | 36.0% | 20.3% | 31.5% | 32.4% | 20.4% | 34.4% | 29.2% | 20.3% | 28.8% | 27.1% | 29.9% |
| caleyi | 41.8% | 26.9% | 36.3% | 37.0% | 22.6% | 39.8% | 34.7% | 23.7% | 36.8% | 31.4% | 34.7% |
| camaldulensis | 34.8% | 23.6% | 31.9% | 35.4% | 20.6% | 34.5% | 29.9% | 21.8% | 30.0% | 29.0% | 30.2% |
| cladocalyx | 41.3% | 25.8% | 38.4% | 36.3% | 23.0% | 39.7% | 34.4% | 23.3% | 36.0% | 30.8% | 35.4% |
| cloeziana | 24.3% | 35.9% | 22.2% | 22.5% | 35.8% | 23.0% | 20.1% | 23.4% | 20.0% | 17.3% | 21.9% |
| coolabah | 39.2% | 24.0% | 36.1% | 32.8% | 20.4% | 38.4% | 33.5% | 19.8% | 34.0% | 27.7% | 31.4% |
| curtisii | 30.6% | 32.7% | 27.9% | 30.3% | 29.6% | 30.5% | 26.1% | 31.7% | 25.9% | 24.3% | 28.0% |
| dawsonii | 49.4% | 31.9% | 46.9% | 41.6% | 28.1% | 48.6% | 46.3% | 28.7% | 46.4% | 38.2% | 42.3% |
| decipiens | 35.2% | 23.8% | 32.4% | 34.3% | 22.6% | 34.1% | 31.1% | 20.8% | 31.9% | 29.2% | 34.6% |
| erythrocorys | 23.5% | 23.0% | 22.0% | 23.2% | 19.5% | 22.2% | 19.9% | 19.5% | 20.1% | 18.1% | 22.2% |
| fibrosa | 42.1% | 24.8% | 39.8% | 37.8% | 21.7% | 39.1% | 37.0% | 23.2% | 34.2% | 33.8% | 35.6% |
| globulus | 33.5% | 23.5% | 30.3% | 34.1% | 19.6% | 32.4% | 28.0% | 19.6% | 28.7% | 26.0% | 29.3% |
| grandis | 38.3% | 26.2% | 35.3% | 39.5% | 23.8% | 37.6% | 33.8% | 23.6% | 34.6% | 33.7% | 34.9% |
| guilfoylei | 35.6% | 29.4% | 31.5% | 34.8% | 28.5% | 32.0% | 29.3% | 29.2% | 29.1% | 28.1% | 30.8% |
| lansdowneana | 45.6% | 27.7% | 42.0% | 39.2% | 25.5% | 44.6% | 40.8% | 26.5% | 40.6% | 35.6% | 38.7% |
| leucophloia | 41.0% | 26.7% | 38.6% | 36.7% | 24.3% | 40.9% | 36.3% | 26.0% | 38.5% | 31.5% | 35.4% |
| marginata | 26.4% | 37.5% | 23.1% | 24.5% | 36.6% | 24.4% | 21.3% | 24.4% | 21.2% | 18.9% | 22.9% |
| melliodora | 39.6% | 39.6% | 39.6% | 39.6% | 39.6% | 39.6% | 39.6% | 39.6% | 39.6% | 39.6% | 39.6% |
| melliodora x sideroxylon | 22.9% | 22.9% | 22.9% | 22.9% | 22.9% | 22.9% | 22.9% | 22.9% | 22.9% | 22.9% | 22.9% |
| microcorys | 29.8% | 24.3% | 24.1% | 27.4% | 22.6% | 27.0% | 22.2% | 22.7% | 22.7% | 20.0% | 24.7% |
| ANBG9806169 | 27.5% | 39.3% | 25.4% | 25.1% | 38.4% | 26.0% | 23.6% | 26.9% | 23.3% | 21.3% | 25.0% |
| paniculata | | 26.5% | 40.3% | 39.5% | 30.6% | 43.3% | 39.9% | 25.6% | 42.8% | 34.7% | 38.6% |
| pauciflora | 27.2% | | 24.6% | 29.4% | 37.7% | 28.4% | 25.8% | 28.2% | 23.5% | 25.6% | 28.6% |
| polyanthemos | 41.4% | 23.1% | | 37.5% | 23.3% | 41.0% | 37.0% | 24.3% | 37.8% | 32.4% | 34.9% |
| pumila | 38.5% | 27.8% | 36.2% | | 24.5% | 37.7% | 33.9% | 25.9% | 34.5% | 33.7% | 35.9% |
| regnans | 31.8% | 36.9% | 24.6% | 25.9% | | 26.5% | 23.4% | 25.3% | 22.2% | 22.1% | 25.8% |
| shirleyi | 44.2% | 27.7% | 40.6% | 39.0% | 25.2% | | 40.2% | 25.0% | 37.8% | 34.6% | 37.2% |
| sideroxylon | 40.4% | 23.2% | 36.0% | 34.9% | 21.3% | 39.9% | | 21.4% | 35.4% | 30.4% | 33.7% |
| tenuipes | 25.6% | 25.4% | 22.9% | 25.8% | 22.3% | 23.3% | 20.4% | | 19.7% | 18.7% | 23.5% |
| victrix | 41.7% | 21.8% | 35.2% | 35.0% | 20.4% | 35.8% | 33.5% | 20.6% | | 30.2% | 32.4% |
| viminalis | 33.4% | 23.5% | 30.0% | 34.2% | 19.9% | 32.2% | 28.6% | 19.7% | 28.8% | | 29.3% |
| virginea | 36.2% | 26.0% | 32.2% | 35.5% | 23.5% | 34.7% | 31.0% | 24.7% | 31.1% | 28.1% | |

**Supplementary Tables S11. Matrix of pairwise rearrangements.** Shows the proportion of the genome that is rearranged for all genome pairs. Read down the genome list (far left column) and extend across until the comparison genome is found. As genomes are different lengths the proportion that species X has rearranged to species Y is different to the proportion that species Y has rearranged to species X.

| | albens | brandiana | caleyi | camaldulensis | cladocalyx | cloeziana | coolabah | curtisii | dawsonii | decipiens | erythrocorys |
|---|---|---|---|---|---|---|---|---|---|---|---|
| albens | | 74.4% | 82.6% | 70.3% | 76.8% | 54.9% | 80.1% | 55.6% | 83.3% | 71.4% | 49.3% |
| brandiana | 79.9% | | 80.7% | 73.8% | 79.1% | 60.4% | 78.6% | 61.5% | 81.3% | 77.3% | 55.6% |
| caleyi | 82.5% | 74.7% | | 71.9% | 77.1% | 56.4% | 81.6% | 57.6% | 83.8% | 73.4% | 51.9% |
| camaldulensis | 71.4% | 69.0% | 71.9% | | 70.3% | 56.7% | 69.2% | 57.0% | 72.1% | 70.0% | 52.5% |
| cladocalyx | 80.5% | 77.1% | 80.5% | 72.3% | | 58.1% | 78.1% | 59.3% | 80.9% | 75.0% | 53.4% |
| cloeziana | 63.0% | 62.2% | 63.7% | 62.2% | 61.9% | | 62.4% | 68.0% | 62.6% | 63.5% | 57.8% |
| coolabah | 80.6% | 73.0% | 81.8% | 68.2% | 74.6% | 54.6% | | 54.7% | 81.1% | 70.3% | 49.9% |
| curtisii | 67.2% | 67.0% | 68.5% | 67.0% | 67.3% | 72.1% | 66.4% | | 66.7% | 68.2% | 62.4% |
| dawsonii | 80.6% | 72.7% | 80.7% | 68.3% | 74.2% | 53.2% | 77.8% | 52.4% | | 69.6% | 46.7% |
| decipiens | 72.3% | 71.8% | 72.4% | 69.3% | 71.1% | 56.7% | 70.0% | 57.4% | 72.3% | | 50.4% |
| erythrocorys | 55.4% | 55.3% | 55.8% | 54.9% | 54.5% | 55.4% | 54.2% | 55.0% | 53.0% | 54.5% | |
| fibrosa | 84.2% | 76.3% | 85.3% | 72.8% | 78.4% | 57.0% | 83.0% | 57.7% | 84.1% | 74.3% | 51.4% |
| globulus | 72.1% | 70.4% | 73.0% | 81.9% | 70.9% | 57.8% | 69.8% | 58.1% | 72.4% | 71.6% | 52.2% |
| grandis | 70.7% | 68.5% | 71.7% | 80.6% | 69.4% | 55.6% | 68.4% | 55.4% | 70.3% | 70.3% | 49.5% |
| guilfoylei | 70.4% | 71.3% | 71.4% | 70.2% | 70.1% | 67.6% | 69.4% | 68.7% | 70.3% | 72.0% | 61.7% |
| lansdowneana | 84.3% | 75.5% | 83.6% | 71.0% | 77.2% | 55.4% | 81.3% | 55.5% | 83.7% | 73.0% | 50.1% |
| leucophloia | 80.3% | 76.5% | 80.8% | 71.6% | 76.2% | 57.7% | 79.1% | 58.0% | 79.4% | 72.7% | 52.7% |
| marginata | 61.6% | 61.2% | 62.7% | 61.1% | 61.1% | 81.2% | 60.9% | 66.7% | 61.2% | 61.9% | 56.5% |
| melliodora | 83.4% | 73.8% | 81.7% | 69.2% | 74.9% | 53.4% | 79.3% | 54.6% | 82.7% | 70.9% | 49.1% |
| melliodora x sideroxylon | 63.5% | 63.5% | 63.5% | 63.5% | 63.5% | 63.5% | 63.5% | 63.5% | 63.5% | 63.5% | 63.5% |
| microcorys | 72.5% | 72.0% | 72.5% | 71.3% | 72.0% | 68.0% | 71.3% | 70.0% | 72.7% | 73.6% | 63.5% |
| ANBG9806169 | 60.3% | 60.7% | 61.9% | 61.1% | 60.2% | 84.0% | 60.1% | 66.3% | 59.5% | 61.2% | 56.9% |
| paniculata | 84.1% | 76.7% | 85.6% | 72.8% | 79.1% | 58.2% | 82.3% | 58.2% | 84.3% | 73.7% | 51.1% |
| pauciflora | 62.6% | 61.2% | 63.4% | 62.7% | 62.4% | 86.7% | 61.6% | 67.8% | 62.0% | 63.3% | 57.5% |
| polyanthemos | 83.2% | 75.0% | 83.3% | 70.8% | 77.2% | 55.7% | 81.0% | 55.7% | 87.8% | 72.0% | 50.5% |
| pumila | 73.6% | 72.1% | 74.1% | 83.1% | 72.9% | 59.3% | 71.2% | 59.4% | 73.4% | 73.6% | 54.1% |
| regnans | 61.7% | 62.1% | 62.2% | 61.2% | 60.4% | 85.9% | 60.2% | 66.5% | 60.4% | 62.4% | 56.0% |
| shirleyi | 83.6% | 75.5% | 84.6% | 71.6% | 77.1% | 56.6% | 82.5% | 57.6% | 82.9% | 72.7% | 52.5% |
| sideroxylon | 84.0% | 74.8% | 83.0% | 71.0% | 76.1% | 55.7% | 80.9% | 55.7% | 83.9% | 72.2% | 50.1% |
| tenuipes | 70.0% | 68.7% | 70.7% | 69.4% | 68.9% | 74.6% | 68.3% | 78.4% | 68.5% | 69.9% | 65.7% |
| victrix | 82.7% | 75.5% | 83.5% | 71.7% | 77.4% | 57.1% | 84.3% | 57.5% | 83.1% | 73.3% | 52.0% |
| viminalis | 70.9% | 69.4% | 71.7% | 80.4% | 69.6% | 56.5% | 69.3% | 56.8% | 71.0% | 70.3% | 51.9% |
| virginea | 75.3% | 74.2% | 76.1% | 73.1% | 74.4% | 60.4% | 73.2% | 61.4% | 76.0% | 79.5% | 55.7% |

| | ***fibrosa*** | ***globulus*** | ***grandis*** | ***guilfoylei*** | ***lansdowneana*** | ***leucophloia*** | ***marginata*** | ***melliodora*** | ***melliodora x sideroxylon*** | ***microcorys*** | ***ANBG9806169*** |
|---|---|---|---|---|---|---|---|---|---|---|---|
| ***albens*** | 83.7% | 69.6% | 72.0% | 63.0% | 84.0% | 78.6% | 55.6% | 83.6% | 84.7% | 62.3% | 53.8% |
| ***brandiana*** | 81.2% | 73.0% | 75.1% | 69.3% | 80.8% | 80.9% | 61.3% | 79.4% | 81.6% | 67.4% | 60.6% |
| ***caleyi*** | 84.8% | 71.4% | 73.5% | 64.6% | 83.5% | 80.0% | 57.9% | 82.7% | 83.8% | 63.3% | 56.3% |
| ***camaldulensis*** | 73.2% | 80.6% | 83.5% | 64.6% | 71.7% | 71.4% | 57.6% | 70.6% | 73.3% | 63.2% | 57.1% |
| ***cladocalyx*** | 81.8% | 72.1% | 74.4% | 66.9% | 80.9% | 78.9% | 59.3% | 79.6% | 81.8% | 65.7% | 57.5% |
| ***cloeziana*** | 63.4% | 62.1% | 63.8% | 67.1% | 62.6% | 63.3% | 82.7% | 61.5% | 65.1% | 65.1% | 84.3% |
| ***coolabah*** | 83.0% | 68.0% | 70.4% | 62.3% | 81.7% | 78.2% | 55.7% | 80.6% | 82.4% | 61.6% | 54.5% |
| ***curtisii*** | 68.1% | 66.9% | 68.3% | 72.7% | 67.0% | 67.6% | 73.2% | 66.4% | 69.4% | 71.0% | 71.7% |
| ***dawsonii*** | 81.5% | 67.8% | 69.5% | 60.7% | 81.7% | 75.8% | 53.8% | 81.5% | 82.2% | 60.0% | 52.4% |
| ***decipiens*** | 74.4% | 70.0% | 71.8% | 65.3% | 73.3% | 71.6% | 57.2% | 72.2% | 73.7% | 64.3% | 56.7% |
| ***erythrocorys*** | 55.5% | 54.7% | 55.5% | 58.5% | 55.0% | 54.8% | 55.4% | 54.0% | 56.7% | 57.8% | 55.4% |
| ***fibrosa*** | | 72.0% | 74.5% | 66.6% | 83.8% | 81.1% | 58.1% | 84.0% | 85.3% | 65.1% | 56.1% |
| ***globulus*** | 73.6% | | 85.0% | 66.2% | 73.1% | 71.2% | 58.2% | 71.2% | 74.7% | 64.7% | 57.6% |
| ***grandis*** | 71.9% | 80.7% | | 63.4% | 71.2% | 70.1% | 56.6% | 70.1% | 72.9% | 63.3% | 55.8% |
| ***guilfoylei*** | 72.1% | 70.4% | 71.7% | | 71.1% | 71.5% | 68.3% | 70.4% | 72.7% | 74.1% | 66.0% |
| ***lansdowneana*** | 83.7% | 70.4% | 72.4% | 63.8% | | 79.5% | 56.5% | 84.3% | 85.7% | 63.4% | 54.8% |
| ***leucophloia*** | 81.8% | 70.4% | 72.9% | 65.8% | 80.2% | | 58.2% | 79.4% | 81.1% | 64.8% | 57.5% |
| ***marginata*** | 62.4% | 60.7% | 62.8% | 66.7% | 61.8% | 61.9% | | 60.8% | 64.1% | 63.9% | 83.1% |
| ***melliodora*** | 82.4% | 69.0% | 71.2% | 62.2% | 82.7% | 76.7% | 54.8% | | 75.5% | 75.5% | 75.5% |
| ***melliodora x sideroxylon*** | 63.5% | 63.5% | 63.5% | 63.5% | 63.5% | 63.5% | 63.5% | 63.5% | | 63.5% | 63.5% |
| ***microcorys*** | 73.7% | 71.5% | 73.8% | 76.9% | 73.0% | 73.4% | 68.6% | 70.9% | 74.1% | | 68.1% |
| ***ANBG9806169*** | 61.2% | 60.5% | 62.7% | 64.2% | 60.4% | 61.2% | 83.9% | 59.9% | 63.1% | 63.5% | |
| ***paniculata*** | 86.2% | 72.1% | 73.8% | 65.4% | 84.0% | 79.4% | 58.8% | 83.9% | 85.5% | 65.2% | 56.9% |
| ***pauciflora*** | 63.3% | 62.4% | 63.3% | 66.6% | 62.4% | 61.7% | 86.2% | 60.4% | 63.3% | 65.5% | 91.9% |
| ***polyanthemos*** | 84.0% | 70.2% | 72.2% | 63.6% | 83.4% | 78.1% | 56.3% | 83.5% | 84.4% | 62.6% | 55.5% |
| ***pumila*** | 74.5% | 81.6% | 83.6% | 68.2% | 74.3% | 73.0% | 59.9% | 72.9% | 75.9% | 66.6% | 58.1% |
| ***regnans*** | 62.4% | 60.6% | 62.3% | 66.9% | 61.7% | 61.8% | 85.5% | 59.9% | 63.4% | 64.7% | 91.3% |
| ***shirleyi*** | 84.7% | 70.9% | 73.1% | 64.7% | 83.1% | 79.7% | 57.8% | 83.3% | 84.2% | 65.1% | 56.7% |
| ***sideroxylon*** | 84.1% | 69.6% | 71.6% | 63.8% | 84.0% | 78.5% | 56.7% | 84.8% | 88.2% | 62.4% | 55.6% |
| ***tenuipes*** | 70.8% | 69.3% | 70.8% | 75.1% | 69.5% | 70.2% | 74.8% | 67.6% | 70.3% | 72.7% | 73.5% |
| ***victrix*** | 83.5% | 71.3% | 73.2% | 65.6% | 82.7% | 80.7% | 57.9% | 82.2% | 83.7% | 64.0% | 57.2% |
| ***viminalis*** | 73.3% | 85.5% | 83.7% | 64.7% | 71.4% | 70.1% | 57.0% | 70.0% | 73.3% | 63.3% | 56.7% |
| ***virginea*** | 76.7% | 73.1% | 75.1% | 68.9% | 75.9% | 75.1% | 61.3% | 74.5% | 77.4% | 68.3% | 60.4% |

| | paniculata | pauciflora | polyanthemos | pumila | regnans | shirleyi | sideroxylon | tenuipes | victrix | viminalis | virginea |
|---|---|---|---|---|---|---|---|---|---|---|---|
| ***albens*** | 84.0% | 56.5% | 83.0% | 69.9% | 53.4% | 82.8% | 83.2% | 55.5% | 80.4% | 69.3% | 72.6% |
| ***brandiana*** | 82.1% | 58.8% | 79.7% | 74.3% | 59.4% | 81.0% | 79.4% | 60.6% | 78.6% | 73.6% | 77.7% |
| ***caleyi*** | 86.0% | 57.0% | 82.7% | 71.7% | 54.8% | 84.7% | 82.6% | 57.7% | 81.5% | 70.7% | 74.1% |
| ***camaldulensis*** | 73.0% | 56.7% | 71.2% | 81.0% | 55.2% | 72.7% | 70.8% | 58.4% | 70.5% | 80.3% | 72.1% |
| ***cladocalyx*** | 81.8% | 57.7% | 80.5% | 73.2% | 56.5% | 80.6% | 79.3% | 58.4% | 78.6% | 71.8% | 75.9% |
| ***cloeziana*** | 64.1% | 84.3% | 62.6% | 62.4% | 84.5% | 63.2% | 62.6% | 67.7% | 62.0% | 62.3% | 64.9% |
| ***coolabah*** | 82.5% | 55.2% | 80.9% | 68.2% | 52.9% | 82.8% | 80.2% | 54.0% | 82.4% | 67.7% | 71.1% |
| ***curtisii*** | 68.8% | 72.1% | 66.8% | 67.3% | 71.1% | 68.5% | 66.9% | 76.1% | 66.8% | 66.6% | 69.5% |
| ***dawsonii*** | 81.3% | 53.7% | 85.5% | 67.6% | 51.1% | 80.2% | 81.3% | 52.9% | 78.3% | 67.0% | 70.7% |
| ***decipiens*** | 73.2% | 56.1% | 71.5% | 70.9% | 55.4% | 72.2% | 71.7% | 56.2% | 71.2% | 70.1% | 78.2% |
| ***erythrocorys*** | 55.0% | 54.9% | 54.5% | 54.5% | 52.7% | 54.5% | 54.3% | 56.3% | 54.3% | 53.6% | 56.3% |
| ***fibrosa*** | 86.5% | 57.0% | 84.2% | 72.5% | 55.4% | 84.9% | 83.8% | 57.7% | 81.8% | 72.4% | 75.2% |
| ***globulus*** | 74.1% | 58.3% | 72.0% | 81.5% | 56.0% | 73.1% | 71.4% | 57.7% | 71.3% | 86.5% | 73.4% |
| ***grandis*** | 72.3% | 55.1% | 70.3% | 79.3% | 53.6% | 71.9% | 70.2% | 55.7% | 70.2% | 81.1% | 71.8% |
| ***guilfoylei*** | 72.4% | 66.7% | 70.7% | 71.3% | 66.6% | 71.4% | 70.2% | 68.3% | 70.4% | 69.9% | 72.5% |
| ***lansdowneana*** | 84.4% | 55.5% | 83.6% | 71.1% | 54.0% | 83.7% | 84.3% | 55.9% | 81.1% | 70.5% | 74.0% |
| ***leucophloia*** | 80.8% | 57.1% | 79.4% | 71.4% | 56.2% | 81.5% | 79.0% | 59.2% | 80.4% | 70.5% | 74.5% |
| ***marginata*** | 63.6% | 83.3% | 61.5% | 61.7% | 82.8% | 62.5% | 61.9% | 66.3% | 60.8% | 61.0% | 63.5% |
| ***melliodora*** | 75.5% | 75.5% | 75.5% | 75.5% | 75.5% | 75.5% | 75.5% | 75.5% | 75.5% | 75.5% | 75.5% |
| ***melliodora x sideroxylon*** | 63.5% | 63.5% | 63.5% | 63.5% | 63.5% | 63.5% | 63.5% | 63.5% | 63.5% | 63.5% | 63.5% |
| ***microcorys*** | 74.6% | 67.4% | 72.3% | 72.8% | 67.2% | 73.7% | 71.8% | 69.4% | 71.7% | 71.2% | 74.6% |
| ***ANBG9806169*** | 61.5% | 88.9% | 60.4% | 60.0% | 89.3% | 61.3% | 60.6% | 66.3% | 60.2% | 60.4% | 62.6% |
| ***paniculata*** | | 56.3% | 83.8% | 72.3% | 59.0% | 85.0% | 84.1% | 57.2% | 83.4% | 72.0% | 75.8% |
| ***pauciflora*** | 61.8% | | 60.4% | 62.7% | 91.7% | 63.3% | 62.5% | 66.7% | 61.8% | 62.6% | 65.8% |
| ***polyanthemos*** | 83.9% | 53.6% | | 71.0% | 53.9% | 82.8% | 83.6% | 56.4% | 81.0% | 70.1% | 72.8% |
| ***pumila*** | 74.9% | 59.2% | 74.1% | | 57.9% | 74.6% | 73.1% | 60.3% | 72.7% | 82.0% | 75.3% |
| ***regnans*** | 64.6% | 90.9% | 60.7% | 61.4% | | 63.2% | 61.0% | 66.3% | 60.9% | 60.8% | 64.0% |
| ***shirleyi*** | 85.3% | 57.3% | 82.8% | 71.9% | 56.2% | | 83.3% | 57.9% | 82.1% | 71.3% | 74.3% |
| ***sideroxylon*** | 84.2% | 55.0% | 83.2% | 70.3% | 53.8% | 83.5% | | 55.8% | 81.0% | 69.7% | 73.3% |
| ***tenuipes*** | 70.4% | 72.5% | 69.0% | 70.1% | 72.5% | 70.6% | 69.1% | | 68.6% | 68.8% | 71.5% |
| ***victrix*** | 84.5% | 56.9% | 82.0% | 71.7% | 55.9% | 83.4% | 82.1% | 57.5% | | 71.3% | 74.0% |
| ***viminalis*** | 72.8% | 57.1% | 70.6% | 80.3% | 54.7% | 71.7% | 70.4% | 56.6% | 69.8% | | 71.9% |
| ***virginea*** | 77.0% | 60.6% | 74.7% | 74.7% | 59.1% | 76.2% | 74.7% | 61.5% | 74.2% | 72.5% | |

**Supplementary Tables S12. Matrix of pairwise shared sequence (synteny + rearranged).** Shows the proportion of the genome that is shared for all genome pairs. Read down the genome list (far left column) and extend across until the comparison genome is found. As genomes are different lengths the proportion that species X shares with species Y is different to the proportion that species Y shares with species X.

| | *albens* | *brandiana* | *caleyi* | *camaldulensis* | *cladocalyx* | *cloeziana* | *coolabah* | *curtisii* | *dawsonii* | *decipiens* | *erythrocorys* |
|---|---|---|---|---|---|---|---|---|---|---|---|
| ***albens*** | | 25.6% | 17.4% | 29.7% | 23.2% | 45.1% | 19.9% | 44.4% | 16.7% | 28.6% | 50.7% |
| ***brandiana*** | 20.1% | | 19.3% | 26.2% | 20.9% | 39.6% | 21.4% | 38.5% | 18.7% | 22.7% | 44.4% |
| ***caleyi*** | 17.5% | 25.3% | | 28.1% | 22.9% | 43.6% | 18.4% | 42.4% | 16.2% | 26.6% | 48.1% |
| ***camaldulensis*** | 28.6% | 31.0% | 28.1% | | 29.7% | 43.3% | 30.8% | 43.0% | 27.9% | 30.0% | 47.5% |
| ***cladocalyx*** | 19.5% | 22.9% | 19.5% | 27.7% | | 41.9% | 21.9% | 40.7% | 19.1% | 25.0% | 46.6% |
| ***cloeziana*** | 37.0% | 37.8% | 36.3% | 37.8% | 38.1% | | 37.6% | 32.0% | 37.4% | 36.5% | 42.2% |
| ***coolabah*** | 19.4% | 27.0% | 18.2% | 31.8% | 25.4% | 45.4% | | 45.3% | 18.9% | 29.7% | 50.1% |
| ***curtisii*** | 32.8% | 33.0% | 31.5% | 33.0% | 32.7% | 27.9% | 33.6% | | 33.3% | 31.8% | 37.6% |
| ***dawsonii*** | 19.4% | 27.3% | 19.3% | 31.7% | 25.8% | 46.8% | 22.2% | 47.6% | | 30.4% | 53.3% |
| ***decipiens*** | 27.7% | 28.2% | 27.6% | 30.7% | 28.9% | 43.3% | 30.0% | 42.6% | 27.7% | | 49.6% |
| ***erythrocorys*** | 44.6% | 44.7% | 44.2% | 45.1% | 45.5% | 44.6% | 45.8% | 45.0% | 47.0% | 45.5% | |
| ***fibrosa*** | 15.8% | 23.7% | 14.7% | 27.2% | 21.6% | 43.0% | 17.0% | 42.3% | 15.9% | 25.7% | 48.6% |
| ***globulus*** | 27.9% | 29.6% | 27.0% | 18.1% | 29.1% | 42.2% | 30.2% | 41.9% | 27.6% | 28.4% | 47.8% |
| ***grandis*** | 29.3% | 31.5% | 28.3% | 19.4% | 30.6% | 44.4% | 31.6% | 44.6% | 29.7% | 29.7% | 50.5% |
| ***guilfoylei*** | 29.6% | 28.7% | 28.6% | 29.8% | 29.9% | 32.4% | 30.6% | 31.3% | 29.7% | 28.0% | 38.3% |
| ***lansdowneana*** | 15.7% | 24.5% | 16.4% | 29.0% | 22.8% | 44.6% | 18.7% | 44.5% | 16.3% | 27.0% | 49.9% |
| ***leucophloia*** | 19.7% | 23.5% | 19.2% | 28.4% | 23.8% | 42.3% | 20.9% | 42.0% | 20.6% | 27.3% | 47.3% |
| ***marginata*** | 38.4% | 38.8% | 37.3% | 38.9% | 38.9% | 18.8% | 39.1% | 33.3% | 38.8% | 38.1% | 43.5% |
| ***melliodora*** | 16.6% | 26.2% | 18.3% | 30.8% | 25.1% | 46.6% | 20.7% | 45.4% | 17.3% | 29.1% | 50.9% |
| ***melliodora x sideroxylon*** | 36.5% | 36.5% | 36.5% | 36.5% | 36.5% | 36.5% | 36.5% | 36.5% | 36.5% | 36.5% | 36.5% |
| ***microcorys*** | 27.5% | 28.0% | 27.5% | 28.7% | 28.0% | 32.0% | 28.7% | 30.0% | 27.3% | 26.4% | 36.5% |
| ***ANBG9806169*** | 39.7% | 39.3% | 38.1% | 38.9% | 39.8% | 16.0% | 39.9% | 33.7% | 40.5% | 38.8% | 43.1% |
| ***paniculata*** | 15.9% | 23.3% | 14.4% | 27.2% | 20.9% | 41.8% | 17.7% | 41.8% | 15.7% | 26.3% | 48.9% |
| ***pauciflora*** | 37.4% | 38.8% | 36.6% | 37.3% | 37.6% | 13.3% | 38.4% | 32.2% | 38.0% | 36.7% | 42.5% |
| ***polyanthemos*** | 16.8% | 25.0% | 16.7% | 29.2% | 22.8% | 44.3% | 19.0% | 44.3% | 12.2% | 28.0% | 49.5% |
| ***pumila*** | 26.4% | 27.9% | 25.9% | 16.9% | 27.1% | 40.7% | 28.8% | 40.6% | 26.6% | 26.4% | 45.9% |
| ***regnans*** | 38.3% | 37.9% | 37.8% | 38.8% | 39.6% | 14.1% | 39.8% | 33.5% | 39.6% | 37.6% | 44.0% |
| ***shirleyi*** | 16.4% | 24.5% | 15.4% | 28.4% | 22.9% | 43.4% | 17.5% | 42.4% | 17.1% | 27.3% | 47.5% |
| ***sideroxylon*** | 16.0% | 25.2% | 17.0% | 29.0% | 23.9% | 44.3% | 19.1% | 44.3% | 16.1% | 27.8% | 49.9% |
| ***tenuipes*** | 30.0% | 31.3% | 29.3% | 30.6% | 31.1% | 25.4% | 31.7% | 21.6% | 31.5% | 30.1% | 34.3% |
| ***victrix*** | 17.3% | 24.5% | 16.5% | 28.3% | 22.6% | 42.9% | 15.7% | 42.5% | 16.9% | 26.7% | 48.0% |
| ***viminalis*** | 29.1% | 30.6% | 28.3% | 19.6% | 30.4% | 43.5% | 30.7% | 43.2% | 29.0% | 29.7% | 48.1% |
| ***virginea*** | 24.7% | 25.8% | 23.9% | 26.9% | 25.6% | 39.6% | 26.8% | 38.6% | 24.0% | 20.5% | 44.3% |

| | ***fibrosa*** | ***globulus*** | ***grandis*** | ***guilfoylei*** | ***lansdowneana*** | ***leucophloia*** | ***marginata*** | ***melliodora*** | ***melliodora x sideroxylon*** | ***microcorys*** | ***ANBG9806169*** |
|---|---|---|---|---|---|---|---|---|---|---|---|
| ***albens*** | 16.3% | 30.4% | 28.0% | 37.0% | 16.0% | 21.4% | 44.4% | 16.4% | 15.3% | 37.7% | 46.2% |
| ***brandiana*** | 18.8% | 27.0% | 24.9% | 30.7% | 19.2% | 19.1% | 38.7% | 20.6% | 18.4% | 32.6% | 39.4% |
| ***caleyi*** | 15.2% | 28.6% | 26.5% | 35.4% | 16.5% | 20.0% | 42.1% | 17.3% | 16.2% | 36.7% | 43.7% |
| ***camaldulensis*** | 26.8% | 19.4% | 16.5% | 35.4% | 28.3% | 28.6% | 42.4% | 29.4% | 26.7% | 36.8% | 42.9% |
| ***cladocalyx*** | 18.2% | 27.9% | 25.6% | 33.1% | 19.1% | 21.1% | 40.7% | 20.4% | 18.2% | 34.3% | 42.5% |
| ***cloeziana*** | 36.6% | 37.9% | 36.2% | 32.9% | 37.4% | 36.7% | 17.3% | 38.5% | 34.9% | 34.9% | 15.7% |
| ***coolabah*** | 17.0% | 32.0% | 29.6% | 37.7% | 18.3% | 21.8% | 44.3% | 19.4% | 17.6% | 38.4% | 45.5% |
| ***curtisii*** | 31.9% | 33.1% | 31.7% | 27.3% | 33.0% | 32.4% | 26.8% | 33.6% | 30.6% | 29.0% | 28.3% |
| ***dawsonii*** | 18.5% | 32.2% | 30.5% | 39.3% | 18.3% | 24.2% | 46.2% | 18.5% | 17.8% | 40.0% | 47.6% |
| ***decipiens*** | 25.6% | 30.0% | 28.2% | 34.7% | 26.7% | 28.4% | 42.8% | 27.8% | 26.3% | 35.7% | 43.3% |
| ***erythrocorys*** | 44.5% | 45.3% | 44.5% | 41.5% | 45.0% | 45.2% | 44.6% | 46.0% | 43.3% | 42.2% | 44.6% |
| ***fibrosa*** | | 28.0% | 25.5% | 33.4% | 16.2% | 18.9% | 41.9% | 16.0% | 14.7% | 34.9% | 43.9% |
| ***globulus*** | 26.4% | | 15.0% | 33.8% | 26.9% | 28.8% | 41.8% | 28.8% | 25.3% | 35.3% | 42.4% |
| ***grandis*** | 28.1% | 19.3% | | 36.6% | 28.8% | 29.9% | 43.4% | 29.9% | 27.1% | 36.7% | 44.2% |
| ***guilfoylei*** | 27.9% | 29.6% | 28.3% | | 28.9% | 28.5% | 31.7% | 29.6% | 27.3% | 25.9% | 34.0% |
| ***lansdowneana*** | 16.3% | 29.6% | 27.6% | 36.2% | | 20.5% | 43.5% | 15.7% | 14.3% | 36.6% | 45.2% |
| ***leucophloia*** | 18.2% | 29.6% | 27.1% | 34.2% | 19.8% | | 41.8% | 20.6% | 18.9% | 35.2% | 42.5% |
| ***marginata*** | 37.6% | 39.3% | 37.2% | 33.3% | 38.2% | 38.1% | | 39.2% | 35.9% | 36.1% | 16.9% |
| ***melliodora*** | 17.6% | 31.0% | 28.8% | 37.8% | 17.3% | 23.3% | 45.2% | | 24.5% | 24.5% | 24.5% |
| ***melliodora x sideroxylon*** | 36.5% | 36.5% | 36.5% | 36.5% | 36.5% | 36.5% | 36.5% | 36.5% | | 36.5% | 36.5% |
| ***microcorys*** | 26.3% | 28.5% | 26.2% | 23.1% | 27.0% | 26.6% | 31.4% | 29.1% | 25.9% | | 31.9% |
| ***ANBG9806169*** | 38.8% | 39.5% | 37.3% | 35.8% | 39.6% | 38.8% | 16.1% | 40.1% | 36.9% | 36.5% | |
| ***paniculata*** | 13.8% | 27.9% | 26.2% | 34.6% | 16.0% | 20.6% | 41.2% | 16.1% | 14.5% | 34.8% | 43.1% |
| ***pauciflora*** | 36.7% | 37.6% | 36.7% | 33.4% | 37.6% | 38.3% | 13.8% | 39.6% | 36.7% | 34.5% | 8.1% |
| ***polyanthemos*** | 16.0% | 29.8% | 27.8% | 36.4% | 16.6% | 21.9% | 43.7% | 16.5% | 15.6% | 37.4% | 44.5% |
| ***pumila*** | 25.5% | 18.4% | 16.4% | 31.8% | 25.7% | 27.0% | 40.1% | 27.1% | 24.1% | 33.4% | 41.9% |
| ***regnans*** | 37.6% | 39.4% | 37.7% | 33.1% | 38.3% | 38.2% | 14.5% | 40.1% | 36.6% | 35.3% | 8.7% |
| ***shirleyi*** | 15.3% | 29.1% | 26.9% | 35.3% | 16.9% | 20.3% | 42.2% | 16.7% | 15.8% | 34.9% | 43.3% |
| ***sideroxylon*** | 15.9% | 30.4% | 28.4% | 36.2% | 16.0% | 21.5% | 43.3% | 15.2% | 11.8% | 37.6% | 44.4% |
| ***tenuipes*** | 29.2% | 30.7% | 29.2% | 24.9% | 30.5% | 29.8% | 25.2% | 32.4% | 29.7% | 27.3% | 26.5% |
| ***victrix*** | 16.5% | 28.7% | 26.8% | 34.4% | 17.3% | 19.3% | 42.1% | 17.8% | 16.3% | 36.0% | 42.8% |
| ***viminalis*** | 26.7% | 14.5% | 16.3% | 35.3% | 28.6% | 29.9% | 43.0% | 30.0% | 26.7% | 36.7% | 43.3% |
| ***virginea*** | 23.3% | 26.9% | 24.9% | 31.1% | 24.1% | 24.9% | 38.7% | 25.5% | 22.6% | 31.7% | 39.6% |

| | paniculata | pauciflora | polyanthemos | pumila | regnans | shirleyi | sideroxylon | tenuipes | victrix | viminalis | virginea |
|---|---|---|---|---|---|---|---|---|---|---|---|
| **albens** | 16.0% | 43.5% | 17.0% | 30.1% | 46.6% | 17.2% | 16.8% | 44.5% | 19.6% | 30.7% | 27.4% |
| **brandiana** | 17.9% | 41.2% | 20.3% | 25.7% | 40.6% | 19.0% | 20.6% | 39.4% | 21.4% | 26.4% | 22.3% |
| **caleyi** | 14.0% | 43.0% | 17.3% | 28.3% | 45.2% | 15.3% | 17.4% | 42.3% | 18.5% | 29.3% | 25.9% |
| **camaldulensis** | 27.0% | 43.3% | 28.8% | 19.0% | 44.8% | 27.3% | 29.2% | 41.6% | 29.5% | 19.7% | 27.9% |
| **cladocalyx** | 18.2% | 42.3% | 19.5% | 26.8% | 43.5% | 19.4% | 20.7% | 41.6% | 21.4% | 28.2% | 24.1% |
| **cloeziana** | 35.9% | 15.7% | 37.4% | 37.6% | 15.5% | 36.8% | 37.4% | 32.3% | 38.0% | 37.7% | 35.1% |
| **coolabah** | 17.5% | 44.8% | 19.1% | 31.8% | 47.1% | 17.2% | 19.8% | 46.0% | 17.6% | 32.3% | 28.9% |
| **curtisii** | 31.2% | 27.9% | 33.2% | 32.7% | 28.9% | 31.5% | 33.1% | 23.9% | 33.2% | 33.4% | 30.5% |
| **dawsonii** | 18.7% | 46.3% | 14.5% | 32.4% | 48.9% | 19.8% | 18.7% | 47.1% | 21.7% | 33.0% | 29.3% |
| **decipiens** | 26.8% | 43.9% | 28.5% | 29.1% | 44.6% | 27.8% | 28.3% | 43.8% | 28.8% | 29.9% | 21.8% |
| **erythrocorys** | 45.0% | 45.1% | 45.5% | 45.5% | 47.3% | 45.5% | 45.7% | 43.7% | 45.7% | 46.4% | 43.7% |
| **fibrosa** | 13.5% | 43.0% | 15.8% | 27.5% | 44.6% | 15.1% | 16.2% | 42.3% | 18.2% | 27.6% | 24.8% |
| **globulus** | 25.9% | 41.7% | 28.0% | 18.5% | 44.0% | 26.9% | 28.6% | 42.3% | 28.7% | 13.5% | 26.6% |
| **grandis** | 27.7% | 44.9% | 29.7% | 20.7% | 46.4% | 28.1% | 29.8% | 44.3% | 29.8% | 18.9% | 28.2% |
| **guilfoylei** | 27.6% | 33.3% | 29.3% | 28.7% | 33.4% | 28.6% | 29.8% | 31.7% | 29.6% | 30.1% | 27.5% |
| **lansdowneana** | 15.6% | 44.5% | 16.4% | 28.9% | 46.0% | 16.3% | 15.7% | 44.1% | 18.9% | 29.5% | 26.0% |
| **leucophloia** | 19.2% | 42.9% | 20.6% | 28.6% | 43.8% | 18.5% | 21.0% | 40.8% | 19.6% | 29.5% | 25.5% |
| **marginata** | 36.4% | 16.7% | 38.5% | 38.3% | 17.2% | 37.5% | 38.1% | 33.7% | 39.2% | 39.0% | 36.5% |
| **melliodora** | 24.5% | 24.5% | 24.5% | 24.5% | 24.5% | 24.5% | 24.5% | 24.5% | 24.5% | 24.5% | 24.5% |
| **melliodora x sideroxylon** | 36.5% | 36.5% | 36.5% | 36.5% | 36.5% | 36.5% | 36.5% | 36.5% | 36.5% | 36.5% | 36.5% |
| **microcorys** | 25.4% | 32.6% | 27.7% | 27.2% | 32.8% | 26.3% | 28.2% | 30.6% | 28.3% | 28.8% | 25.4% |
| **ANBG9806169** | 38.5% | 11.1% | 39.6% | 40.0% | 10.7% | 38.7% | 39.4% | 33.7% | 39.8% | 39.6% | 37.4% |
| **paniculata** | | 43.7% | 16.2% | 27.7% | 41.0% | 15.0% | 15.9% | 42.8% | 16.6% | 28.0% | 24.2% |
| **pauciflora** | 38.2% | | 39.6% | 37.3% | 8.3% | 36.7% | 37.5% | 33.3% | 38.2% | 37.4% | 34.2% |
| **polyanthemos** | 16.1% | 46.4% | | 29.0% | 46.1% | 17.2% | 16.4% | 43.6% | 19.0% | 29.9% | 27.2% |
| **pumila** | 25.1% | 40.8% | 25.9% | | 42.1% | 25.4% | 26.9% | 39.7% | 27.3% | 18.0% | 24.7% |
| **regnans** | 35.4% | 9.1% | 39.3% | 38.6% | | 36.8% | 39.0% | 33.7% | 39.1% | 39.2% | 36.0% |
| **shirleyi** | 14.7% | 42.7% | 17.2% | 28.1% | 43.8% | | 16.7% | 42.1% | 17.9% | 28.7% | 25.7% |
| **sideroxylon** | 15.8% | 45.0% | 16.8% | 29.7% | 46.2% | 16.5% | | 44.2% | 19.0% | 30.3% | 26.7% |
| **tenuipes** | 29.6% | 27.5% | 31.0% | 29.9% | 27.5% | 29.4% | 30.9% | | 31.4% | 31.2% | 28.5% |
| **victrix** | 15.5% | 43.1% | 18.0% | 28.3% | 44.1% | 16.6% | 17.9% | 42.5% | | 28.7% | 26.0% |
| **viminalis** | 27.2% | 42.9% | 29.4% | 19.7% | 45.3% | 28.3% | 29.6% | 43.4% | 30.2% | | 28.1% |
| **virginea** | 23.0% | 39.4% | 25.3% | 25.3% | 40.9% | 23.8% | 25.3% | 38.5% | 25.8% | 27.5% | |

**Supplementary Tables S13. Matrix of pairwise unaligned.** Shows the proportion of the genome that is unaligned for all genome pairs. Read down the genome list (far left column) and extend across until the comparison genome is found.

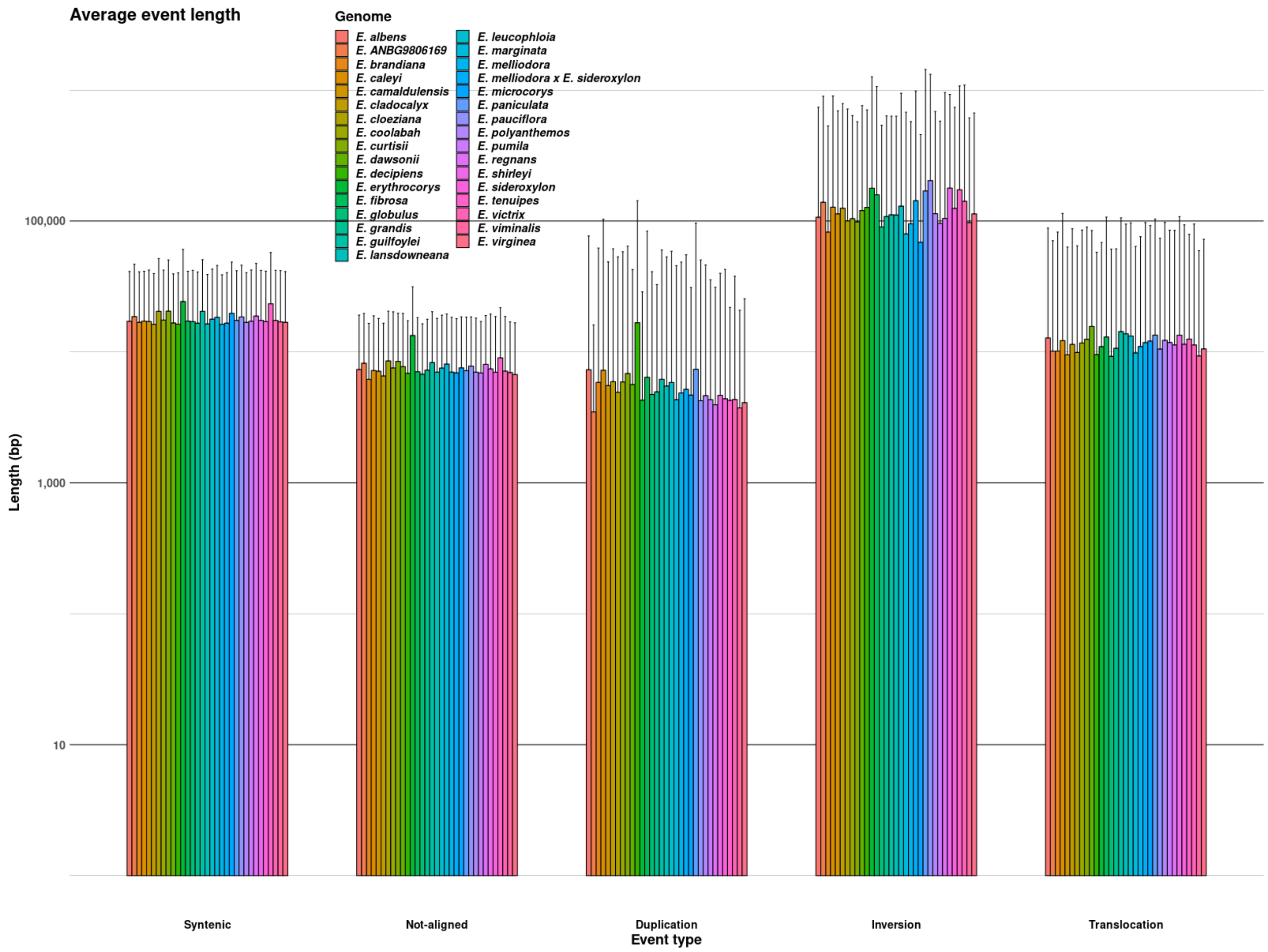


**Supplementary Figure S18.** Average length of syntenic, unaligned, rearrangement (combined duplications, inversions, and translocations), duplicated, inverted, and translocated regions, for all comparisons within each genome. Error bars show standard deviation of region lengths.

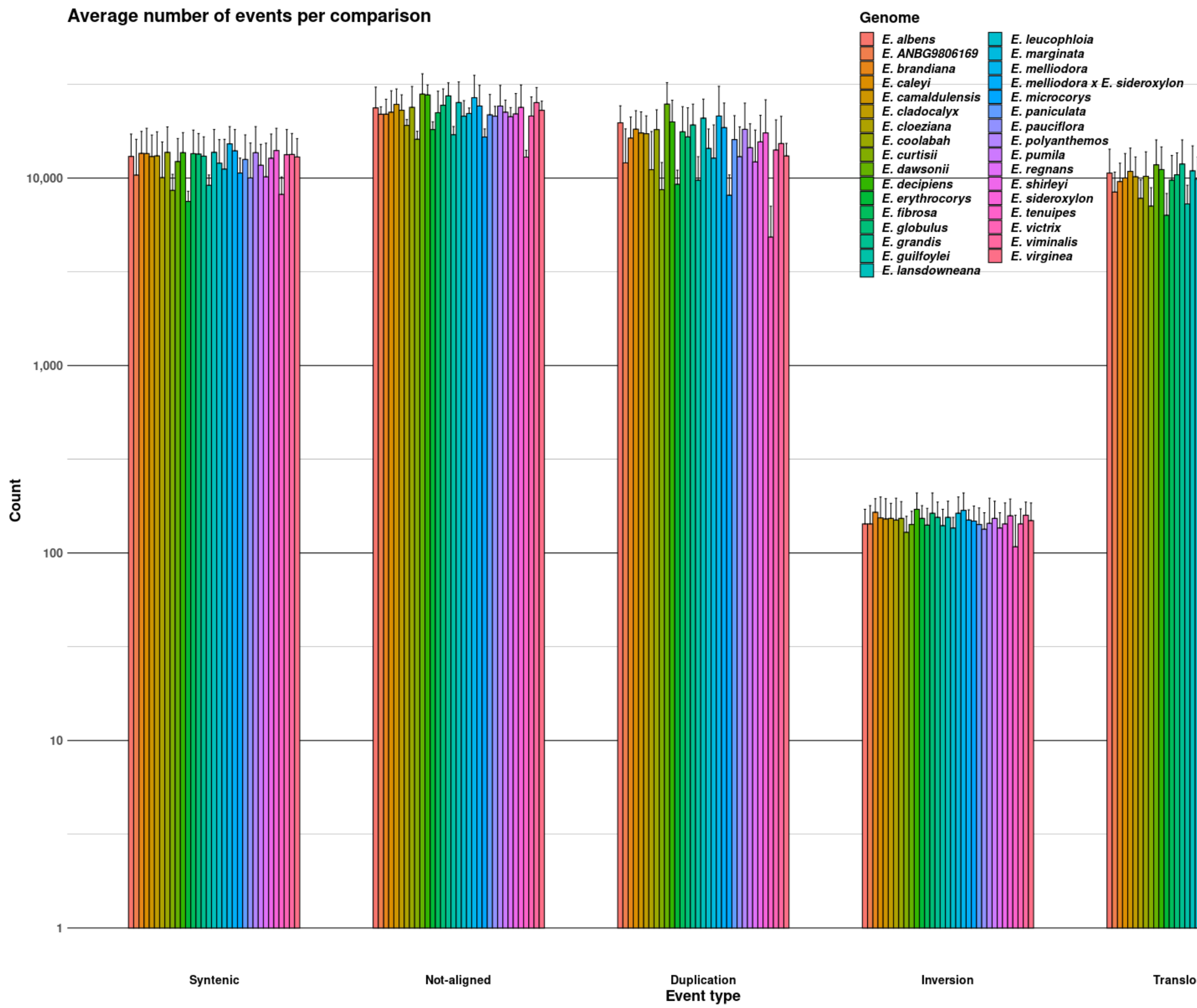


**Supplementary Figure S19.** Average number of syntenic, unaligned, rearrangement (combined duplications, inversions, and translocations), duplicated, inverted, and translocated regions, for all comparisons within each genome. Error bars show the minimum and maximum number of events counted within a comparison for each genome

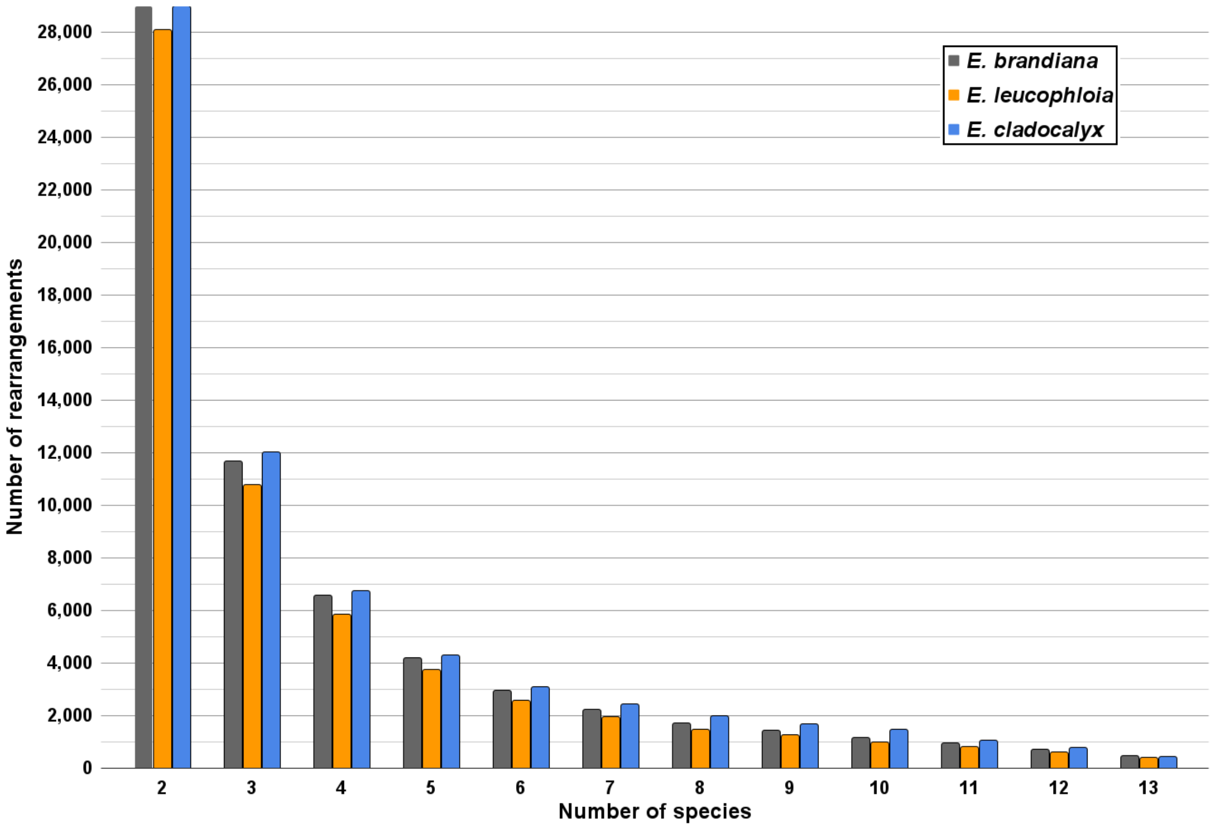


**Supplementary Figure S20.** Ignoring the phylogenetic relationships between genomes, shows the number of rearrangements shared by an increasing number of *Adnataria* genomes using *E. brandiana*, *E. cladocalyx*, and *E. leucophylla* as the outgroup/genetic architecture.

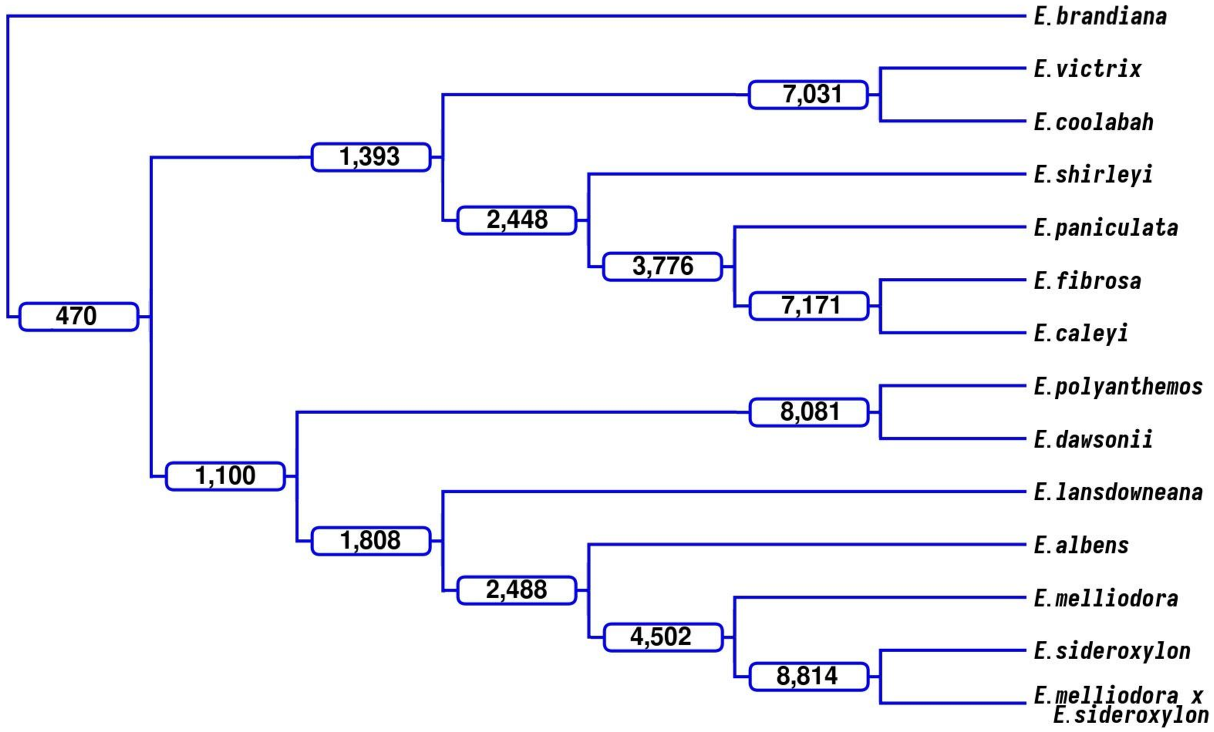


**Supplementary Figures S21.** Repeat of Adnataria phylogeny of common rearrangements. *E. brandiana* is the out genome

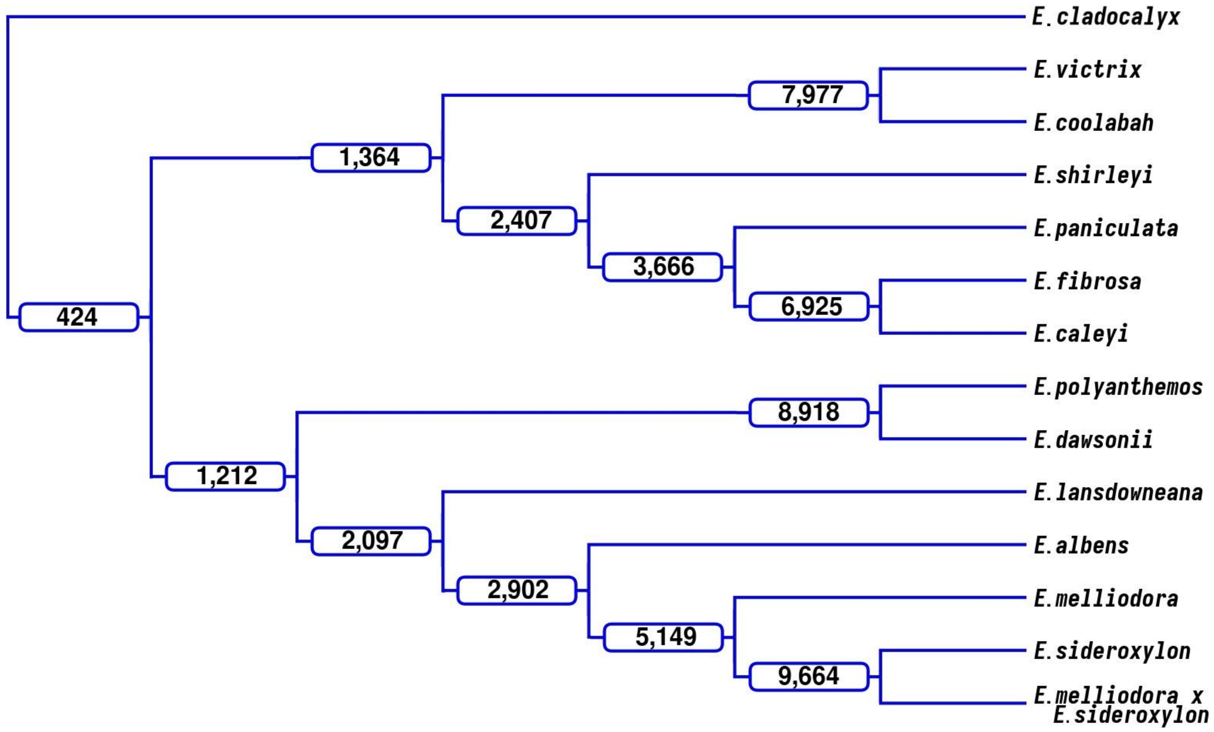


**Supplementary Figures S22.** Repeat of Adnataria phylogeny of common rearrangements. *E. cladocalyx* is the out genome.

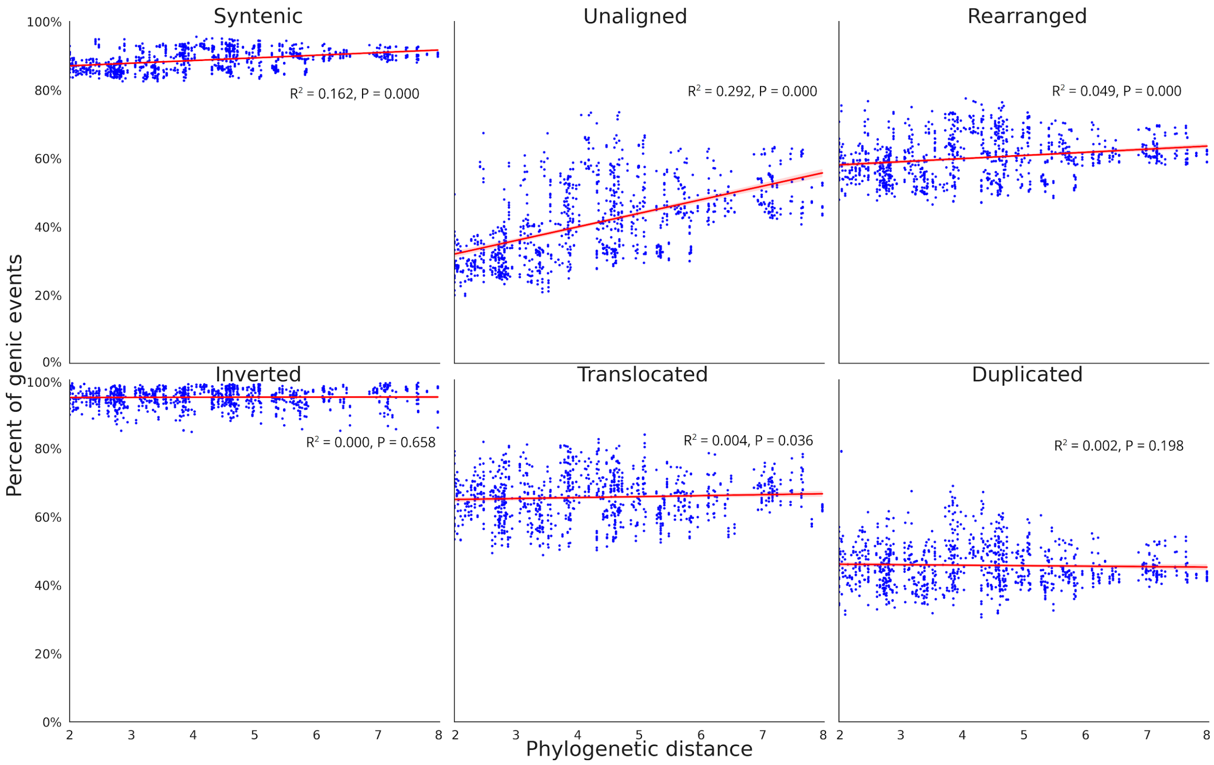


**Supplementary Figures S23. Phylogenetic distance and the number of events syntenic, rearrangement, and unaligned events.** Pairwise genome conservation and loss, as phylogenetic distance increases. The number of events between both Eucalyptus genomes with an alignment pair that was identified as syntenic, rearranged, or unaligned, plotted against the phylogenetic distance of the two genomes. The unaligned proportion is the species-specific fraction of the genome between genome pairs, resulting from either an insertion, deletion, differential inheritance, or sequence divergence. Rearranged events are broken down into inverted, translocated, and duplicated regions. Phylogenetic distance was calculated as the sum of branch lengths between each genome pair within phylogeny.
